# Stimulus-responsive Self-Assembly of Protein-Based Fractals by Computational Design

**DOI:** 10.1101/274183

**Authors:** Nancy E. Hernández, William A. Hansen, Denzel Zhu, Maria E. Shea, Marium Khalid, Viacheslav Manichev, Matthew Putnins, Muyuan Chen, Anthony G. Dodge, Lu Yang, Ileana Marrero-Berríos, Melissa Banal, Phillip Rechani, Torgny Gustafsson, Leonard C. Feldman, Sang-Hyuk Lee, Lawrence P. Wackett, Wei Dai, Sagar D. Khare

## Abstract

Fractal topologies, which are statistically self-similar over multiple length scales, are pervasive in nature. The recurrence of patterns at increasing length scales in fractal-shaped branched objects, *e.g.*, trees, lungs, and sponges, results in high effective surface areas, and provides key functional advantages, *e.g.*, for molecular trapping and exchange. Mimicking these topologies in designed protein-based assemblies will provide access to novel classes of functional biomaterials for wide ranging applications. Here we describe a computational design approach for the reversible self-assembly of proteins into tunable supramolecular fractal-like topologies in response to phosphorylation. Computationally-guided atomic-resolution modeling of fusions of symmetric, oligomeric proteins with Src homology 2 (SH2) binding domain and its phosphorylatable ligand peptide was used to design iterative branching leading to assembly formation by two enzymes of the atrazine degradation pathway. Structural characterization using various microscopy techniques and Cryo-electron tomography revealed a variety of dendritic, hyperbranched, and sponge-like topologies which are self-similar over three decades (~10nm-10μm) of length scale, in agreement with models from multi-scale computational simulations. Control over assembly topology and formation dynamics is demonstrated. Owing to their sponge-like structure on the nanoscale, fractal assemblies are capable of efficient and phosphorylation-dependent reversible macromolecular capture. The described design framework should enable the construction of a variety of novel, spatiotemporally responsive biomaterials featuring fractal topologies.

**One Sentence Summary:** We report a computationally-guided bottom up design approach for constructing fractal-shaped protein assemblies for efficient molecular capture.

## Main Text

Fractional-dimensional (fractal) geometry – a property of shapes that are invariant or nearly invariant to scale magnification or contraction across many length scales – is a common feature of many natural objects^1,2^. Fractal forms are ubiquitous in geology, *e.g.*, in the architecture of mountain ranges, coastlines, snowflakes, and in physiology, *e.g.,* neuronal and capillary networks, and nasal membranes, where highly efficient molecular exchange occurs due to a fractal-induced high surface area:volume ratio^3^. Fabrication of fractal-like nanomaterials affords high physical connectivity within patterned objects^4^, ultrasensitive detection of target binding moieties by patterned nanosensors^5^, and rapid exchange and dispersal of energy and matter^6^. An intimate link between structural fractal properties of designed, nanotextured materials and functional advantages (e.g., detection sensitivity) has been demonstrated^5^, and synthetic fractal materials are finding applications in sensing, molecular electronics, high-performance filtration, sunlight collection, surface charge storage, and catalysis, among myriad other uses^7,8^. Many fractal fabrication efforts have relied on top-down patterning of surfaces^9^. The bottom-up design of supramolecular fractal topologies – both deterministic (e.g., Sierpinski’s triangles)^10,11^ and stochastic fractals (e.g., arborols)^12,13^– has been performed with small molecule building blocks such as inorganic metal-ligand complexes or synthetic dendritic polymers utilizing co-ordinate or covalent bonds, respectively. Self-similar quasi-fractal shapes built with DNA origami have been reported^14–16^, however, fractal topologies have not been designed with proteins, which possess a wide range of functionality, biocompatibility, and whose properties are dynamically controllable by reversible post-translational modifications^17^. While fractal-like topologies have been detected as intermediates in the formation of natural protein-based biomaterials such as biosilica and silk^18,19^, and observed in peptide assemblies^20–22^, their tunable construction by utilizing reversible non-covalent interactions between protein building blocks under mild conditions remains a fundamental design challenge.

Self-assembly of engineered proteins^23^ provides a general framework for the controllable and bottom-up fabrication of novel biomaterials with chosen supramolecular topologies but these approaches have, thus far, been applied to the design of integer (two or three)-dimensional ordered patterns such as layers, lattices, and polyhedra^24–30^. While external triggers such as metal ions and redox conditions have been used to trigger synthetic protein and peptide assemblies^20,21,31–34^, phosphorylation – a common biological stimulus used for dynamic control over protein function – has yet to be utilized for controlling protein assembly formation.

Among stochastic fractals, an arboreal (tree-like) shape is an elementary topology that can be generated using stochastic branching algorithms, *e.g.*, L-systems^35,36^, in which the probability of branching, length and number of branches, and branching angle ranges at each iteration determine the emergent topology (Fig. 1A). Theoretical and simulation studies on the self-assembly of ‘patchy’ colloidal particles^37,38^ have shown that a variety of topologies, including fractal-like topologies^39–41^, can result from stochastic self-assembly processes involving strong, anisotropic short-range forces^42–46^. Under conditions where inter-molecular interaction energy is much larger (more negative) than thermal energy, emergent large-scale aggregates are expected to be out-of-equilibrium kinetically trapped states rather than (usually crystalline) globally stable thermodynamic minima. As reorganization of aggregate morphologies, once formed, is expected to be unfavorable, we reasoned that these kinetic traps can be utilized to produce an elaborate, tunable and responsive structural (and, thus, functional) diversity of self-assembled protein-based systems^47^.

**Figure 1:**
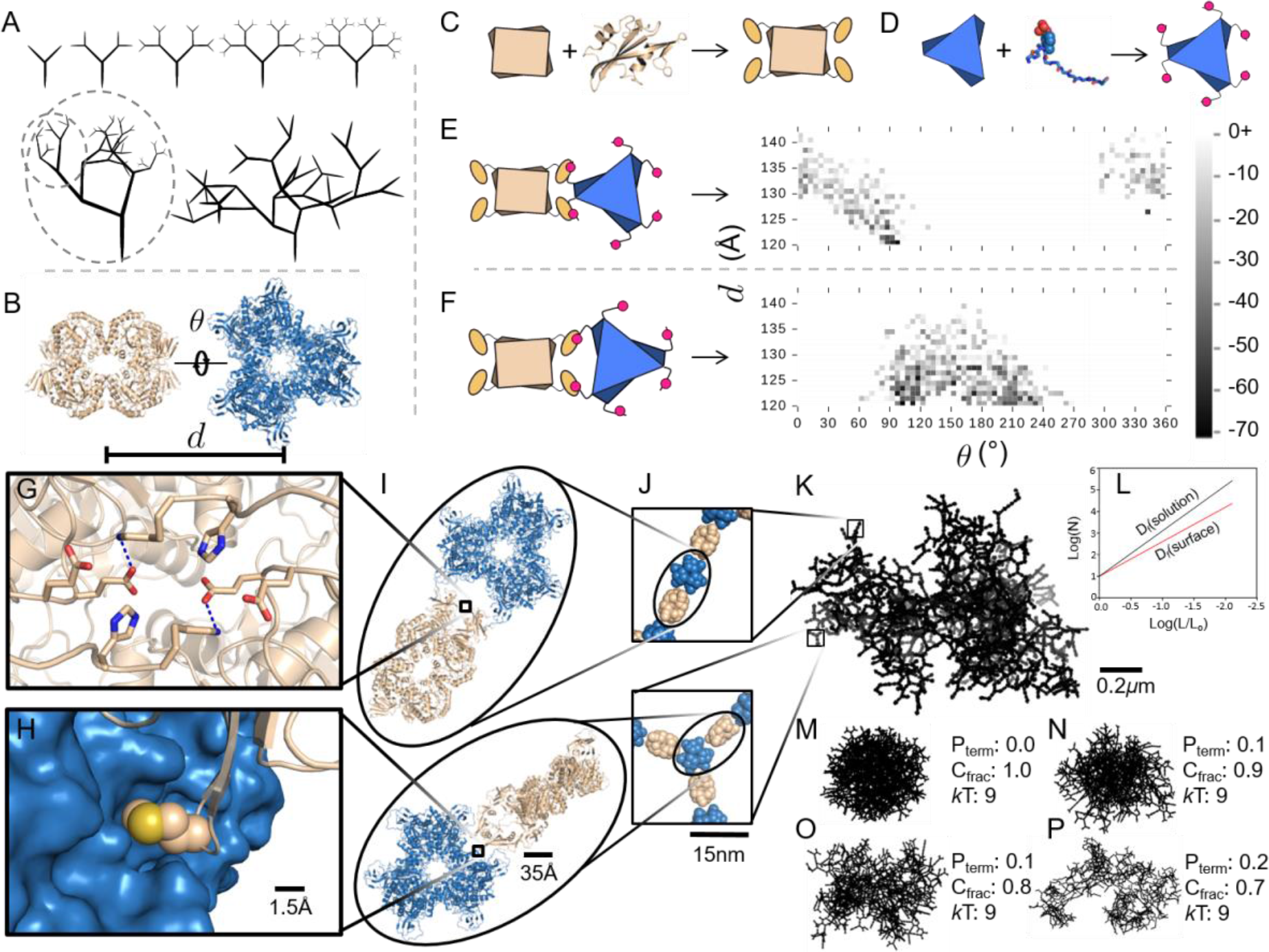
Multi-scale Computational Design Approach for fractal assembly design with pY-AtzA and AtzC-SH2. (**A**) Cartoon representations of an ordered self-similar scaling fractal, an unordered self-similar scaling fractal—note concentric circles are self-similar at different scales—and an unordered statistically self-similar fractal. (**B**) Two-component library of AtzC (tan) and AtzA (blue) positions was generated by varying the rigid body degrees of freedom along paired *C*_2_ symmetry axes. (**C and D**) Design and modeling of assembly at the molecular scale was performed by fusing an SH2 binding domain and its corresponding phosphorylatable peptide to AtzC (C) and AtzA (D) respectively. Linker between the SH2 domain and AtzC was designed to ensure symmetric binding between the hexamer and tetramer leading to propagation. (**E** and **F**) Flexibility analysis was performed by evaluation of the Rosetta energy landscape of symmetrical connections and the probability of observing different connection distances and angles were calculated using the Boltzmann distribution for two binding modes: vertex (E) and Edge (F). Boltzmann weighted connection probabilities were utilized in a stochastic chain-growth program with a coarse-grained protein model to generate emergent structures. Atomic interactions that stabilized novel interfaces formed from physically connected components (**G and H**) dictate the rotation along the C-symmetric axis between components (**I and J**) which ultimately produce combinations of orientations that lead to fractal-like topologies (**K**) on the micrometer scale. (**L**) Representation of expected fractal dimension (slope) for fractals analyzed in solution and on surfaces. (**M** to **P**) Examples of fractal simulation output across varying termination probabilities (P_term_) and fraction of components (C_frac_) at fixed *k*T.

To implement a general approach for tunably designing arboreal fractal morphologies using triggerable self-assembly of protein building blocks, we envisioned the need for: (a) a set of branching components whose binding to each other would lead to propagation of the assembly (Fig. 1A), (b) a modular system for connecting, with high affinity, these components reversibly in response to a chosen chemical trigger, and (c) degeneracy of protein-protein binding modes (geometries), such that stochastic but anisotropic, directional propagation of multiple branching geometries leads to emergent fractal-like supramolecular topologies (Fig. S1). We chose (a) the oligomeric enzymes AtzA (hexameric) and AtzC (tetrameric) of the atrazine biodegradation pathway^48^ featuring dihedral (*D*_3_ and *D*_2_, respectively) symmetry (Fig. 1B), (b) a phosphorylatable peptide (pY) tag with its corresponding engineered high-affinity “superbinder” Src homology 2 (SH2) domain^49^, and (c) linker segments that can stabilize multiple binding orientations, respectively, as design elements encoding these properties (Fig. 1B,C,D). We have previously utilized a similar binding domain-peptide fusion strategy to design non-propagating multi-component enzyme complexes^50^.

The sequences of the designed protein components were obtained using a procedure implemented in the Rosetta macromolecular modeling program^51^ aimed at making a maximum of three divalent connections between each AtzA and AtzC mediated by SH2 domain-phosphopeptide binding (Fig. 1C,D). Divalent connections between components were sought to enable avidity leading to strong, directional, short-range interactions (“aeolotropic interactions”^44^) that would promote fractal growth (Fig. 1E,F). We also reasoned that geometric degeneracy in the form of multiple propagatable (but still anisotropic) binding modes would favor fractal structures (Fig. S1). In the first step of the design procedure, one of the *C*_2_ axes of the crystallographic structures of the two components were aligned. (Fig. 1B). Two alignments (Fig. 1E,F), obtained by rotating AtzA (hexamer) by 180° about its *C*_3_ axis, were considered, and the remaining two symmetry-compatible degrees of freedom for placement – the inter-component center-of-mass distance *d* and rotation angle *θ* about the aligned axis of symmetry – were sampled (Fig. 1B,E,F). For every value of *d* we sampled several discrete values of *θ* that, if uniformly adopted, were predicted to lead to an infinitely propagatable integer-dimensional lattice (Fig. S1). The resulting propagatable placements were evaluated using RosettaMatch^31^ for geometrically feasible fusion to the SH2 domain and phosphopeptide with the C-terminal AtzC and N-terminal of AtzA, respectively (Fig. S2A,B). Loop closure of successful SH2 domain and phosphopeptide placements was performed using Rosetta Kinematic Loop Closure (Fig. S2C). Next, optimization of the new intra- and inter-component interfaces was performed using RosettaDesign (SI 1.3, Fig. S3). Five AtzA-AtzC fusion protein pairs were chosen for experimental characterization based on removal of steric clashes (as reflected by calculated Rosetta energy, Table S1), tight interface packing between SH2 domain and AtzC, and visual examination of design models (Fig. S3). We found that short, flexible linker sequences (eg. Gly-Gly-Ser) between the SH2 domain and AtzC led to the most efficient interface packing in designs (Fig. S3) while still potentially allowing multiple binding modes: several mutations were common among design models obtained at different (single) values of (*d, θ*) suggesting geometric degeneracy in binding by each variant (Table S1) would be feasible. Indeed, several other values from the propagatable angle set are energetically feasible for each designed AtzC-SH2 variant (Table S1).

To fully evaluate the predicted geometric degeneracy and anisotropy of binding in designed inter-component interactions, the conformational landscape over all (*d*, *θ)* pairs (Fig. 1E,F) was constructed using Rosetta SymmetricFastRelax simulations for a designed hexamer-tetramer complex, and the calculated energies (Figs. 1E,F) were Boltzmann-weighted (using a simulation temperature parameter, T) to obtain a probability distribution ***P***(*d*, *θ*) for branching geometry. This distribution, in turn, was used as input for a coarse-grained stochastic chain-growth tree generation algorithm for predicting ensembles of emergent topologies on the micrometer length scale (SI 1.6). Similar hierarchical approaches have previously been developed for modeling protein crystallization^52^, and colloidal particle^43^ and protein self-assembly^45^. In our approach, preferred inter-component interaction modes at the sub-nanometer scale (Fig. 1G,H) guide the emergence of higher order structures on the nanometer (Fig. 1I,J) and micrometer length scales (Fig. 1K). For comparison with experiments, ~100s of emergent structures in the resulting ensemble were analyzed to determine fractal dimension (*D*_F_) using the box counting image processing technique (Fig. 1L). The fractal dimension of an object is a measure of how its mass or shape scales as a function of length scale (Fig. S4): an object is considered fractal if this scaling exponent is non-integer and typically less than the Euclidean dimension in which the object is placed on a set of length scales (SI 5.4). For example, the *D*_F_ of vasculature patterns on the two-dimensional surface of human retina^53^ is 1.7, and a diffusion-limited aggregation cluster in three-dimensional space has a *D*_F_ is 2.3^54^. In our simulations, a variety of assembly sizes and fractal dimensions, *D*_F_, could be obtained by varying three parameters: (1) the fraction of growth sites selected at each growing layer allowed to continue propagation (*c*_frac_) which reflects the stoichiometry of the two components, (2) the probability of termination at any chosen propagatable branching point (P_term_), which reflects the affinity of interactions (lower the affinity, higher the the P_term_), and (3) the Boltzmann factor (*k*_B_T), which determines the sampling of inter-component conformational diversity calculated from Rosetta simulations (Fig 1M-P, Fig. S5, S6).

Genes encoding the designed AtzA and AtzC variants and the corresponding fusions of wild type domains were constructed and cloned into an *E. coli* BL21(DE3) strain harboring a second plasmid for the inducible expression of GroEL/ES chaperones to aid protein yields. Purified AtzA designs were each phosphorylated using Src kinase and the presence of phosphotyrosine was confirmed using ELISA assays (Fig. S7); binding and assembly formation with purified AtzC-SH2 domain fusions was assessed using Biolayer Interferometry and Dynamic Light Scattering, respectively. Phosphorylation, binding and complete conversion of monomers into 1-10 μm-sized particles upon mixing was best detected with the proteins pY-AtzAM1 and AtzCM1 (Fig. S8, S9, S10). Either phosphorylation levels were lower (Fig. S9A) or inter-component binding was weaker (Fig. S9D) with other designs, therefore, we chose pY-AtzAM1:AtzCM1 design pair for further characterization of assembly-disassembly processes (Fig. 2A). Apart from fusion of pY-tag and SH2 domain, these proteins feature 2 and 3 substitutions compared to their wild type parent, respectively (Table S1; Fig. S11 and S12).

**Figure 2:**
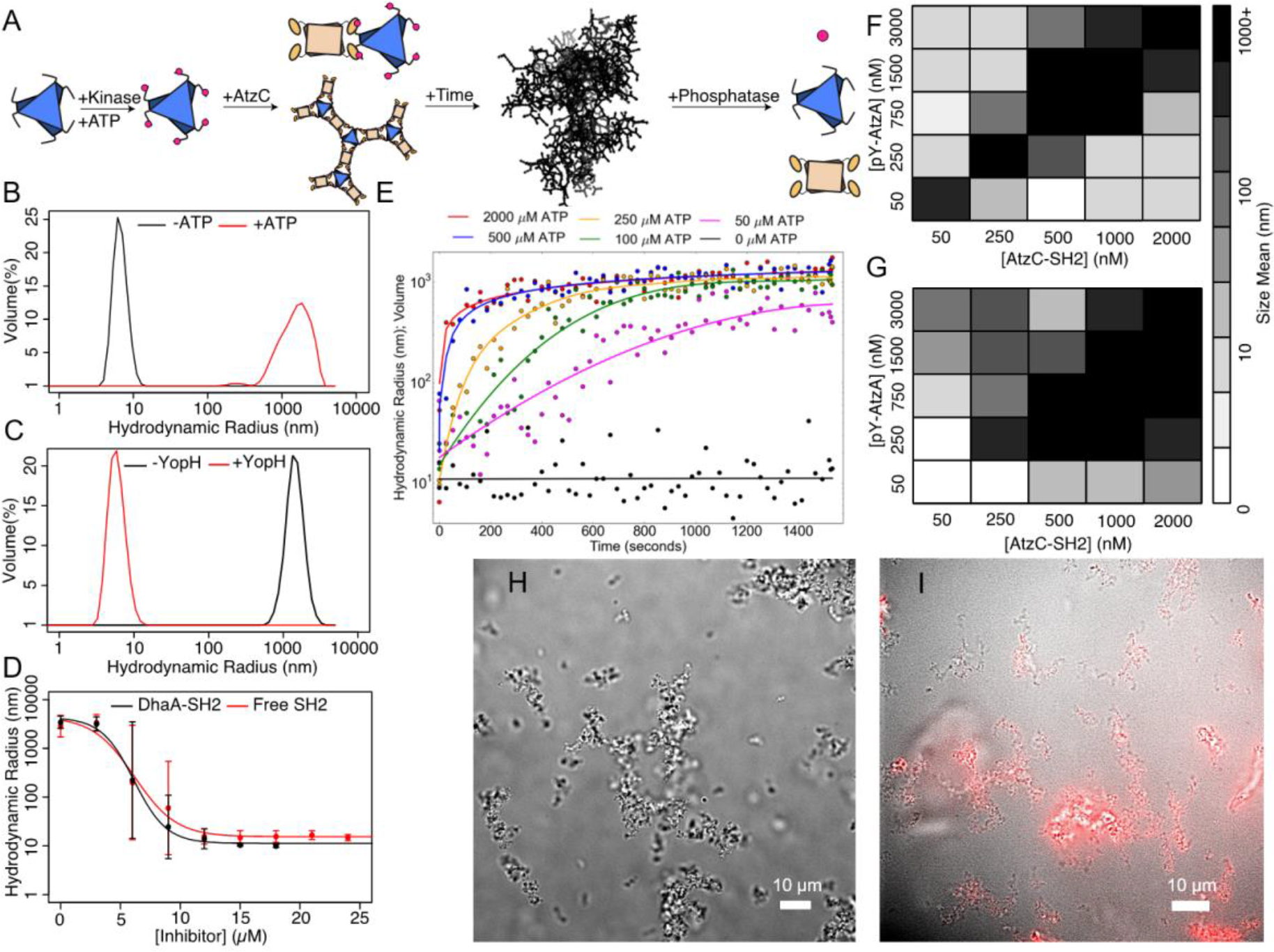
Assembly Formation, Dissolution and Inhibition *in vitro*. (**A**) Using Src kinase, the AtzAM1 can be phosphorylated (pY-AtzAM1) and incubated with AtzCM1-SH2 to form an assembly. The phosphatase (YOP) enzyme can be used for disassembly. (**B and C**) Assemblies were expected to form (B) and dissolve (C), respectively, as confirmed by DLS measurements. (**D**) Incubation of assembling components with various concentrations of free SH2 domain and a different (monovalent) SH2 fusion protein led to robust inhibition. (**E**) ATP concentration was shown to control the rate of assembly formation (highest concentration of ATP to lowest, starting from top to bottom at time 0) (**F and G**) Assembly formation is highly sensitive to stoichiometry of the components. Varying the stoichiometry (F and G) and the use of a weaker-binding SH2-peptide interaction (F) leads to a perturbation of the assembly formation zone compared to the “superbinder” SH2 (G). (**H**) The fractal-like structure observed by light microscopy. (**I**) Fluorescence microscopy image of assembly formed by Alexa Fluor 647^TM^-labeled AtzCM1-SH2 and pY-AtzAM1.

Assembly formation by a mixture of the two components and Src kinase enzyme was ATP dependent (Fig. 2B), was accompanied by the visible and spectrophotometrically measurable (Fig. S13) appearance of turbidity, which could be reversed by adding a phosphatase (YopH) enzyme. The resulting distribution of particle sizes was detected by measuring hydrodynamic radii using Dynamic Light Scattering (DLS) (Fig. 2C). Upon completion of assembly formation, the apparent size of the particles as measured by DLS was between 1-10 μ m; however, this range represents the upper limit of measurement for the instrument; actual particle sizes were expected to be larger. Addition of monovalent competitive inhibitors, *i.e.* isolated SH2 domain or SH2 domain fused to an unrelated monovalent protein (SH2-DhaA) inhibited assembly formation in a concentration-dependent manner, demonstrating that the SH2-pYtag binding interaction underlies assembly formation. The apparent IC_50_ for the observed inhibition was ~2×[AtzA-pY] (measured as monomers) at two different concentrations of the components (Fig. 2D, S14 to S16), and in each case ~3×[AtzA-pY] was required for complete inhibition. According to our design model, each pY-AtzA (hexamer) makes at least two and at most three divalent connections for assembly propagation (Fig. 1E,F); thus, the observed inhibition stoichiometries are consistent with the existence of the designed divalent connections between AtzA-pY and AtzC-SH2 in the assemblies.

As the phosphorylation reaction requires ATP, assembly formation rates could be controlled by varying the concentration of added ATP. For [AtzA-pY] and [AtzC-SH2] of 3 μ M and 2 μM, respectively, [ATP] > 250 μM led to complete conversion of monomers to assemblies within 5 minutes, whereas significantly slower rates of conversion were observed with lower [ATP] (Fig. 2E, S17, Table S2. Visualization of assemblies using optical and fluorescence microscopy (with Alexa-647-labeled AtzC-SH2) revealed the existence of large (>10 μm) dendritic structures (Fig. 2F, G), whose formation could be observed in real time by adding kinase and ATP to a mixture of the two component proteins (Movie S1, Fig. S18).

Apparent hydrodynamic radius (Fig. 2F, G) and polydispersity measured with DLS (Fig. S19 and S20) could be controlled by varying the relative stoichiometry of the two components, and by using a weaker binding affinity variant of the SH2 domain fused to AtzC. A comparison of assembly formation trends for the lower (Fig. 2F) and higher affinity (Fig. 2G) SH2-domain-containing constructs shows that robust assembly formation is observed at nearly equal concentrations of the two components. Assemblies can be formed at concentrations as low as 50 nM (Dissociation constant, *K*_D_, for the weaker and tighter interactions were measured as ~40 and ~7 nM, respectively; Fig. S10), whereas when one component is present in excess, assembly formation is inhibited, as expected from our branch propagation design model (Fig. 1). Assembly formation by non-stoichiometric concentration combinations with the higher affinity SH2 domain variant (Fig. 2F, G) indicates that the inhibition caused by an excess of the binding partner is dynamic. Inhibition of assembly formation due to stoichiometric excess can also be overcome in an affinity-dependent manner: the zone of stoichiometries where assembly formation occurs is larger for the higher affinity SH2 domain variant (Fig. 2G) compared to the lower affinity variant (Fig. 2F). These results highlight the importance of high affinity in stabilizing the designed kinetically trapped aggregate state: under conditions of uneven stoichiometry (e.g. 250 nM AtzA-pY; 1000 nM AtzC-SH2) and in the absence of kinetic traps, all AtzA components should be bound by an excess of AtzC-SH2 domains, and no assemblies should result (expected particle size is <50 nm). This behavior is observed for the weaker-affinity SH2 variant at this stoichiometry (Fig. 2F). In stark contrast, for the high affinity SH2 variant (Fig. 2G), we observe micrometer-sized assemblies indicating the presence of aeolotropic kinetic trapping^44^ and network formation by clusters of tightly bound AtzA-pY-AtzC-SH2 assemblies (Movie S1).

We next investigated if the dynamic and dendritic structures observed in solution by optical and fluorescence microscopy (Fig. 2H, I) could form fractals on solid surfaces, and if the topology of the surface-adsorbed assemblies could be controlled by varying component stoichiometry. Due to the substantial increase of surface area derived from fractal patterns, surface-adsorbed fractals at the nanometer-micrometer scale are attractive design targets for applications in many fields like catalysis, fractal electronics, and the creation of nanopatterned sensors^4,5^. Assemblies with a chosen stoichiometry of components were generated in buffer, dropped on the surface of a silicon (or mica) chip, and the solvent was evaporated at room temperature (298 K) under a dry air atmosphere. Visualization of these coated surfaces using Helium Ion and Atomic Force microscopy reveals striking, intricately textured patterns that coat up to 100 μm^2^ areas (Fig. 3A-E). Various morphologies on the micron scale including rod-like, tree-like, fern-like, and petal-like were observed (Fig. 3A-E); image analysis revealed fractal dimensions between 1.4-1.5 (Fig. 3A,B) to the more Diffusion Limited Aggregation (DLA)-like 1.78 (Figs.3C,D, S21, and S22). Assembly sizes and fractal dimensions could be tuned by varying the stoichiometry of components (Fig. 3F), although some heterogeneity in morphologies was present in each sample. At 1:1 stoichiometry of the two components, DLA-like topologies with ~10 μm size were observed, whereas more dendritic assemblies were observed when unequal stoichiometry samples were used (Fig. 3F). Similarly, smaller assembly sizes resulted when the concentration of one component became limiting.

**Figure 3:**
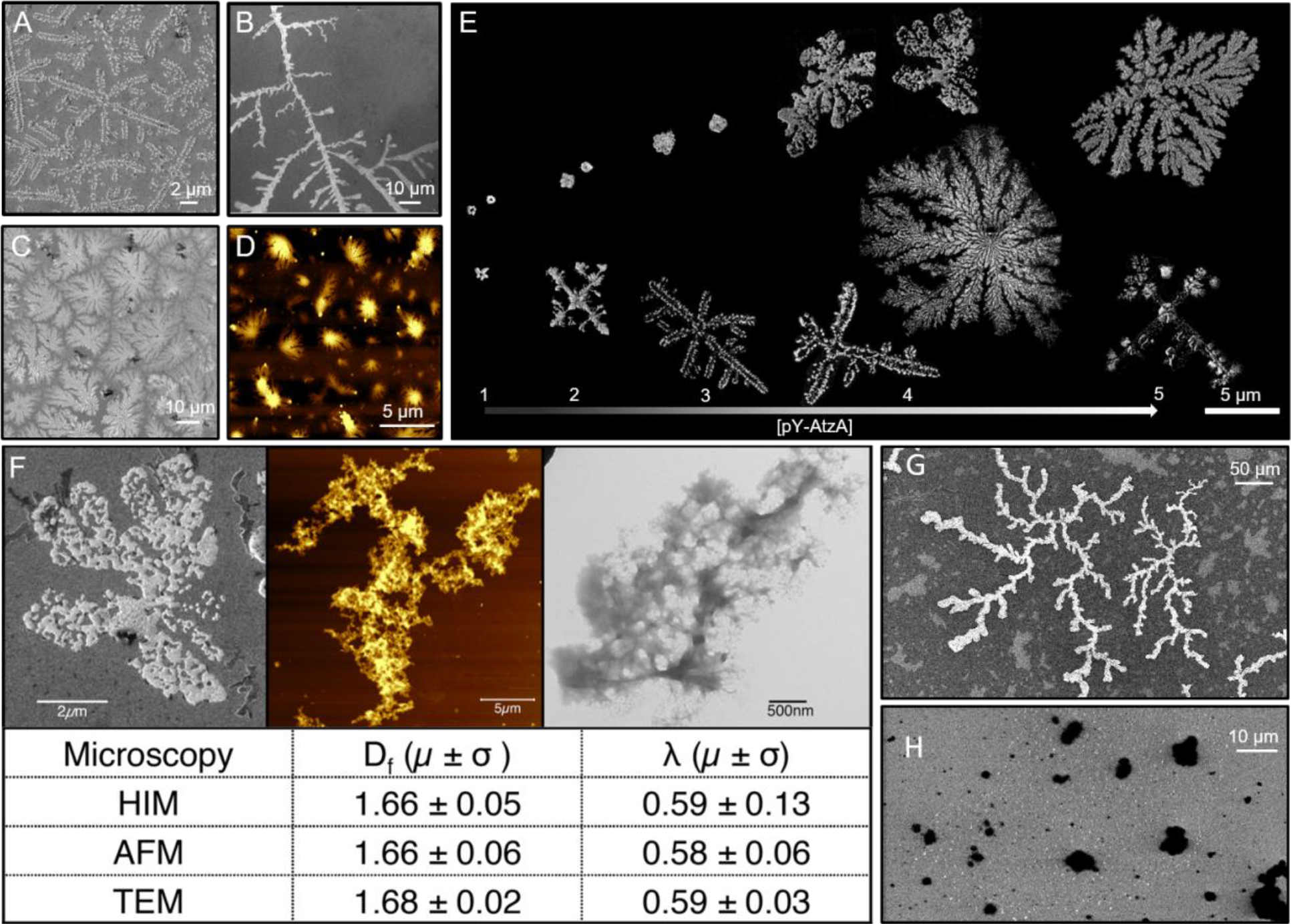
Assembly formation and characterization with Helium Ion Microscopy (HIM), Atomic Force Microscopy (AFM), and Transmission Electron Microscopy (TEM), all reveal fractal-like topologies on a surface, (**A to G**) Longer fractal-like structures, branch-like, and flower-like structures are seen in HIM (**A** to **C**) and AFM (**D**). (**E**) Representative HIM images for assemblies obtained at different concentrations of pY-AtzAM1 (250 nM-3 μM) while maintaining a fixed concentration of AtzCM1-SH2 (2 μM). Increasing concentrations of pY-AtzM1 result in larger assemblies with higher fractal dimensions. (**F**) D_f_ and λ, the fractal dimension and lacunarity of the images, are similar for images obtained from different microscopy techniques. HIM images show fractal-like assembly formation with pY-AtzAM1 and AtzCM1-SH2 (**G**), while the Gly-Ser-rich linker-containing variants form globular assemblies under these conditions (**H**).

Fractal patterns were not observed at any component stoichiometry without addition of ATP and Src kinase, with unphosphorylated proteins, or upon drying the buffer (to preclude precipitation-induced assembly formation by the salt in the buffer) demonstrating that fractal structures are formed by designed components (Fig. S23). Similarly, fractal topologies were not detected when long ((GSS)_10_), conformationally flexible Gly-Ser-rich linkers were used to fuse the SH2 domain and pY tag to AtzC, and AtzA, respectively. In mixtures of these proteins, a densely packed globular topology was detected with HIM, typical of amorphous precipitates (Fig. 3G,H, S24). Thus, the surface-induced patterns observed with designed AtzC and AtzA are selectively formed upon inter-component association in the designed geometries but not upon isotropic, random association as expected for the highly flexible Gly-Ser-rich linker-containing variants.

Transmission electron microscopy of designed AtzA-AtzC proteins also revealed branching, dendritic networks reminiscent of fractal intermediates observed in biosilica formation^14^ (Fig. S25). To further investigate the conformations of designed assemblies in solution and to obtain sufficiently high-resolution structures to test the validity of our design approach, we characterized the assemblies using cryo-electron tomography (cryo-ET; Fig. 4A-F, S26, Movies S2, S3). Assemblies generated by mixing 3 μM pY-AtzA and 2 μM AtzC-SH2 (or corresponding AtzA and AtzC fusions with Gly-Ser-rich linkers as controls) were blotted on a grid, frozen, and visualized on a cryo-electron microscope. Due to the increased image contrast from Volt phase plates in our microscope setup, pY-AtzA and AtzC-SH2 complexes in assembly tomograms were easily identified as density clusters. In contrast, constructs with Gly-Ser-rich linkers connecting pY and SH2 domain with AtzA and AtzC did not form porous clusters but instead (~90% of the sample) formed large, dense globular clumps (Fig S26B) where individual components were not resolvable (SI. 5.3). These large topology changes on the micron scale (as observed by both cryo-ET and HIM) upon conformational flexibility changes at the nanometer scale, further re-inforce the importance of directional association in our modular fractal assembly design framework.

**Figure 4:**
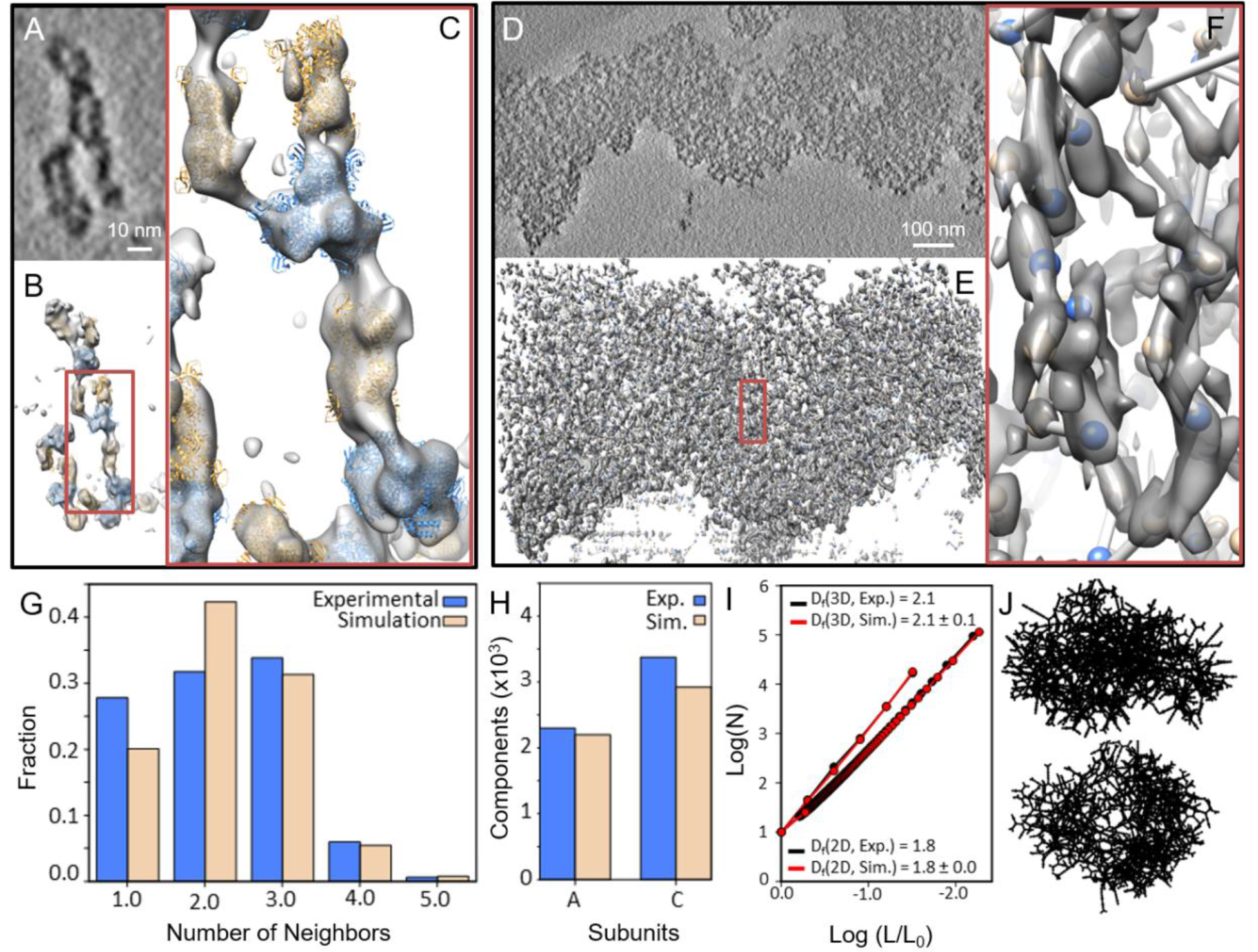
Assembly characterization with Cryo-electron Tomography, observed topologies in solution for a small (**A-C**) and large assembly (**D-F**), in which the subtomograms were extracted and fit with the Rosetta models. For the small assemblies, atomic resolution models of pY-AtzAM1 (blue) and AtzCM1-SH2 (tan) were fit to reveal the intercomponent connections along assembly branches. For the large assembly, due to the lower resolvability in this region of the sample, only geometric centers of density were used to assign to individual components (blue and tan spheres; SI 3.6) (**G**) Spatially proximal neighbor distribution from the Cryo-ET-derived images compared to simulated assemblies. (**H**) The relative component distribution in the cryo-ET image and from simulations. (**I**) Image analysis (2D), using a box counting method, of the Cryo-ET tomography subtomograms converted into 2D projections show similar fractal dimension. Standard deviation in simulation is calculated from 100 simulations comprised of ~5000 components. 3D box counting revealed similar fractal dimension (slope) between the experimentally observed and simulated assemblies. (**J**) Parameters for the simulation that match the experimental data are: P_term_: 0.1, C_frac_: 1.0, and *k*T: 9.0; two representative 2D projections with the matching parameters are shown.

Computational annotation of the density clusters formed by designed components in cryo-ET-derived images was performed based on individual molecular envelopes of components derived from Rosetta models of pY-AtzA and AtzC-SH2, respectively, to identify inter-component connections along assembly branches (Fig. 4A-C). The topology of the largest, nearly fully interconnected assembly based on electron density (Fig. 4D-F), consisting of approximately 6000 individual protein components, was further analyzed, and compared with an ensemble of simulated structures with approximately the same number of components. We compared the observed distributions of nearest-neighbor counts for AtzA-pY (Fig. 4G, Fig.S27), relative numbers of component types incorporated (Fig. 4H), angular distribution (C-A-C connections, Fig. S28), and the observed fractal dimension (Fig. 4I) of the assemblies with ensembles of structures generated using computational modeling (Fig. 4J) and found good agreement between the data and our simulations performed at specific parameter values (P_term_: 0.1, C_frac_: 1.0, and *k*T: 9.0). The observed nearest neighbor distribution for the AtzA-pY component shows that a large majority of these proteins are connected to 1, 2, or 3 neighboring AtzC-SH2, in agreement with the divalent connections envisioned in the design model and implemented in the simulated assemblies (Fig. 1). Additionally, a small but significant number of AtzA-pY proteins have 4 AtzC neighbors in both the computational ensemble and the cryo-ET images, which indicates physically unconnected components being proximal to each other in space due to the packing in the assembly, although a small number of monovalent connections cannot be definitively ruled out in the cryo-ET images (Fig. 4G). We found that the fractal dimensions from the cryo-ET images and simulations (2.1) show good agreement (Fig. 4I, J). The expected fractal dimension for a DLA-like cluster, which results from isotropic interactions, is 2.3 and the observed decreased fractal dimension (2.1) indicates the anisotropic nature^32–34^ of the underlying protein-protein interactions as encoded in the design approach (Fig. 1). Particle counting (and volume estimation) in a convex hull enclosing the largest assembly component yields an approximate local concentration of the proteins as ~600-700 μM, a ~125-fold increase compared to their bulk concentration (3 μM AtzA-pY and 2 μM AtzC-SH2). Particle density in the fractal assembly is ~70,000 particles/μm^3^ whereas calculated density of 2D and 3D crystalline lattices of similar volumes is estimated to be ~4000 and ~40,000 particles/μm^3^ (Fig. S1). The high particle density in the fractal while maintaining porosity leads to a high effective surface area, a characteristic feature of macroscopic fractal objects such as trees, and sponges. Although we could assign individual density clusters to individual components, the thickness of the ice in this region of the sample lowers resolvability and precludes direct measurement of orientation of the AtzA-pY and AtzC-SH2 components with respect to each other for comparison with Rosetta-calculated landscapes (Fig. 1). While there is significant heterogeneity in assembly sizes (~60% of the proteins adsorbed on the cryo-ET grid are parts of smaller assemblies) and topologies (Fig. S29), the observed increase in the effective concentrations concomitant with a high effective surface area with numerous solvent channels (Fig. 4A-F) indicates that induced fractal-like structure formation is a viable strategy to engineer protein assemblies with favorable sponge-like properties.

We next investigated if the observed textured, sponge-like topology, resulting in a high surface area:volume in the fractal assembly, endows it with similar enhanced material capture (“soaking up”) properties on the nanoscale as observed for macroscopic sponges. We reasoned that the lacunarity (“gappiness”) of the fractal structure and use of an excess AtzA-pY component under fractal-forming stoichiometries (Fig. 2F,G), would lead to several phosphopeptide sites on AtzA being open and accessible. The observed large pore sizes (Fig.4D-F) would enable access to these sites for molecular capture of nanometer-sized, macromolecular moieties bearing SH2 domains. In contrast, due to their dense, globular structure, amorphous assemblies generated with Gly-Ser-rich linker-containing components would have less available binding sites resulting in a lower loading capacity (Fig. S24, S26). To test the molecular capture properties of assemblies, we first used two fusion proteins in which macromolecular cargo proteins were fused to an SH2 domain: SH2-GFP, SH2-DhaA (an engineered DhaA enzyme for the degradation of the groundwater pollutant 1,2,3-trichloropropane (TCP)), and measured the amount of cargo proteins captured by fractal and globular assemblies generated using identical amounts of component proteins (Fig. 5). Indeed, fractal assemblies captured greater amounts of cargo, as evidenced by fluorescence (GFP) and enzymatic activity (DhaA) measurements, respectively (Fig. 5C). Fluorescence microscopy of SH2-GFP containing assemblies revealed that, as anticipated from cryo-ET studies, the immobilized cargo protein was distributed throughout the assembly, and localized to the surface, for fractal and globular assemblies, respectively (Fig. 5D-G). To develop a more broadly applicable approach for exploiting the efficient molecular capture properties of fractal assemblies, we generated and utilized a SH2-Protein A fusion protein to capture a fluorescent IgG antibody. As observed for SH2-GFP and SH2-DhaA, fractal assemblies can efficiently capture this antibody (Fig. 5H-K, S30). Furthermore, incubation of antibody-loaded assemblies with YopH phosphatase enzyme permits release of captured cargo antibodies (Fig. 5A-C). As all full-length IgG antibodies universally have the binding sites for Protein A (their Fc-domains), antibody-loaded fractal assemblies should enable (**a**) efficient molecular capture of a variety of macromolecular and small-molecule antigens, and (**b**) phosphorylation-dependent antibody purification^55–57^.

**Figure 5:**
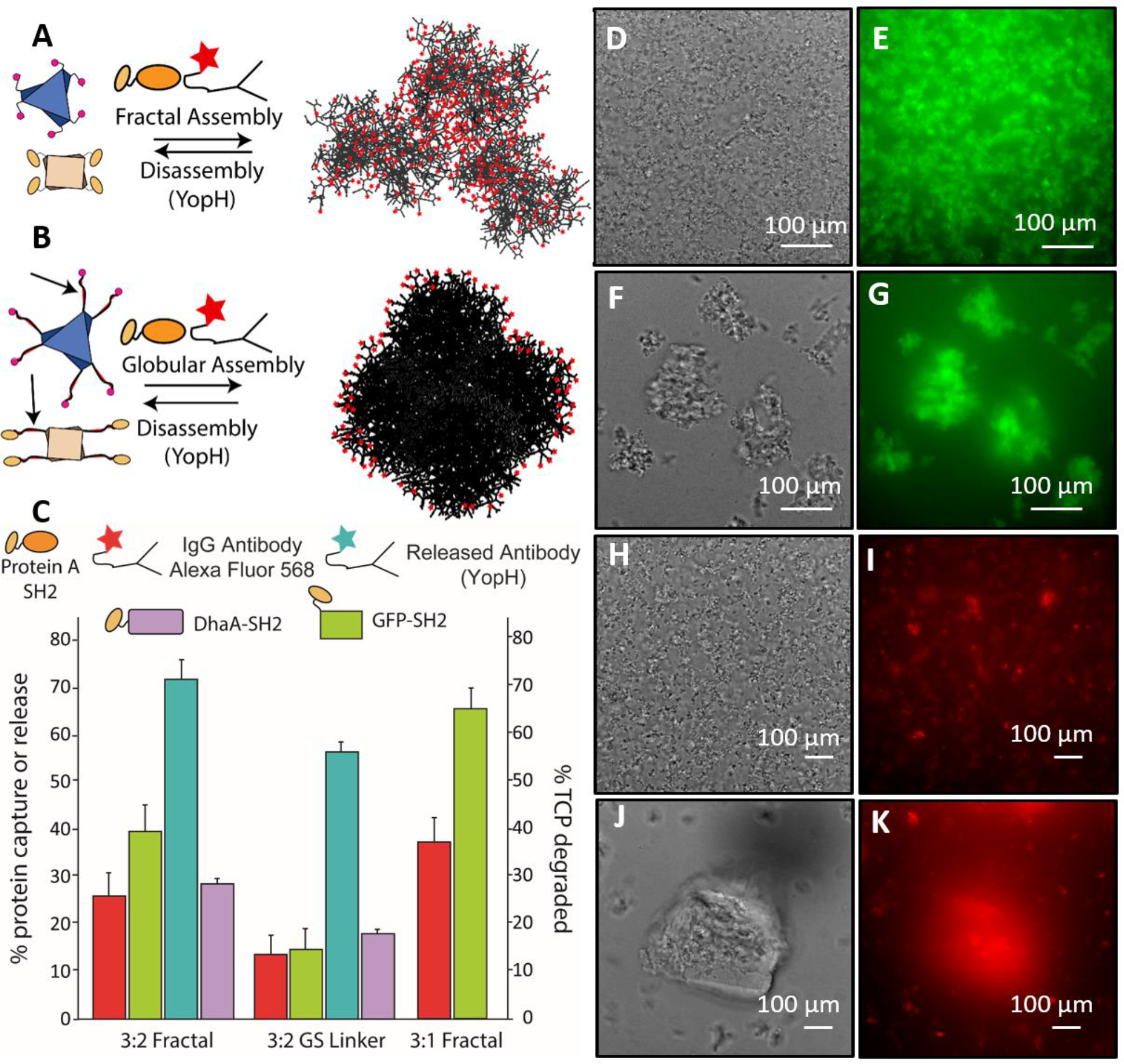
Fractal assemblies capture and release greater amounts of cargo compared to globular assemblies. Scheme for the envisioned reversible capture of cargo proteins for the (**A**) fractal and (**B**) globular structures. The red stars denote an example cargo protein (antibody). (**C**) % protein capture was measured for 3:2 fractal (assemblies obtained with 3 AtzA-pY: 2 AtzC-SH2), 3:2 GS linker (globular assemblies obtained at the same stoichiometry with fusion proteins containing long GS-rich linkers), and 3:1 fractal (assemblies obtained with 3 AtzA-pY: 1 AtzC-SH2). The 3:2 fractal captured more IgG antibody (red bars) and GFP-SH2 (green barss), and degraded more substrate TCP (purple bar; reflecting the higher capture efficiency of enzyme DhaA-Sh2) than the 3:2 GS linker assemblies. In addition, 3:2 fractal released more captured antibody compared to the 3:2 GS linker when incubated with YopH phosphatase (blue bars). (**D**, **E**) Confocal fluorescence microscopy images of the 3-component assembly with GFP-SH2 showing the topology of incorporation of GFP-SH2 in fractal and (**F-G**) the incorporation of GFP-SH2 into the globular assemblies. (**H-I**) the IgG antibody Alexa Fluor 568 incorporation into the fractal assembly and (**J-K**) the incorporation into the globular assembly.

In our design framework, fractal loading capacity is determined by the number and accessibility of open phosphopeptide binding sites in the assembly. Thus, assemblies formed by 3 (AtzA-pY):1 (SH2-AtzC) are expected to have a greater loading capacity compared to those formed by 3 (AtzA-pY):2 (SH2-AtzC). Indeed, as anticipated, more antibody was captured and released by the former compared to the latter (Fig. 5C), demonstrating that customized optimization of molecular capture-and-release of specific nanoscale objects should be possible by varying component stoichiometry to obtain the fractal properties on the nano-micrometer scales. Finally, we asked if the observed functional advantages of fractal topology over a globular one would extend to the capture and transport of small molecules within the assembly by measuring the efficacy of atrazine degradation. We incorporated as cargo AtzB – the third pathway enzyme, apart from AtzA and AtzC, required to convert atrazine to the relatively benign metabolite cyanuric acid (Fig. S31-S34). While both the fractal and globular assemblies appear to be more robustly active under harsh reactions compared to unassembled enzymes (Fig.S35), and when immobilized on a Basotect^®^ polymer foam (Fig. S36), both globular and fractal assemblies are equally active (Fig. S37). The significantly small size of atrazine (*Rg* < 1nm) and other metabolic pathway intermediates likely allows them to diffuse equally efficiently in either assembly as the smaller solvent channel size in the globular assembly may not be an impediment for small molecule guest as opposed to macromolecular guest molecules. Constructing fractal-like shapes with smaller sized proteins may allow access to smaller solvent channels.

In conclusion, our results demonstrate an approach by which fusion proteins may be designed to self-assemble into fractal-like morphologies on the 10 nm-10 μm length scale. The design strategy is conceptually simple, modular, and should be applicable to any set of oligomeric proteins featuring cyclic, dihedral, and other symmetries, such that multivalent connections, anisotropy and geometric degeneracy of binding can be used to controllably generate a broad range of sizes and morphologies of fractal shapes with proteins. In contrast with computational design of integer-dimensional protein assemblies where considerable remodeling of protein surfaces is necessary to meet the exacting geometric requirements for inter-component binding^24,25^, our design approach to obtain fractal-like morphologies involves few substitutions on protein surfaces. Instead, design goals are encoding high affinity via fusion of binding domains, binding anisotropy and geometric degeneracy via short, flexible loops (SI 5.1). Although we used SH2 domain-pY peptide fusions as the high-affinity modular connecting elements to endow phosphorylation responsiveness, the same design strategy should be applicable for the incorporation of other peptide recognition domains, responsive to other chemical or physical stimuli. The combination of multivalency and chain flexibility is a key determinant of other recently discovered phases formed by proteins, including droplets formed by liquid-liquid phase separation^58^. Our results show that this rich phase behavior of proteins^44^ also includes fractal-like morphologies that form colloidal particles with constituent microscopic molecular networks which may be visualized at high resolution using cryo-ET. Given the wide-ranging applications of fractal-like nanomaterials for molecular capture, further development in the design of protein-based fractals described here is expected to enable the production of novel classes of bionanomaterials and devices.

## Acknowledgments

The authors declare that all data supporting the findings of this study are available within the paper and its supplementary information files. Code used for coarse-grained simulations of assembly formation is available upon request. SDK and LWP acknowledge support from NSF (grant #1330760). NEH acknowledges the NSF Graduate Research Fellowship (#DGE-1433187). MC acknowledges grant #R01GM080139. Cryoelectron microscopy was supported by the Rutgers New Jersey CryoEM/ET Core Facility. We thank H. Cho, M. Liu, A. Permaul, O. Dineen, I. Patel, and R. Patel, for experimental assistance; K-B Lee, G. Montelione, and V. Nanda for technical advice, and D. Baker, V. Nanda for helpful discussions.

## Supplementary Information

Methods

Supplementary Discussion Supplementary References

Figures S1-S37

Table S1 to S3

Movies S1-S3

## Author Contributions

NEH, WAH, and SDK, designed the research. WAH designed proteins using Rosetta, developed fractal growth simulation methods and analyzed microscopy images. NEH, DZ, MES, and MK expressed and purified all the proteins used in the study, performed enzyme activity and assembly formation assays. NEH, VM, TG, LPF performed Helium Ion Microscopy. NEH, MP, PR and S.-H, Lee performed fluorescence microscopy and bright-field microscopy. DZ performed the DLS experiments. MK performed the BLI experiments. LY performed the TEM experiments. IMB performed confocal microscopy (DIC and fluorescence). AGD, LWP performed the polymer foam immobilization and activity assays. WD and MB performed the Cryo-electron tomography experiments and analyses. WAH and MC performed computational analyses of the Cryo-ET data. SDK, NEH and WAH wrote the manuscript. All authors commented on the manuscript.

