## Supplementary Information for "Stimulus-responsive Self-Assembly of Protein-Based Fractals by Computational Design"

**Title: Stimulus-responsive Self-Assembly of Enzymatic Fractals by Computational Design**

This PDF file includes:

Methods
Supplementary Discussion
Supplementary References
Supplementary Figures. S1 to S37
Supplementary Tables S1 to S3
Movies S1 to S3

### Methods

### INDEX


**SI 1 Computational methods**

- **SI 1.1** Preparation of scaffolds
- **SI 1.2** RosettaMatch: simultaneous fusion domain and peptide pair stitching
- **SI 1.3** Rosetta Design: interface design
- **SI 1.4** Generation of energy landscape using Rosetta FastRelax
- **SI 1.5** Coarse-graining AtzA-C oligomers for stochastic fractal growth simulations
- **SI 1.6** Stochastic fractal assembly simulation
- **SI 1.7** Temperature, concentration fraction, and termination parameter sweep
- **SI 1.8** Preparing fractal models for image analysis
- **SI 1.9** Preparing helium ion microscopy (HIM) images for image analysis
- **SI 1.10** Determining fractal lacunarity and 2-D fractal dimension with ImageJ
- **SI 1.11** Comparison of simulated and experimentally observed assemblies from Cryo-EM

**SI 2 Experimental characterization**

- **SI 2.1** Creation of the designed AtzA, AtzB, and AtzC fusion constructs
- **SI 2.2** AtzA and AtzC expression and purification
- **SI 2.3** AtzB expression and purification
- **SI 2.4** Src human kinase, super binder SH2 domain, SH2-DhaA expression and purification
- **SI 2.5** YopH phosphatase construct, expression, and purification
- **SI 2.6** Biuret hydrolase and cyanuric acid hydrolase expression and purification
- **SI 2.7** Enzyme-linked immunosorbent assay (ELISA)
- **SI 2.8** Bio-layer interferometry (BLI)
- **SI 2.9** Phosphorylation, assembly formation, and disassembly
- **SI 2.10** Dynamic light scattering (DLS)
- **SI 2.11** DLS Inhibition Experiment
- **SI 2.12** DLS Titration Experiment
- **SI 2.13** DLS Kinetics (varying ATP) Experiment

**SI 3 Microscopy experiments**

- **SI 3.1** Transmission electron microscope (TEM)

**SI 3.2** Atomic force microscopy (AFM)
**SI 3.3** Helium ion microscopy (HIM)
**SI 3.4** High-resolution fluorescence microscopy
**SI 3.5** Cryo-EM Tomographic tilt series acquisition and reconstruction
**SI 3.6** Cryo-EM AtzAM1 and AtzCM1 model fitting and statistical analysis
**SI. 3.7**. Confocal Microscopy

**SI 4. Enzymatic and fractal-incorporation assays**
**SI 4.1** Enzymatic activity measurements using the Berthelot assay
**SI 4.2** Temperature stress activity assays
**SI 4.3** Shaking stress activity assays
**SI 4.4** Construction and assay of Basotect® polymer foam with trapped assemblies and free enzymes
**SI 4.5** Gfp-SH2 incorporation using fluorescence
**SI 4.6** Dhaa-SH2 incorporation using TCP degradation assays
**SI 4.7** Goat anti-mouse IgG Antibody incorporation

**Supplementary discussion**
**SI 5.1** Molecular features determining fractal formation
**SI. 5.2** Fractal dimension and lacunarity
**SI. 5.3** Comparison of control (GS-rich-linker containing) and designed assembly topologies
**SI 5.4** Evaluating the effects of AtzB-SH2 on overall fractal structure and topology

**SI 1. Computational methods**

**(SI 1.1) Preparation of scaffolds –** Crystal structure files for AtzA (PDB:4V1X) and AtzC (PDB:2QT3) were subject to several preparatory scripts to clean, symmetrize, and process the files for Rosetta Design[1–3]. The processed crystal structure files were then subject to a Rosetta Fast Relax[4] protocol to obtain starting structures of sufficiently low Rosetta Energy to serve as starting structures. To prepare a library of conformations predicted to propagate into a fractal structure, we first aligned the proteins along paired $C_2$ symmetry axes (A+B chains for both AtzA and AtzC). We then translated AtzC along the aligned $C_2$ symmetry axis until the backbone (N, C, Ca, O) atoms of each structure were at least 3Å apart to find the minimum starting distance between the centers of mass (125Å). From the minimum starting distance we translated AtzC(monomer) in intervals of 1Å to a maximum distance of 145Å. For each translated AtzC(monomer) position we rotated the AtzC(monomer) about the $C_2$ symmetry axis at a core set of angles i.e., 180 ± (0, 35.25, 54.75, 90, 125.25, 144.75, and 180). As detailed below, we used this scaffold library to stabilize simultaneously a subset of rotations about the paired C-symmetric axes known to favor 2D or 3D crystal geometries (Fig. S1). These values were chosen as they were expected to allow propagation of the assembly in a stochastic manner – we hypothesized that a mixture of connections at a set of angles should result in a fractal assembly (Fig. S1).

**(SI 1.2) RosettaMatch: simultaneous fusion domain and peptide pair stitching –** After visual inspection of the two-component scaffold placements, we noted the accessibility of the AtzA N-terminus and the AtzC C-terminus along the $C_2$ symmetry axis (chains A+B). Therefore, we decided to fuse the N-terminus of an fyn-SH2 super-binder (PDB:1A0T) to the C-terminus of AtzC and the C-terminus of the fyn-SH2 peptide binding partner to the N-terminus of AtzA. To achieve simultaneous fusion, we converted the SH2-peptide crystal structure into an all-Cα 'ligand' file and used RosettaMatch[5] (Fig. S2) with geometric constraints to sample all sterically feasible rigid body placements of the SH2-peptide between each AtzA-AtzC pair in the two-component scaffold library. RosettaMatch requires a set of 6 geometric degrees of freedom, imposed as constraints in order to orient a given "ligand" (SH2-peptide Ca trace) model with respect to each desired amino acid contact. Geometric constraints used to coordinate the SH2 domain-peptide complex for simultaneous fusion were derived from backbone atom positions and orientations using a non-redundant protein library generated by the RCSB-PDB[6]. From N to C terminus, regardless of secondary structure, we collected distances and angles between backbone atoms (Cα, nitrogen, and carboxyl carbon) up to and including 7 residues downstream (sequence-space) of each residue along the primary structure. The averages and standard deviations of these distributions were used to place geometric constraints between residues of the AtzA-AtzC termini and the all-Cα SH2-peptide ligand to force only geometrically allowable backbone fusions. The full-atom SH2-peptide crystal structure was re-threaded back onto each of the matched SH2-peptide ligands creating 7,005 models with paired termini in proximally close and geometrically favorable positions. Rosetta GeneralizedKIC (kinematic loop closure)[7] was used to covalently link the paired termini and generate 3 potential linker-models for each matched SH2-peptide model, creating a library of 21,015 fused and bound AtzA-AtzC pairs. Geometric constraint files and GenKIC XML protocol files are provided in a supplementary data zipped file.

**(SI 1.3) Rosetta Design: interface design –**A Rosetta-based design protocol was used to stabilize the novel interfaces formed from the placements of SH2-peptide and protein monomers obtained using RosettaMatch (followed by loop closure) as described in SI 1.2. In the design protocol we allowed linker residues (±4 residues around the fusion site) and AtzC-SH2 domain interface residues (Cα-Cα distance <6Å) to change residue identity. All backbone atoms with the exception of the linker residues were constrained with atom-coordinate constraints to favor the SH2-peptide placements determined in the RosettaMatch step. Mutations at the linker region, ±4 residues around the fusion site, which alleviated steric clashes with the backbone or sidechains of AtzC/SH2 domain were accepted such that they stabilized multiple intercomponent angles of attachment (Table S1). All other mutations were reverted to native residue identities before a subsequent round of repack and energy minimization[8]. Based on our calculations, we selected 5 variants of AtzA fused to the SH2-peptide recognition sequence and 5 variants of the AtzC fused to the SH2 domain for experimental characterization. Table S1 shows a list of mutations and the value of rotation stabilized for each design. Rosetta command lines and XML files used for design are provided in supplementary zipped file.

**(SI 1.4) Generation of energy landscape –** To evaluate the energy landscape of the designed component pair (pY-AtzAM1 and AtzC-SH2M1) along the symmetrically aligned rotation-translation degrees of freedom we performed a Rosetta Symmetric FastRelax protocol on conformations of AtzA-AtzC pairs. Each conformation was represented by three parameters, translation (*d*), rotation (*θ*), and axis-binding preference (vertex or edge centered). Parameter values d in range 120Å to 145Å in steps of 1Å; θ in range 0° to 360° in steps of 5° were used to identify their predicted preferred binding modes. We generated this energy profile and plot in (Figure 1E-F) conformations whose evaluated binding energy scored better than the wild-type components (504 models). This binding energy was calculated by separating the binding components to a distance of >500 Å and repacking the components. Binding energy values were used to compute a Boltzmann-weighted probability distribution used in coarse grained simulations described below. The Rosetta command lines and XML files for energy landscape generation are provided in supplementary zipped file.

**(SI 1.5) Coarse-graining AtzA-C oligomers for stochastic fractal growth simulations –**we used a coarse-grained representation of our symmetric oligomers by reducing each chain to 10 representative points in space (60 and 40 for whole hexamer and tetramer respectively). To coarse-grain we used a K-means-like clustering algorithm to place the 10 points at locations with the highest concentration of Cα atoms in each monomer (chain A). We then calculated and applied the symmetric transform to the 10 representative points to obtain a coarse-grained representation of each oligomer (hexamer and tetramer). When each representative point is converted to a sphere with a 12Å radius, the coarse-grained model effectively mimics the overall shape and size of the full-atom model.

**(SI 1.6) Stochastic fractal assembly simulation–** In order to predict the supramolecular structure and topology we developed a stochastic fractal assembly simulation protocol that utilizes Boltzmann weighted probability distributions for an ensemble of predicted low-energy binding modes along the $C_2$-symmetry axes of the AtzA-AtzC pairs as described in SI 1.5. The algorithm operates by starting with one oligomer (AtzA for this study) and attaches each complementary oligomer layer-by-layer. The Boltzmann probability distribution was used to decide how the oligomers in each layer were placed. A few key assumptions were made during the simulations. We assumed: 1) The symmetric divalent connection along a $C_2$-symmetry axis (two chains of pY-AtzA bound two chains of AtzC-SH2) would be energetically more likely than the monovalent connection formed between just one chain from each oligomer—reducing the probability of monovalent connection to an insignificant value. 2) Flexibility in the linker region would only lead to variations along the $C_2$-symmetry axis via the translation and rotation parameters used in design—maintaining the inherent symmetry found in either oligomer. 3) Mixed vertex-centered and edge-centered species could occur around a single AtzA. This would lead to a substructure where two AtzC oligomers have a 180°-angle about an AtzA component, different from the 120°-angles when pure edge-centred and vertex-centered binding geometries are considered. 4) Changes in size and topology would arise from concentration changes of the enzymes and would need to be represented in the algorithm. 5) During fractal growth it is possible (and likely) that oligomers in one layer could come within 125Å (minimum connected distance) of other oligomers within another layer even if they are not directly connected. The details of this algorithm are described below.

Energy landscapes calculated in SI 1.4 were used to stochastically propagate the coarse grained A-C components during simulation. We varied the $kT$ term to obtain a total of 5 different Boltzmann weighted probability distributions ($kT$ = 1, 3, 5, 7, and 9). Propagation was achieved by alternating layers of AtzA and AtzC components starting from an initial seed component (pY-AtzA in this study) which would continue until either placement of new components was determined either impossible or improbable or an external criterion was met (number of layers, size of particle, etc.). The propagation algorithm involves 6 steps at any given layer:

1) Using $C_{frac}$, randomly select a fraction of the components from the previous layer (or seed if 1st iteration) to continue propagation. Components not chosen will cease to propagate for the remainder of the simulation. For example, if $C_{frac}$ =0.33, a third of the available connection points are considered propagatable. $C_{frac}$ models the relative stoichiometry of the two components.

2) Iterate over the selected components determined in step 1 and:

2a) Randomly select an available $C_2$-symmetry axis of the individual selected in (2).

2b) Using a Boltzmann weighted probability distribution of possible $d$-$\theta$-axis conformations, attempt to apply a component with an orientation at random along the $C_2$ symmetric axis (2a).

2c) Choose whether or not to keep the selected $C_2$-symmetry axis (2b) based on a termination probability ($P_{term}$). The termination probability models the binding affinity. For example, a weaker binding SH2 domain variant would have a higher $P_{term}$ compared to a tight-binding variant.

2d.1) If (2c) passes the term, apply the rigid body transformation (*d* and *θ*) and append as a member of the next layer.

2d.2) If (2c) fails the term, mark the $C_2$-symmetry axis (2a) of the individual selected in (2) as unviable and continue.

3) Repeat 2a-d.1 until all $C_2$-symmetry axes of individual (2) are exhausted.

4) Perform a coarse grid-based clash check to ensure new layer members are sterically non-clashing with any components of the assembly.

5) Repeat Steps 2-4 until all of the components chosen in (1) are exhausted.

6) Move to the next layer.

**(SI 1.7) Temperature, fraction, and term parameter sweep –** Varying the fraction $C_{frac}$ (1) and termination probability $P_{term}$ (5-6b) parameters gave rise to changes in topology and structure. We created 100 fractal models for each combination of $C_{frac}$ (range: 0.1-1.0, interval: 0.1) and $P_{term}$ (range: 0.0-0.9, interval: 0.1) using the 5 different Boltzmann weighted probability distributions (with varying temperature)—creating 50,000 total fractal assemblies. These parameter sweep simulations were performed for 15 layers for each parameter combination. We analyzed each particle's individual size, number of layers, AtzA branch ratio (number of AtzC units bound to a unit of AtzA, lacunarity ($\lambda$), and dimensionality ($D_f$) from a 2D image. For every combination of temperature, fraction, and term we averaged the data across the 100 fractal assemblies. The results can be found in Figure S5 and S6. In the range $C_{frac}$ values 0.5-1 and $P_{term}$ values 0.0-0.4, particle diameter, branch ratio, fractal dimension, and lacunarity increase with increasing values of $C_{frac}$ and decreasing values of $P_{term}$. Particle diameter and branch ratio decrease with increasing values of $kT$ while lacunarity increases. Fractal dimension first increases until $kT = 5$ and then decreases. In this range the number of layers are shown to remain the same.

**(SI 1.8) Preparing fractal models for image analysis –** Each coarse-grained assembly model obtained above was analyzed using a PyMOL script that would color the assembly components black, convert the background to white, show as spheres of scale 12Å, orient the image such that the longest diameters are in the X-Y plane, remove the glossy lighting and shine from the sphere models, and finally ray-trace render the image. This PyMOL script is provided in supplementary zipped file.

**(SI 1.9) Preparing helium ion microscopy (HIM) images for image analysis –** HIM images were loaded into ImageJ[9]. The initial image contrast was enhanced with 5-20% saturated pixels setting; this can be achieved with Process -> Enhance Contrast. We then create a new blank (black) image with the same pixel dimensions as the HIM image. Gaussian noise is added to the blank image with a standard deviation 5-10 (Process -> Noise -> Add Specified Noise). Background noise is subtracted from the HIM image using the noisy blank image (Process -> Image Calculator -> set Image1 to HIM image and image2 to noisy blank -> set operation to subtract). Finally, we create a binary image from the processed HIM image with subtracted background. The resulting image contains white protein islands on a black background. Individual fractal islands are then copy/pasted into a new blank (black) image using the polygon selection tool and are ready for fractal analysis.

**(SI 1.10) Determining fractal lacunarity and 2-D fractal dimension with ImageJ -** The FracLac package[10] designed for ImageJ was used to determine both the 2D lacunarity and fractal dimension ($D_f$). With FracLac mode on, outside of the standard parameters, we checked the 'alternate random generator' box and allowed the minimum pixel size to be 1, and the color code was turned off. We then ran in batch-mode to process all of the fractal images. ImageJ outputs four files: summary, box count per grid, scan types, and batch data. Lacunarity and dimension were taken from the summary file for the parameter sweep while the 2D log vs log plot values were taken from the box counting grid file ($\varepsilon$ and $F$ ).

**(SI 1.11) Comparison of simulated and experimentally observed assemblies from Cryo-EM –** Fitting of the experimentally computed protein density (from Cryo-electron tomography) resulted in Cartesian coordinates representing the center of mass of the oligomeric components. To compare the experimental results to simulation we ran the simulation until at least a total of 5000 components were present in the model and calculated the geometric centers for all oligomeric components in the coarse-grained assembly to create new center-of-mass models. Using the experimentally derived Cartesian coordinates and the center-of-mass models we performed a computational analysis (SI 3.6) to evaluate the fractal size, nearest component neighbor distances, and relative AtzA-AtzC ratio (Fig. 3H,I). We analyzed the 3D fractal dimension (Fig. 3J) with a 3D box counting program that counts the number of geometric centers within a scaling (doubling) box size. The 2D fractal dimension (Fig. 3J) was calculated in the same way as previously mentioned (Fig. S4). We found highest agreement of simulations with $kT = 9$, $P_{term} = 0.1$, and $C_{frac} = 1.0$. An array of fractal images that represent the average fractal for each value of $P_{term}$ and $C_{frac}$ at $kT = 9$ can be found in Figure S6.

**SI 2. Experimental characterization methods**

**(SI 2.1) Creation of the designed AtzA, AtzB, and AtzC fusion constructs –** The DNA sequence of the full-length *atzA* was amplified from the *pMD4::atzA*; *atzB* amplified from *pAAJLS3::atzB;* and *atzC* was amplified from *pKK223-3::atzC*.[9–12] The Src kinase activator phosphopeptide sequence, EPQYEEIPIYL, was created by ordering two complementary primers that formed a linear fragment encoding the peptide sequence, used with the amplified *atzA* gene and inserted into the linearized *pET15b+* vector through Gibson Assembly.[13] The Fyn SH2 superbinder gene was ordered as a gBlock fragment[13,14] and inserted into *pET29b+* (linearized with *Nde*I and *Xho*I) using Gibson Assembly.The Fyn SH2 amplified gene was designed to be placed on the C-terminal side of the *pET15b+::atzB* and *pET29b+::atzC* with a flexible GSS linker between the proteins. The Fyn SH2 superbinder amplified gene *SH2* and the *atzC* amplified gene were both inserted into the *pET29b+* linear vector using Gibson Assembly. The *atzBSH2* fusion gene was ordered as a Gibson fragment[13] and inserted into the *pET15b+* linear vector using Gibson Assembly. Point mutations were introduced using the QuickChange Site-Directed Directed Mutagenesis Kit (Agilent Technologies) to create the final designs for AtzA and AtzC models. DNA sequencing was used to confirm proper insertion and mutations (Genscript).

**(SI 2.2) AtzA and AtzC expression and purification –** The *pET15b+::atzApep* and *pET29b+::atzCSH2* plasmids were co-transformed into *Escherichia coli* BL21 (DE3) with *pAG* plasmid containing genes for the chaperone proteins, *groEL* and *groES* [15].For expression of the AtzA models a 10 mL LB culture with 30 µg/mL of chloramphenicol and 100 µg/mL of ampicillin was inoculated with a single colony and incubated overnight at 37°C and 250 rpm. For the expression of the AtzC models a 10 mL LB culture with 30 µg/mL of chloramphenicol and 50 µg/mL of kanamycin was inoculated. After growing overnight, the 10 mL cultures of the AtzA and AtzC models were used to inoculate 500 mL of LB media, which was grown at 37°C to an $OD_{600}$ of 0.5-0.6, at which point the expression of chaperones was induced with the addition of 1% (wt/vol) L-arabinose and grown for an additional 1-2 hours at 16°C . Expression of the AtzA and AtzC models was then induced with 0.1mM IPTG (isopropyl-β-D-thiogalacto-pyranoside) and grown overnight at 16°C. All subsequent steps were performed at 4°C. Cells were centrifuged at 6,000 x g for 30 min. Cell pellets were re-suspended in 30 mL of 25 mM HEPES, 200 mM NaCl, 5% glycerol, 40 mM imidazole, pH 7.5, and lysed by sonication. Cell extracts were obtained by centrifugation at 50,000 x *g* for 30 min at 4°C. Protein purification was performed using 5 mL Ni-NTA agarose resin (Qiagen) equilibrated with 10 mL of 25 mM HEPES, 200 mM NaCl, 5% glycerol, 40 mM imidazole, pH 7.5. The lysate was applied to the resin, the resin was washed with 45 mL of the same buffer, and the protein eluted with 20 mL of 25 mM HEPES, 200 mM NaCl, 5% glycerol, 400 mM imidazole, pH 7.5,. The purified protein was buffer exchanged (PD10-desalting column, GE Healthcare #17085101) into 50 mM HEPES, 100 mM NaCl, 5% glycerol, pH 7.4 (HNG). AtzA was expressed in high yields and precipitated if imidazole was not removed immediately after elution from the Ni-column. No precipitation was observed upon exchange of AtzA into HNG buffer and AtzC variants did not precipitate in either buffer (for hours-days) at 4°C. Proteins were frozen using liquid nitrogen and stored at -80°C.

**(SI 2.3) AtzB expression and purification –** The *pET15b+::atzBSH2* plasmid was transformed into *E.coli* BL21 (DE3) cells. For expression of AtzB, a 10 mL LB culture with 100 μg/mL of ampicillin was inoculated overnight at 37°C and 250 rpm. The 10 mL overnight culture was used to inoculate 500 mL of LB media which was grown to an $OD_{600}$ of 0.5-0.7 and induced with 1 mM IPTG and grown overnight at 16°C. The same purification protocol for the AtzA and AtzC models was used for AtzB. AtzBSH2 did not express if grown with zinc sulfate, as had been done customarily in previous literature.[11]

**(SI 2.4) Src human kinase, super binder SH2 domain, SH2-DhaA expression and purification –** The expression plasmid for Src human kinase[16] (gift from John Chodera, Nicholas Levinson, and Markus Seeliger. Addgene plasmid # 79700 was co-transformed with the expression plasmid for *Yersinia* YopH protein tyrosine phosphatase (PTPase)[16] (gift from John Chodera, Nicholas Levinson, and Markus Seeliger, Addgene plasmid # 79749) into *E. coli* Rosetta2 (DE3) (Novagen). For Src kinase expression a 10 mL LB culture with 50 μg/mL spectinomycin and 100 μg/mL of ampicillin was inoculated with a single colony and incubated overnight at 37°C, 250 rpm. The overnight culture was used to inoculate 500 mL of LB media which was grown to an $OD_{600}$ of 0.5-0.7 and induced with 1mM IPTG and grown overnight at 18°C. The super binder SH2 domain and SH2-DhaA were transformed into E. coli BL21 (DE3) and expressed in the same way as the Src kinase above. Purification for the Src kinase was performed similarly and with the same buffers as AtzAM1, AtzBSH2, and AtzCM1. While, the super binder SH2 domain and SH2-DhaA were purified with the same purification protocol but with the following buffers: a wash buffer containing 137 mM NaCl, 2.7 mM KCl, 10 mM $Na_2HPO_4$, 2 mM $KH_2PO_4$, pH 7.4, 20 mM imidazole and an elution buffer containing 137 mM NaCl, 2.7 mM KCl, 10 mM $Na_2HPO_4$, 2 mM $KH_2PO_4$, pH 7.4, 200 mM imidazole. All proteins were buffer exchanged into HNG, frozen in liquid nitrogen and stored at -80°C.

**(SI 2.5) YopH phosphatase construct, expression, and purification –** The linear catalytic domain *YopH* gene (residues 164-468) was amplified from *pET13S-A::YopH*[16] and inserted with Gibson Assembly into a linearized pET15b+ vector. A 10 mL LB culture with 100 μg/mL of ampicillin was inoculated with a single colony and incubated overnight at 37°C. The expression and purification protocol is the same as the protocol used for the Src kinase.

**(SI 2.6) Biuret hydrolase and cyanuric acid hydrolase expression and purification –** Biuret hydrolase (BH)[17] expression strain (*E. coli* DH5α) and the *Moorella* Cyanuric acid hydrolase (CAH)[18] strain (*E.coli* BL21 (DE3)) were provided by Dr. Larry Wackett. A 10 mL culture with 50 μg/mL of kanamycin was inoculated for both BH and CAH and incubated at 37°C until $OD_{600}$ of 0.5-0.7 and induced with 1 mM IPTG for 4 hours at 37°C, 250 rpm. The expression and purification protocol is the same as the protocol used for the Src kinase.

**(SI 2.7) Enzyme-linked immunosorbent assay (ELISA) –** Phosphorylated AtzAM1 (pY-AtzAM1) was loaded onto clear flat-bottom immuno 96-well plates (Thermo Scientific item # 442404) at 20μg/mL and 1.25μg/mL in 50μL 1X PBS (Gibco pH 7.4, #10010023) overnight at 4◦C. Plates were rinsed twice in 200μL 1X TBS (Biorad #1706435). 1% BSA in TBS 0.05% Tween 20 was used to block wells at 200μL block solution for 1.5hr at 25°C under gentle agitation. Anti-phosphotyrosine 4G10 Platinum HRP conjugate (EMD #16-316) was diluted 1:5000 in 1% BSA TBS 0.05% Tween 20 and loaded onto the well at 25°C for 1.5hr under gentle agitation. Excess anti-phosphotyrosine was washed off with 200μL of TBS 0.05% Tween

20 in triplicate. To detect bound antibody, 100μL of TMB substrate reagent (Biolegend #421101) was added to each well and incubated for 5 minutes at 25°C. 100μL of TMB stop solution (Biolegend #423001) was added to the wells. Absorbance was read at 450nm using the Tecan Infinite M200 Pro plate reader.

**(SI 2.8) Bio-layer interferometry (BLI) –** AtzAM1 was phosphorylated using the conditions described below. pY-AtzAM1 was then biotinylated at 10mM Sulfo-NHS-Biotin (APExBIO) for 30min at 25°C. Excess biotin was buffer exchanged with a PD-10 desalting column (GE Healthcare) equilibrated with HNG. Biotinylated pY-AtzAM1 was loaded onto streptavidin (SA) coated biosensors (ForteBio) and used for BLI. AtzCM1 was flowed in from 4nM to 4μM. BLI experiments were performed using the BLItz System (ForteBio).

**(SI 2.9) Phosphorylation, assembly formation, and disassembly –** The phosphorylation protocol was based upon Src kinase activity assay by Sigma (Catalog # S1076). In a final reaction volume of 150μL, 3μM AtzAM1 was mixed into 1X Kinase Activity Buffer (4mM $MgCl_2$, 2.5mM $MnCl_2$, 0.25mM DTT, 5mM MOPS, 2.5mM glycerol-2-phosphate, 1mM EGTA, 400nM EDTA, pH 7.6), 2.5 mM $MnCl_2$,HNG, 2 mM ATP, 800ng Src kinase, and incubated for 7 – 16 hr at 25°C for phosphorylation to occur. After phosphorylating, AtzCM1 was added to a final 2μM concentration. Assembly was allowed to form at 2hr 25°C. Disassembly was performed by adding 4.8μg of YopH phosphatase into the 150μL reaction mixture after assembly formation occurred. Size measurements using DLS were performed to determine assembly formation/disassembly.

**(SI 2.10) Dynamic light scattering (DLS) –** 50 μL of an assembly sample was used for size determination using a Malvern Zetasizer and a quartz cuvette (ZEN2112, Malvern). Ten spectra measures were recorded for eleven replicates at 25 °C. The standard operating procedure accounted for 5% glycerol in solution.

**(SI 2.11) DLS Inhibition Experiment -** 6 μM pY-AtzAM1 was phosphorylated (1X KAB, 2 mM ATP, 1 mM DTT, HNG, 1 μg Src kinase) in a reaction volume of 75 μL. Incubation time was overnight at 25°C. SH2 or SH2-DhaA was added to each sample at 0 μM, 3 μM, 6 μM, 9 μM, 12 μM, 15 μM, 18 μM final concentration and allowed to “block” binding sites on the pY-AtzAM1 for 1 hr at 25°C. AtzCM1 was added to each sample at 2 μM final concentration. Therefore, the final concentrations of all components was 3 μM pyAtzA, 1 μM AtzCM1, 0 μM - 18 μM SH2 or SH2-DhaA. The sample was incubated for 2 hr at 25°C. DLS was performed to analyze assembly sizes. DLS was performed at 25°C, 50 μL/sample volume, in a low-volume quartz sizing cuvette (Malvern; ZEN2112) using a Zetasizer Nano ZS (Malvern). Measurements were performed in triplicates while each sample was read and averaged 15 times. This protocol was repeated at a final concentration of 1 μM pyAtzA, 0.66 μM AtzCM1, 0 μM -6 μM SH2-DhaA. Curve fitting was performed in MATLAB (R2016b; Mathworks) using the general model:

$$f(x) = \frac{A}{1 + e^{-k*(x-x_0)}} + B$$

where *A*, *B, k,* $x_0$ are constants. Adjusted $R^2$ was used to determine model validity. Inhibition concentration 50 (IC50) was determined based upon concentration of inhibitor that resulted in assembly size of 100nm measured.

**(SI 2.12) DLS Titration Experiment –** 6 μM, 3 μM, 1.5 μM, 0.5 μM, 0.1 μM pyAtzA was phosphorylated (as described previously) with an incubation time of overnight at 25°C. Either AtzCM1 wildtype (WT) or AtzCM1 superbinder (SB) was added to each sample at 2 μM, 1 μM, 0.5 μM, 0.25 μM, 0.50 μM final concentration. The sample was allowed to incubate for 2 hr at 25°C. Therefore, the final concentrations of all components was from 3 μM – 0.05 μM pyAtzA, 2 μM – 0.05 μM AtzCM1-WT or AtzCM1-SB. DLS was performed at 25°C, 50 μL/sample volume, in a low-volume quartz sizing cuvette (Malvern; ZEN2112) using a Zetasizer Nano ZS (Malvern). Measurements were performed in duplicate with each sample read and averaged 15 times.

**(SI 2.13) DLS Kinetics (varying ATP) Experiment –** An assembly mixture of 3 μM non-pyAtzA and 2 μM AtzCM1 was prepared (as described previously) and syringe-filtered at 0.22 μm. To each 50 μL reaction volume, 1.2 μg of src kinase was added. Size was monitored continuously for 30 min at 25°C in a low-volume quartz sizing cuvette (Malvern; ZEN2112) using a Zetasizer Nano ZS (Malvern) at 50 μL/sample. Measurements were performed in triplicates. Each sample was read and averaged five times over the course of 25 seconds for a single time point. Curve fitting was performed in MATLAB (R2016b; Mathworks) using sloping spline function, with varying smoothing parameters. Adjusted $R^2$ was used to determine model validity.

**SI 3. Microscopy methods**

**(SI 3.1) Transmission electron microscope (TEM) –** Assembly (3 µM pY-AtzAM1 and 2 µM AtzCM1) and non-assembly (3 µM non-pyAtzA and 2µM AtzCM1) samples were mixed, and diluted ten-fold in deionized water. The diluted samples were applied to the carbon-coated FCF400-Cu grids (Electron Microscopy Sciences, Hatfield, PA) which had been glow-discharged for two hours under UV light to render the grids hydrophilic and adsorptive. A drop of sample (~5uL) was added on a piece of wax film and the grid was placed onto the sample droplet for absorption for two minutes. Excess sample solution was removed with a filter paper. A drop (~5uL) of 1% uranyl acetate was dropped on the wax paper and the grid was placed onto the staining solution droplet for two minutes to stain. Excess staining solution was removed by blotting with a filter paper, the grids were allowed to air dry for two minutes. Images were collected on JEOL 1200EX electron microscope with AMT-XR41 digital camera.

**(SI 3.2) Atomic force microscopy (AFM) –** The assemblies were directly visualized by non-contact mode atomic force microscopy (AFM) Parks Systems. Samples were prepared by depositing 20 µls of sample on silicon wafer and incubated for 5 minutes. After incubation, the silicon was washed with deionized water to remove salt and air dried overnight at 25°C. Assemblies were visualized by an AFM (Parks System). The AFM was used in non-contact mode (330 kHz resonant frequency and 42 N/m spring constant, PPP-NCHR Park Systems, #610-1051). Images were taken with 2048x2048 pixels with scan rates of 2 µm/s to 30 µm/s. The AFM images analysis was performed using Gwyddion software[19].

**(SI 3.3) Helium ion microscopy (HIM) –** The AFM sample preparation on a silicon wafer was used for HIM. Imaging was done on the Carl Zeiss Orion Plus Helium Ion Microscope (Carl Zeiss Microscopy, Peabody, MA) operating at 30 KeV acceleration voltage with a beam currents of about 1 pA. Most samples did not exhibit significant charging therefore electron flood gun was not used for charge neutralization. The vacuum reading in the analysis chamber during imaging was $2x10^{-7}$ torr.

**(SI 3.4) High-resolution fluorescence microscopy –** For the growth video, 20 µL of 3 µM AtzAM1 and 2 µM AtzCM1 sample (with all the required buffers as described previously) was deposited on a glass cover and 0.2 µm of Src kinase was added to the sample to allow for assembly formation to occur. The sample was monitored for an hour. For the 3-component assembly image (3 µM pY-AtzAM1, 1 µM AtzBSH2, 2 µM AtzCM1) the AtzBSH2 protein was dye labeled with the Alexa Fluor™ 647 NHS Ester (Succinimidyl Ester, ThermoFisher Scientific #A2006) and buffer exchanged into HNG with a PD10-desalting column. Fluorescent images along with bright-field images were collected. Images were captured using a Nikon Ti-E inverted microscope. A Coherent Genesis laser at 567 and Coherent Obis Laser at 647 were used for fluorescent imaging, using 1mW power. Images for the assemblies with antibody fluorescence was taken using a 2048x2048 pixel resolution. A 561 nm laser at 12 mW was used and imaged with a 100x TIRF high NA (1.49) oil immersion objective. Samples were placed on a bacto (tm) agar pad at 1.5% w/v (150 mg/10mL). The agar pad was hardened in a gene frame on a 25 mm coverslip and sealed with an 18 mm coverslip on top.

**(SI 3.5) Cryo-EM Tomographic tilt series acquisition and reconstruction –** For cryo-electron tomography, an AtzAM1 and AtzCM1 assembly sample was mixed with 10 nm gold fiducial markers to facilitate alignment in data processing. An aliquot of 3.5ml sample was applied to 2.0/1.0mm Quantifoil holey grids (Quantifoil, Germany) and plunge frozen using a Leica EM GP plunger (Leica). Tomographic tilt series acquisition was performed on a Talos Arctica microscope (Thermo Fisher) operated at an acceleration voltage of 200kV. This microscope was equipped with a field-emission gun, Volt phase plates, Gatan postcolumn energy filter and a K2 summit direct electron detector. Tilt series were collected at 39,000x microscope magnification with -0.5 µm defocus using FEI Tomography software. The sampling of the data was calibrated to be 3.49 Å/pixel. Typically, a tilt series ranged from -60° to 60° at 3° step increment. The accumulated dose for each tilt series was 60 electrons/Å2. Tilt series were aligned based on fiducial gold markers using the IMOD package[20]. 3D tomograms were obtained by weighted backprojection of aligned tilt series. Visualization and annotation of the 3D volumes were done in Chimera[21].

**(SI 3.6) Cryo-EM AtzAM1 and AtzCM1 model fitting and statistical analysis –** AtzAM1 and AtzCM1 complex subtomograms were extracted from 3D tomograms and bandpass filtered to reduce high frequency noises and low frequency gradient from ice thickness variation. Centers of AtzAM1 and AtzCM1 densities were identified as peaks within solid voxel clusters that were approximately sizes of an AtzAM1 hexamer, or an AtzCM1 tetramer. Potential free AtzAM1 or AtzCM1 complexes that were too close to a neighboring voxel peak (<120Å) were removed. Assignment of AtzAM1 or AtzCM1 to an identified voxel cluster was done by applying the condition that AtzAM1 and AtzCM1 alternate in a chain. Densities that had three or more linkers to neighbors were assigned to be AtzAM1. Linear, unbranched assemblies were assigned by first determining identity of one end based on cross-correlation scores between the end peak densities and AtzAM1 or AtzCM1 models computed from their PDB structures. Assignment conflicts were resolved by pruning along the branches in the order of intensity values. The above protocol was first applied to a small assembly, and optimized and validated by human visual inspection before it was used on larger assemblies. Coordinates and connection information of each AtzAM1 or AtzCM1 complex in an assembly were extracted and used for statistical analysis and for comparison to simulation data. The volume of the assembly is defined by the volume of the convex hull that encloses all determined AtzAM1 or AtzCM1 molecule.

**(SI 3.7) Confocal microscopy fluorescent images of fractal and globular assembly with GFP-SH2 and Goat anti-mouse IgG Antibody -** Fluorescently tagged samples were placed in chamber slides and allowed to air dry overnight. Fluorescent images were acquired using a spinning disc confocal microscope (Olympus DSU-IX81) fitted with 482nm and 543nm excitation filters and emission filters of 536nm and 593nm, respectively. Sample images were obtained using the 3D image capture function (Z-stacks) with an approximate depth of 200μm at 1μm intervals (step size) using an oil immersion objective (Olympus UPlanFL N 40X/1.3 Oil) and 300ms as exposure time. Image processing was performed with SlideBook 5.0 (3i, Intelligent Imaging Innovations).

### SI 4 Enzymatic and Fractal-incorporation Assays

**(SI 4.1) Enzymatic activity measured using the Berthelot assay** – Assembled enzyme samples (1.5 μM AtzAM1, 0.5 μM AtzBSH2, and 1 μM AtzCM1) were made by incubating the enzymes in 1X kinase activity buffer (with no DTT), 2.5 mM MnCl2, HNG, 0.2 μM Src kinase, and 2 mM ATP in a total volume of 500 μl at 25°C for 4 hours. The unassembled enzyme samples were prepared using the same conditions, except no ATP was added to the sample. DLS was performed to verify assembly formation. 10 μL of 20 mM Atrazine dissolved in methanol was added to each 500 μL sample, for a final concentration of 400 μM atrazine, and another sample with the same conditions had no substrate added in order to establish a baseline measurement. Each condition was done in triplicate. After the addition of substrate, the samples are shaken at 100 RPM for 1.5 hr at 25°C. 140 μL of each sample is transferred to

PCR tubes, then boiled at 99°C for 1.5 minutes, and then cooled at 4°C. The 140 µl were transferred to 1.5 mL microcentrifuge tubes and spun down at 20,000 rcf for 20 minutes to remove precipitated protein. 80 µl of the supernatant was used for the following steps. 1µg per 20 µL of sample of CAH and 1µg per 20 µL of sample of BH was added to each sample. The samples were incubated at 25°C for 2 hours to allow for the complete conversion of the cyanuric acid to ammonia by CAH and BH. The Berthelot assay was performed in triplicate on the resulting samples to determine the production of ammonia. For every mole of cyanuric acid produced, one mole of ammonia was assumed to have been produced. 20 µL of each sample was added to a 96-well plate (Greiner half area clear #675101). 60 µL of solution A (0.05 g/L sodium nitroprusside and 10g/L phenol) was added and mixed into every sample. Then 80 µL of solution B (5 g/L NaOH and 8.4 mL/L bleach) was added and mixed into every sample. The samples were incubated for 30 minutes at 25°C for a blue color to develop. The absorbance at 630 nm was read using Tecan Infinite M200 Pro plate reader. The extinction coefficient was determined using standards of cyanuric acid at known concentrations in the enzyme activity buffer that had been reacted with the BH and CAH for 2 hours.

**(SI 4.2) Temperature stress activity assays** – Assembled and unassembled enzyme samples were made as described above and incubated at 25°C for 4 hours to allow full assembly formation. The assemblies were then incubated at the following temperatures: 25°C, 40°C, 45°C, 50°C, 55°C, and 60°C for fifteen minutes, and cooled back to 25°C before the addition of 400 µM atrazine. After atrazine was added, the enzyme activity assay was performed as described above.

**(SI 4.3) Shaking stress activity assay** – Assembled and unassembled enzyme samples were made as described above and incubated at 25°C 4 hours. Both samples were shaken at 50, 100, 150, 200, 225, and 250 RPM 25°C for 1 hour before any addition of atrazine. 400 µM atrazine was added to the samples and shaking continued at their respective shaking speeds for 1.5 hour. The rest of the activity assay protocol was conducted the same as described above.

**(SI 4.4) Construction and assay of Basotect® polymer foam with trapped assemblies and free enzymes** – Hydrolyzed TEOS was prepared by combining 7 ml TEOS (Aldrich #131903), 3 ml water, and 0.04 ml 0.1N hydrochloric acid and stirring the solution for 2 hr at room temperature[22]. Basotect® polymer foam (Procter and Gamble UPC# 0 37000 43515 0) was cut into 2.0 x 2.0 x 0.3 cm squares with a razor and 0.250 ml of assemblies or free enzyme solution was spotted onto each 2 x 2 cm face of the foam squares. Aliquots (1.0 or 0.5 ml) of hydrolyzed TEOS were diluted with HNG buffer to a final volume of 10 ml (10% or 5% TEOS). A single application of 5% or 10% hydrolyzed TEOS solutions was done with a small paint brush (Richeson 95822). The TEOS was allowed to set for 2 h, and then liquid was squeezed out of each foam square and total protein concentration in the liquid was measured with the Bradford assay (BioRad #500-0006). To assay activity in the embedded foam, 1 ml of 150 µM atrazine in 1X phosphate buffered saline (pH 7.4) was soaked into the foam squares and incubated for 1.5 hour at 25°C. Liquid was squeezed out after incubation and boiled as above to inactivate eluted enzymes. Cyanuric acid produced during the incubation was assayed as described except that the Berthelot reactions were conducted in 10 x 4 x 45 mm cuvettes (Sarstedt #67-742) and read using a Beckman DU 640 spectrophotometer.

**(SI 4.5) GFP-SH2 incorporation assays –** AtzAM1 and AtzCM1 (along with AtzA/AtzC extended linker versions for globular assemblues) assemblies were formed in a reaction volume of 3 mL, 15 µM AtzAM1 and 10 µM AtzCM1 into 1X Kinase Activity Buffer, 2.5 mM $MnCl_2$, HNG, 2 mM ATP, 0.2 µM Src Kinase, and allowed to form for 10 minutes before the addition of 1.8 µM Gfp-Sh2 protein, and incubated for 4 hr at 25°C for phosphorylation to occur. Samples were spun down for 2 minutes (500 x *g*) and supernatant measured in a black half-area microplate (excitation 395 nm, emission 509 nm) with a gain of 140 on a Tecan Infinite M200 Pro plate reader.

**(SI 4.6) DhaA-SH2 incorporation assays –** AtzAM1 and AtzCM1 (along with AtzA/AtzC extended linker versions for globular assemblies) assemblies were formed in a reaction volume of 3 mL, 15 µM AtzAM1 and 10 µM AtzCM1 into 1X Kinase Activity Buffer, 2.5 mM $MnCl_2$, HNG, 2 mM ATP, 0.2 µM Src Kinase, and allowed to form for 10 minutes before the addition of 1.8 µM of Dhaa-Sh2, and incubated for 4 hr at 25°C. Assemblies were spun down and pellet resuspended with 10 mM TCP and incubated for 1 and 16 hr. Assemblies were spun down again and supernatant was measured at A560 nm.

**(SI 4.7) Goat anti-mouse IgG incorporation assays** - AtzAM1 and AtzCM1 (along with AtzA/AtzC extended linker versions for globular assemblies) assemblies were formed in a reaction volume of 1.8 mL, 12 μM AtzAM1 and 8 μM AtzCM1 into 1X Kinase Activity Buffer, 2.5 mM $MnCl_2$, HNG, 2 mM ATP, 0.2 μM Src Kinase, and allowed to form for 10 minutes before the addition of 8 μM of ProteinA-Sh2, and incubated for 4 hr at 25°C. Assemblies were spun down 20,000 x *g* for 20 min in order to measure fluorescence in supernatant. For disassembly assays with YopH, assembly pellets were spun down 20,000 x g for 20 min, supernatant removed, pellets washed with HNG buffer, and spun down again to remove wash, and resuspended in HNG containing YopH. Assemblies were left shaking at 100 RPM for 12 and 24 hrs at 25°C. Assemblies were spun down again and supernatant measured for released antibody.

### Supplementary Discussion

#### (SI. 5.1) Molecular features determining fractal formation

It has been demonstrated[23] that atomic-level control over component placement is necessary to achieve via computational design, periodic, regularly ordered 2D protein lattices[24] or closed form 3D icosahedra[25]. In contrast, where 2D lattices and 3D closed form assemblies require exacting orientation and rigidity of inter-protein components, fractal assemblies require a degree of flexibility leading to degeneracy of binding modes and anisotropy at the interface of protein components. However, the amount of flexibility needs to be tuned: too little and crystal lattices will form (Fig. S1), too much and globular protein agglomerates will result (as observed for our control assemblies involving long loop connectors).

To obtain a fractal assembly with protein components, based on our data, we hypothesize that three tunable factors contribute: valency, affinity, and flexibility.

Valency, the measure of possible favorable connections between protein components, contributes to the amount of branching as well as the orientation of the inter-protein components. With homomeric $D_2$ and $D_3$ protein components, we anticipated the $D_3$ (AtzA) to make up to 6 connections to the $D_2$ (AtzC) component which is capable of 4 connections. If the affinity of the inter-protein connection is sufficiently strong, and the length of the bridging interactions is kept short we could observe an avidity effect between components—where two bridges (divalent connection) are formed between two components (Fig. 1). The formation of divalent bridging connections, localized to $C_2$ sub-symmetries of D-symmetric proteins, can greatly reduce the flexibility between connected protein components while providing high affinity. In this way, avidity and symmetry together can be utilized to introduce orientational anisotropy and rigidity of the inter-component connections.

To promote avidity, we chose strong (nM affinity) peptide-binding motifs which could be fused to the D-symmetric protein building blocks. During design, to ensure that any divalent connections made between components were restricted to connections along the $C_2$ sub-symmetry axes, we imposed design constraints on the fusion linker lengths—maintaining that no additional residues would be added beyond the residues found in the crystallographic structure files creating a direct fusion (0-residue linker) and introducing Gly and Ser residues (GGS) to promote (limited) flexibility.

**(SI. 5.2) Fractal dimension and lacunarity**

The fractal (Hausdorff-Besicovitch or box counting) dimension[26], a general measure of how the size of an object scales as a function of the size of its building blocks, has been used to characterize simulated/mathematical fractal patterns[27,28] as well as real-world statistical fractals including peptide-based fractals obtained on a surface and imaged with AFM[29]. Intuitively, the dimension of an object can be thought of as the scaling observed for change in its overall size (or mass) upon changing the size of its unit building block. For example, a square has dimension two because its mass (proportional to area) grows with a scaling exponent of 2 with the length of its side (if length is increased $n$-fold then area increases by $n^2$ -fold). A cube has a dimension of three because its mass (volume) will increase by $n^3$ times when length of its side is increased $n$-fold. For some objects, this scaling is observed to non-integer and captures how the object occupies space. For example, a curve with a fractal dimension 2.1 fills space very much like an ordinary 2-dimensional surface, but a curve with a fractal dimension of 2.9 folds to fill space nearly like a 3-dimensional volume. However, any arbitrary curve is not fractal – it has to follow a scaling equation throughout the space it occupies as described below. Another intuitive heuristic is that fractal dimension can be thought of as a measure of the “roughness” of an object’s periphery (“Clouds are not spheres, lightening is not a straight line” – Mandelbrot). The surface of the human brain, for example, is fractal.

To calculate fractal dimension for a fractal $S$, we consider the object $S$ lying on an evenly spaced grid (of size $L_0$) , and count how many boxes are filled by elements of $S$. The box-counting dimension is calculated by seeing how this number changes as we make the grid size

($L$) finer by applying a box-counting algorithm as shown in Fig. S4 (using an image obtained from microscopy in our study). Thus, the fractal dimension, $D_f$, [30–34] calculated from box counting is obtained:

$$Log(N) = D_f . Log(\frac{L}{L_0})$$

The existence of a straight line with a single slope in the above plot indicates that the object is fractal and indicates statistical self-similarity. If different slopes are obtained at different scales, the object is considered multi-fractal.

Another quantity that captures the "gappiness" or "holeyness" of a fractal shaped object is lacunarity (λ)[30]. In box counting analyses, lacunarity λ for each grid of calibre ε (= $L/L_0$) is calculated from the standard deviation, σ, and mean, μ, for pixels per box. That is, there is a λ value for each ε in each series of grid sizes in each grid orientation in a set of grid orientations.

$$\lambda_{\varepsilon,g} = CV_{\varepsilon,g^2} = \left(\frac{\sigma_{\varepsilon,g}}{\mu_{\varepsilon,g}}\right)^2$$

This value is averaged over all orientations to obtain the reported lacunarity, λ, value. We used the implementation in ImageJ software to calculate fractal dimension and lacunarity.

We note that in our analyses, fractals formed by the same components can vary in shape and dimension from isolated assembly on the surface (island) to island as well as in solution. However, despite inter-island variations, every island is self-similar (with the same fractal dimension). Similar type of topological diversity (and uniformity within islands) was also found in studies of silk protein sericin[34], where variation in fractal dimension of observed protein islands was detected depending on the surface conditions but each island was self-similar. For all 2D image analyses in this paper, we derived the fractal dimension (slope), scalability (linear range), and lacunarity from 2D image analysis using ImageJ. Due to the island-to-island variation, all 2D-analyzed $D_f$ and λ values reported in this work are an average of at least 5 individual islands (as many as 20 images were used when available).

When comparing the Cryo-ET data to the computational simulation results, projections were made to be analyzed with the same 2D image analysis. Additionally, 3D-fractal dimension analysis was performed with an in-house 3D box-counting algorithm that works in the same way that 2D image analysis does except the two-dimensional boxes are replaced with three-dimensional cubes (voxels) during the scaling analysis.

**(SI. 5.3) Comparison of control (GS-rich-linker containing) and designed assembly topologies using cryo-ET**

Although we could resolve the cryo-ET-derived density of the fractal assemblies (Fig. S26A and Fig. S26C, respectively), the globular (GS-rich-linker containing) assemblies varied too greatly in topology across samples to analyze—the majority of these images were dominated by dark shadowy particles too dense to obtain meaningful assignments of density to individual protein components (Fig. S26B). However, a few images (<10%) from the GS-linker rich set had small resolvable nm-scale regions where density could be interpreted and assigned to individual protein components (Fig. S26D). For these images, we compared the average monomer-monomer distance across 5 control (GS-rich) and 5 fractal-shaped assemblies (Fig. S27) on the nm-scale. In the fractal-shaped assemblies, the inter-monomer distance is tightly clustered (134 ± 2 Å) among images of large (>25 nm size) assemblies (~40% of the set), suggesting uniformity of inter-component connections in agreement with the design conception. In contrast, in the resolvable parts of the control assembly tomograms (<10% of the entire imaged sample), we see three different types of structures: dispersed assembly (inter-monomer distance ~157Å), fractal-similar assemblies (~134Å), and densely packed globular ball-like structures (~125 Å). These data suggest that GS-rich linker conformational flexibility (1) abrogates anisotropy of inter-component interactions and (2) allows reorganization of assembly structure to yield a more globular structure on the micron scale driven by non-specific protein-protein sticking. The robust catalytic activity of the control assembly (Fig. S37) demonstrates that the observed topologies in the control tomograms are not the result of protein unfolding but are in fact, mediated by the SH2 domain-pY peptide interactions.

**(SI. 5.4) Evaluating the effects of AtzB-SH2 on overall fractal structure and topology**

Upon addition of 1 unit AtzB-SH2 to the 3A:2C ratio (added before phosphorylation) we find that the observed fractals show a marked decrease in both $D_f$ and $\lambda$ compared to the 3A:2C fractals (Fig.S32). When the concentration of A is increased in the three-component assembly we once again see a decrease in $D_f$; however, we also see a decrease in $\lambda$ (Table S3, Fig. S31, S32).

The observed $D_f$ and $\lambda$ in the three-component assembly resemble the $D_f$ and $\lambda$ values in the two-component fractals with low concentrations of A relative to C (Fig. S21, Table S3 ). These findings, combined with the observed incorporation of dye-labeled AtzB-SH2 data (Fig. S34 and Fig. 35D), suggest that AtzB-SH2 is competing for locations to bind the SH2-peptide fused to AtzAM1 and is further changing the structural features of the assembly.

### Supplementary Information References


1. Dantas, G., Kuhlman, B., Callender, D., Wong, M. & Baker, D. A large scale test of computational protein design: Folding and stability of nine completely redesigned globular proteins. *J. Mol. Biol.* **332,** 449–460 (2003).
2. Tinberg, C. E. *et al.* Computational design of ligand-binding proteins with high affinity and selectivity. *Nature* **501,** 212–6 (2013).
3. Meiler, J. & Baker, D. ROSETTALIGAND: Protein-small molecule docking with full side-chain flexibility. *Proteins Struct. Funct. Genet.* **65,** 538–548 (2006).
4. Tyka, M. D. *et al.* Alternate states of proteins revealed by detailed energy landscape mapping. *J. Mol. Biol.* **405,** 607–618 (2011).
5. Zanghellini, A. *et al.* New algorithms and an in silico benchmark for computational enzyme design. *Protein Sci.* **15,** 2785–2794 (2006).
6. Berman, H. M. *et al.* The protein data bank. *Nucleic Acids Res.* **28,** 235–242 (2000).
7. Mandell, D. J. & Kortemme, T. Backbone flexibility in computational protein design. *Current Opinion in Biotechnology* **20,** 420–428 (2009).
8. Fleishman, S. J. *et al.* Rosettascripts: A scripting language interface to the Rosetta Macromolecular modeling suite. *PLoS One* **6,** (2011).
9. Mandelbaum, R. T., Allan, D. I. & Wackett, L. P. Isolation and Characterization of a Pseudomonas sp . That Mineralizes the s -Triazine Herbicide Atrazine. *Appl. Environ. Microbiol.* **61,** 1451–1457 (1995).
10. Seffernick, J. L., Johnson, G., Sadowsky, M. J. & Wackett, L. P. Substrate specificity of atrazine chlorohydrolase and atrazine-catabolizing bacteria. *Appl. Environ. Microbiol.* **66,** 4247–4252 (2000).
11. Seffernick, J. L. *et al.* Hydroxyatrazine N-ethylaminohydrolase (AtzB): An amidohydrolase superfamily enzyme catalyzing deamination and dechlorination. *J. Bacteriol.* **189,** 6989–6997 (2007).
12. Shapir, N., Osborne, J. P., Johnson, G., Sadowsky, M. J. & Wackett, L. P. Purification,

substrate range, and metal center of AtzC: The N-isopropylammelide aminohydrolase involved in bacterial atrazine metabolism. *J. Bacteriol.* **184,** 5376–5384 (2002).

32. Fairbanks, M. S., McCarthy, D. N., Scott, S. A., Brown, S. A. & Taylor, R. P. Fractal electronic devices: Simulation and implementation. *Nanotechnology* **22,** (2011).

33. Murr, M. M. & Morse, D. E. Fractal intermediates in the self-assembly of silicatein filaments. *Proc. Natl. Acad. Sci.* **102,** 11657–11662 (2005).

34. Khire, T. S., Kundu, J., Kundu, S. C. & Yadavalli, V. K. The fractal self-assembly of the silk protein sericin. *Soft Matter* **6,** 2066 (2010).

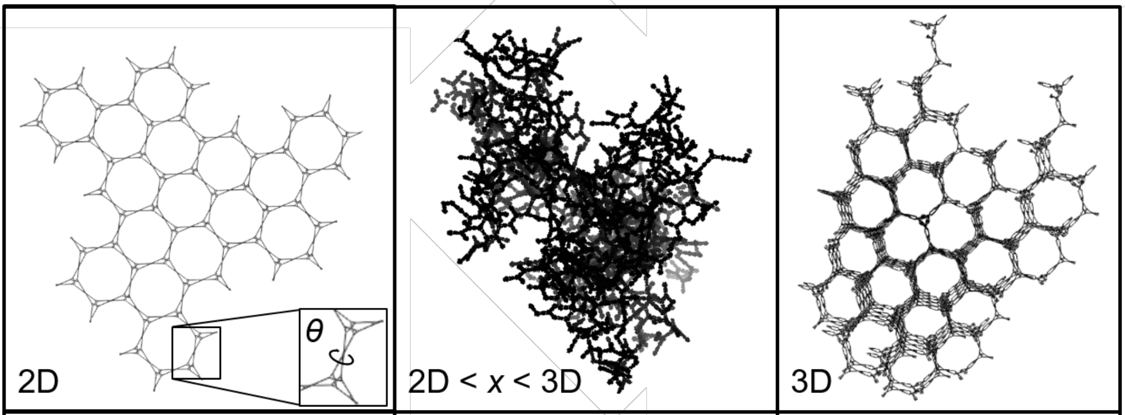


| | | |
|---|---|---|
| Choose 1: $\theta \in \{0, 90, 180, 270\}$ | Choose >1: $\theta \in \{0, 35.25, 54.75, 90, 125.25, 144.75, 180, 215.25, 234.75, 270, 305.25, 324.75\}$ | Choose 1: $\theta \in \{35.25, 54.75, 125.25, 144.75, 215.25, 234.75, 305.25, 324.75\}$ |
| ~4,000 units/μm$^2$ | ~70,000 units/μm$^3$ | ~40,000 units/μm$^3$ |

**Fig. S1. Scheme for designing arboreal fractal morphologies.** Predicted assembly topology based on values of *θ*. For 2D and 3D crystal geometries *θ* would have to be one discrete angle from the angles listed. A fractal would form from stochastic combinations of propagatable *θ* values in the range $0 \leq \theta < 360$. For a cube with length 1μm we calculated the number of theoretical total connected protein units that the 2D and 3D assembly built with AtzC-SH2 and AtzA-pY could occupy to be 4000 and 40,000 units respectively and compared that to the number of units/μm$^3$ observed for the Cryo-ET characterized fractal assembly (70,000 units/μm$^3$).

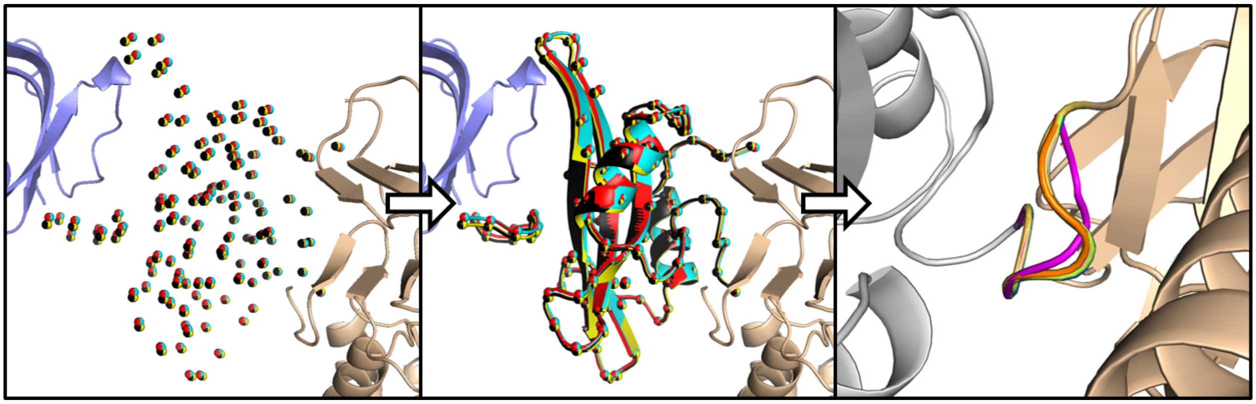

**Fig. S2. Flowchart of interface and linker design method.** (A) The SH2 and peptide binding partner alpha carbons are converted into a ligand file which is used by RosettaMatch to determine placements of the domain-peptide pair (colored red, yellow, cyan, and black for 4 unique solutions) within placements of AtzA and AtzCmonomers (with varied *d* and *θ*). B) The ligand is converted to a full atom model by threading (using the alpha carbons) followed by C) fusion loop closure using Kinematic Loop Closure (KIC).

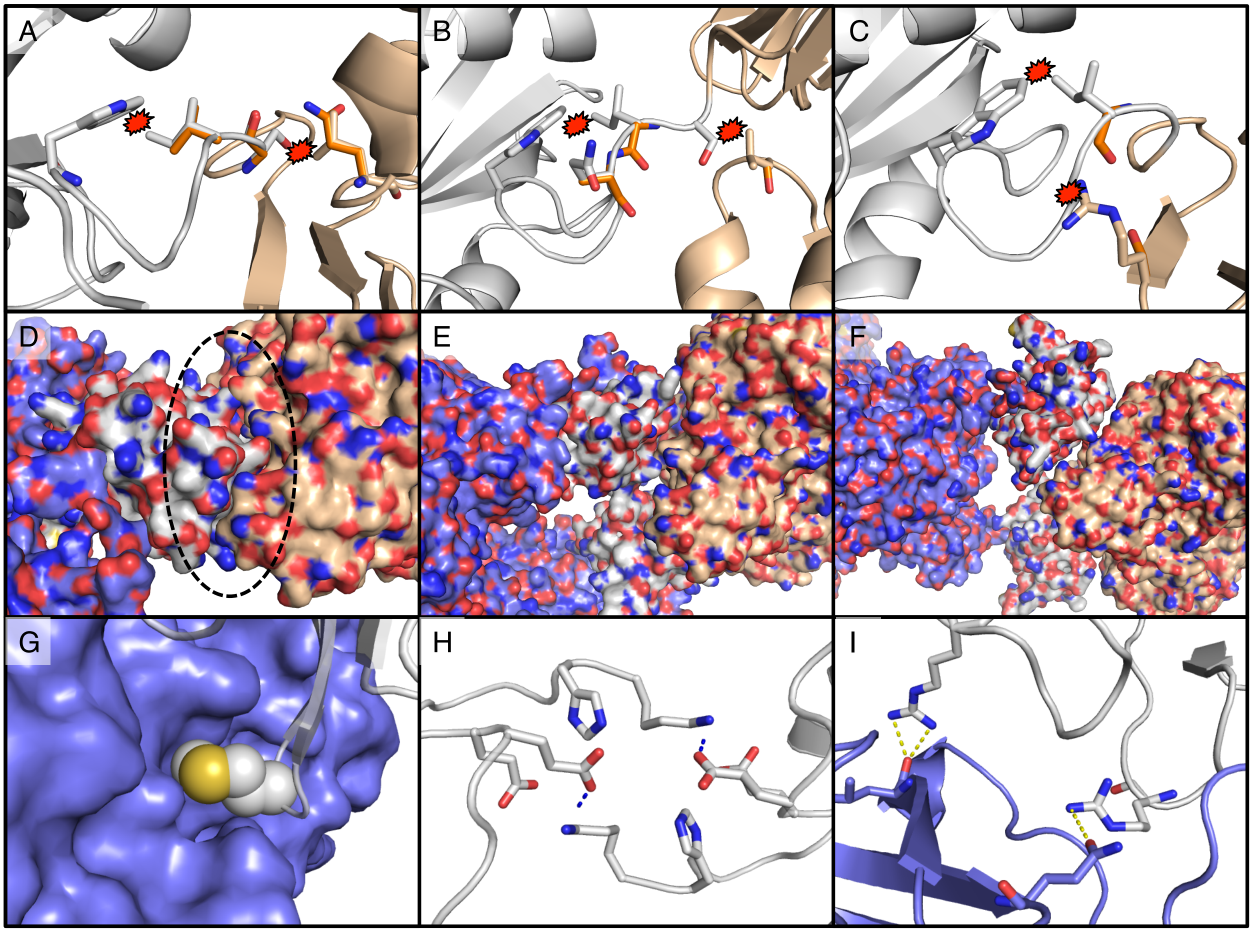


**Fig. S3. Design considerations for selecting substitutions and atomic interactions responsible for orientations of components during simulation.** Substitutions were introduced if they removed clashes (A-C), supported shape complementarity of novel interfaces (D-F), or created new favorable contacts across the components (G-I). (**A-C**) For all images, the AtzC monomer is colored in wheat, the SH2 is colored in white, and AtzA monomer colored in slate. Three different design models at unique core rotations: (A) AtzCM2 at 54.75°, (B) AtzCM3 at 54.75°, and (C) AtzCM5 at 0°. Clashes are denoted with a red star and are defined as heavy atoms within 3Å. (**D-F**) Novel interface complementarity considered in substitution selection. Substitutions that led to favorable shape complementarity and high surface area contact at the novel interfaces (D & E) were kept while design models that showed poor shape complementarity and low surface area contact were rejected (F). Shape complementarity along with novel hydrophobic (G), electrostatic (H), and polar contacts (I) created at novel interfaces are responsible for the decisions made during the stochastic fractal growth simulation.

**Table S1. List of substitutions and reasons for the various AtzA and AtzC designs.**

| | Substitution | Reason | Rotations (within 40 REU of lowest energy) |
|---|---|---|---|
| AtzAM1 | I515Y | Polarity | 0, 35.25, 54.75, 90, 125.25, 144.75, 180, |
| | E557S | Clash | |
| AtzAM2 | I516N | Polarity | 0, 125.25, 144.75 |
| | Q518G | Clash, flexibility | |
| | T519P | Clash, rigidity | |
| AtzAM3 | I516N | Polar contact | 0, 35.25, 54.75, 90, 125.25, 144.75, 180, 305.25 |
| | Q518G | Clash, flexibility | |
| | T519G | Clash, flexibility | |
| AtzAM4 | I516D | Electrostatic | 0, 35.25, 54.75, 90, 125.25, 144.75, 180, 305.26 |
| | Q518G | Clash, flexibility | |
| | T519G | Clash, flexibility | |
| AtzAM5 | I516D | Electrostatic | 0, 54.75, 90, 125.25, 144.75, 180, 305.27 |
| | Q518G | Clash, flexibility | |
| | R960H | Clash | |
| AtzCM1 | V402G | Clash, Flexibility | 0, 54.75, 90, 125.25, 144.75, 180 |
| | I403G | Clash, Flexibility | |
| | Q404S | Clash, H-bond to BB | |
| AtzCM2 | L148Q | H-bond to linker BB | 0, 35.25, 54.75, 90, 125.25, 144.75, 180, 324.75 |
| | V400G | Flexibility | |
| | S402G | Clash | |
| | I403V | Clash | |
| | Q404A | Clash | |
| AtzCM3 | K40G | Clash | 0, 54.75, 90, 125.25, 144.75, 180 |
| | L148S | Clash | |
| | V400I | Hydrophobic contact | |
| | I403G | Clash,Flexibility | |
| | Q404S | Hydrophobic contact | |
| AtzCM4 | R391A | Clash | 0, 54.75, 90, 125.25, 144.75, 180 |
| | V398Y | Polar contact | |
| | V400L | Hydrophobic contact | |
| | V402G | Clash, Flexibility | |
| | I403G | Clash, Flexibility | |
| | Q404S | Clash, H-bond to BB | |
| AtzCM5 | R391S | Clash, H-bond to linker BB | 0, 54.75, 90, 125.25, 144.75, 180 |
| | I393C | Clash | |
| | V398A | Clash | |
| | V400M | Clash | |
| | I403G | Clash, Flexibility | |
| | Q404S | Clash, H-bond to BB | |

Lowest energy = -3150 REU, Polarity = surface hydrophocity reduced, Clash = removed clashing sidechains, Flexibility = introduced backbone flexibility, Rigidity = reduced backbone flexibility, Polar Contact = introduced non-salt bridge side chain hydrogen bonds, Electrostatic = introduced a salt bridge, H-bond to BB = introduced a hydrogen bond to backbone atoms, H-bond to linker BB = introduced a hydrogen bond to the linker backbone atoms, Hydrophobic Contact = introduced a non-polar sidechain interaction

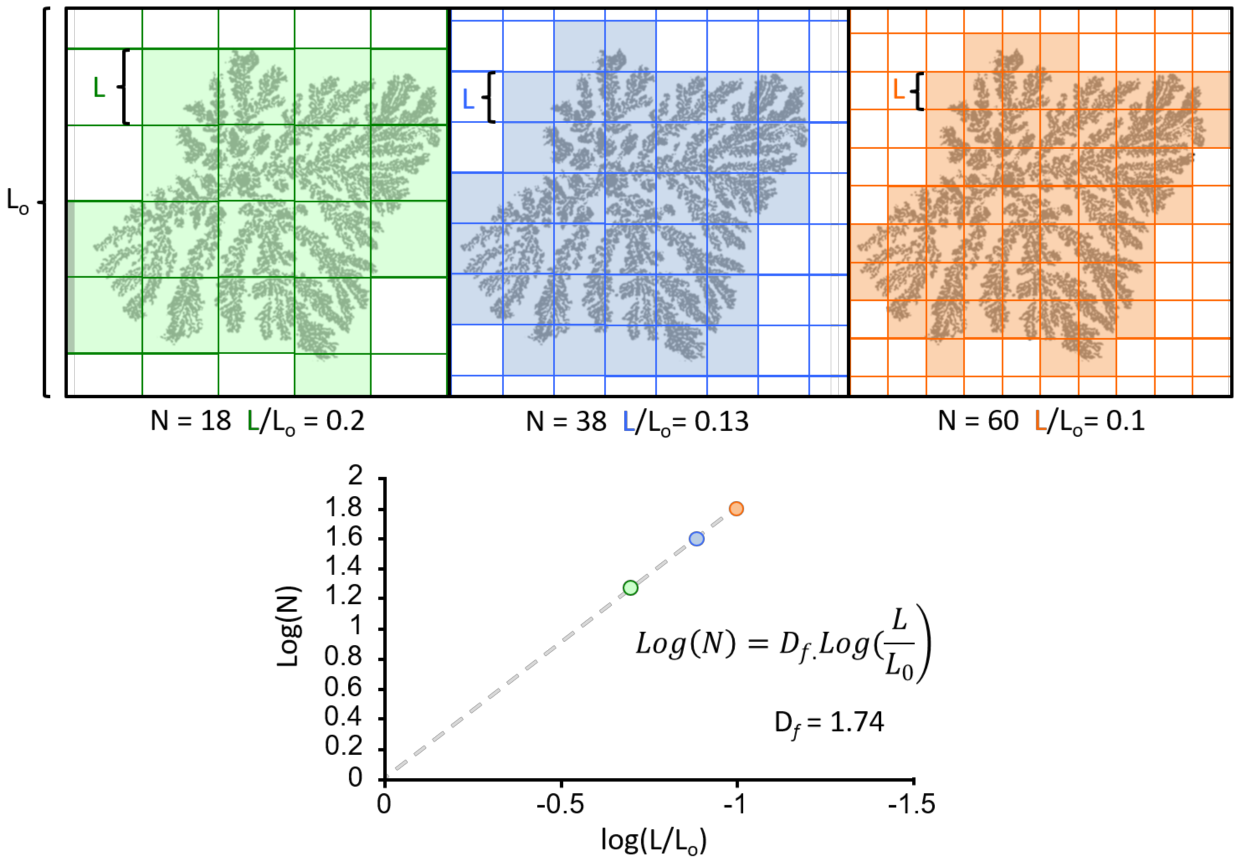


**Fig. S4. Illustration of the box-counting algorithm.** Analysis of a fractal image using the box counting method. As box size (L) is scaled down from left to right, the ratio of relative box size to the initial box decreases while the number of boxes (shaded boxes represent N) that contain the fractal (black image on white background) increases. The log/log relationship (slope) represents the fractal dimension ($D_f$ = 1.74). The individual colored points on the graph represent the respective colored image. Coarse features (large L) are represented closer to 0 while fine features (small L) are more negative. This illustration is just a conceptual representation of the actual method and is not an accurate representation of the fractal dimension. Programs like ImageJ will decrease L to nearly a pixel in length.

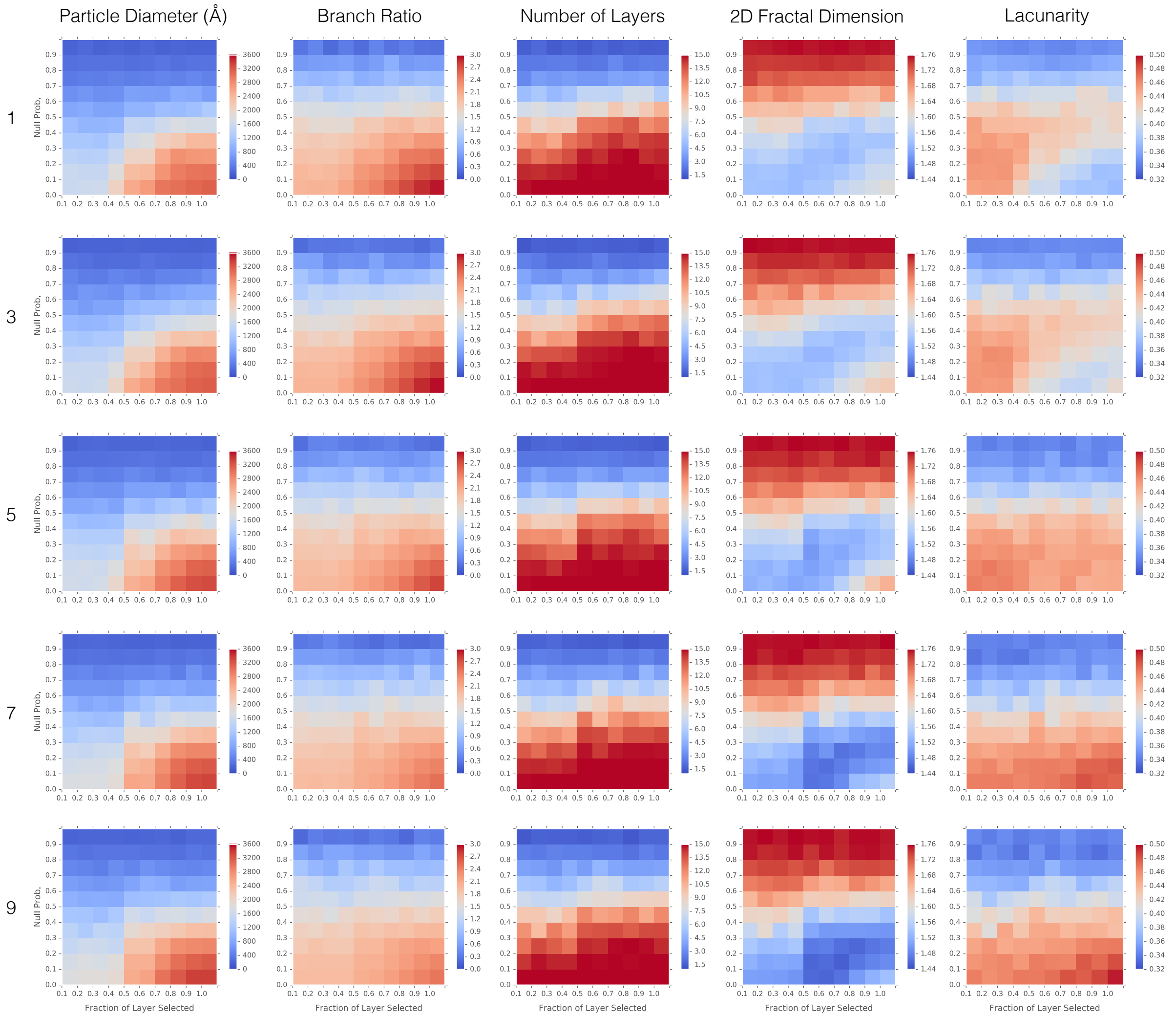


**Fig. S5.** Computational parameter sweep of *k*T (major y-axis), $P_{term}$ (minor y-axis), and $C_{frac}$ (minor x-axis). The various fractal topologies (limited to 15 layers) were evaluated by their particle diameter, branch ratio, layer count, 2D fractal dimension ($D_f$), and Lacunarity. We observe size, shape, and composition trends with varying $P_{term}$ and $C_{frac}$. Less obvious trends in topology via lacunarity and $D_f$ are also observed with changing *k*T. $P_{term}$ values above 0.4 (0.5-0.9) and $C_{frac}$ values below 0.5 (0.0-0.4) show a steep decline in particle size and number of total layers on average—terminating growth during simulation (unlike experimental data). For non-terminating values of $P_{term}$ (0.0-0.4) and $C_{frac}$ (0.5-1.0), $D_f$ is high (~1.7) when the connection probability is high—more isotropic fractal—and low (~1.6) when the connection probability is low—more anisotropic fractal shapes. When the *k*T increases we notice that the relative difference between high and low connection probability is maintained, however, the overall $D_f$ decreases (~1.6 and ~1.5) respectively. This can be attributed to the flatter probability landscape allowing for more 180° bound-angle (mixed vertex and edge centered connections around AtzA)—linearizing the branch connections on average and subsequently decreasing the fractal dimension.

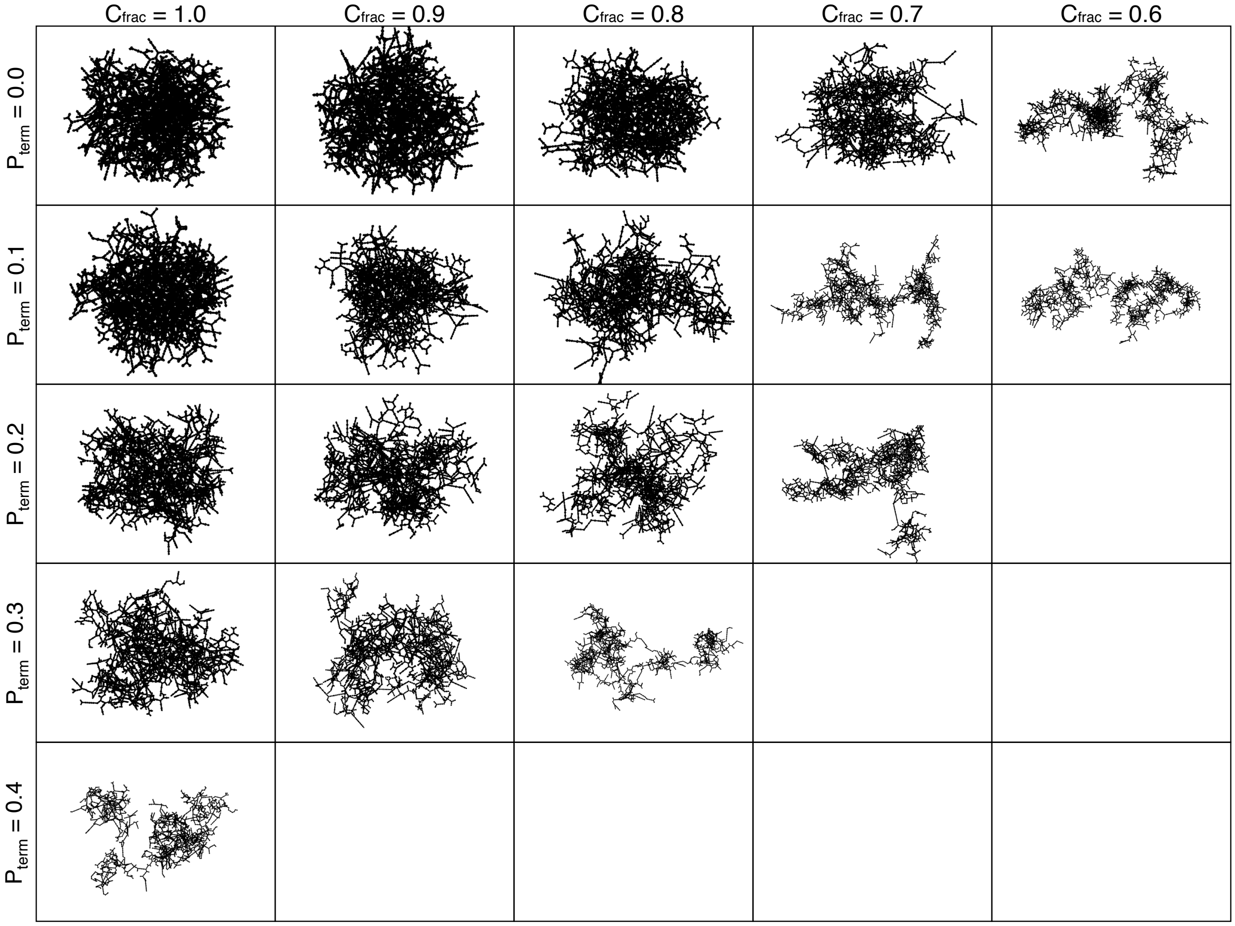


**Fig. S6.** Representative simulated fractal images (approx. 5000 components each and $kT = 9$) that possess the average layer count and branch ratio for varying values of $P_{term}$ (y-axis) and $C_{frac}$ (x-axis) of 100 models. As $C_{frac}$ decreases (or $P_{term}$ increases) fractal dimension decreases and lacunarity increases.

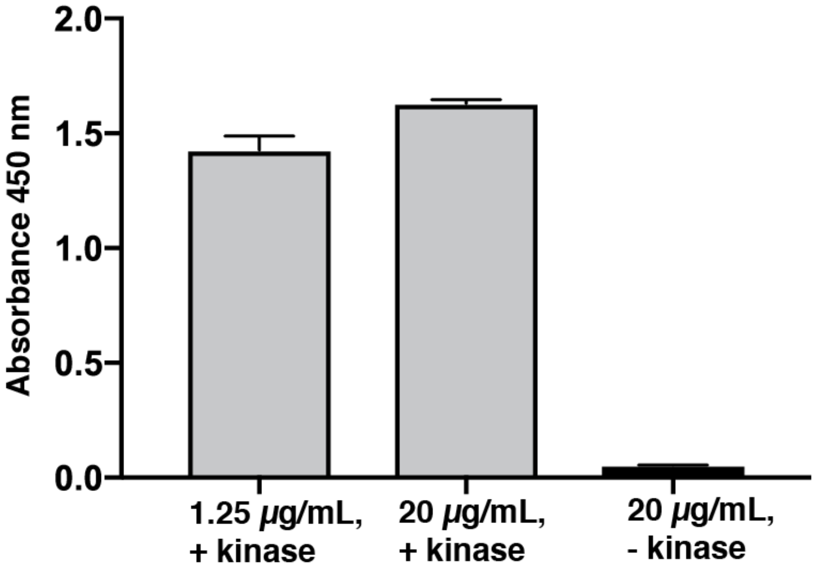


**Fig. S7. Phosphorylation of SH2 peptide AtzA fusion (pY-AtzA) by Src kinase.**
In order to verify phosphorylation of AtzA by Src kinase into phosphorylated SH2 peptide AtzA fusion (pY-AtzA), ELISA with (1:4000 dilution) antiphosphotyrosine-horseradish peroxidase conjugate was performed on pY-AtzA samples either with Src kinase (+) or without Src kinase (-), in phosphorylation reaction buffer at 1.25 μg/mL pY-AtzA or 20 μg/mL pY-AtzA. Data is presented as mean ± 1 standard deviation.

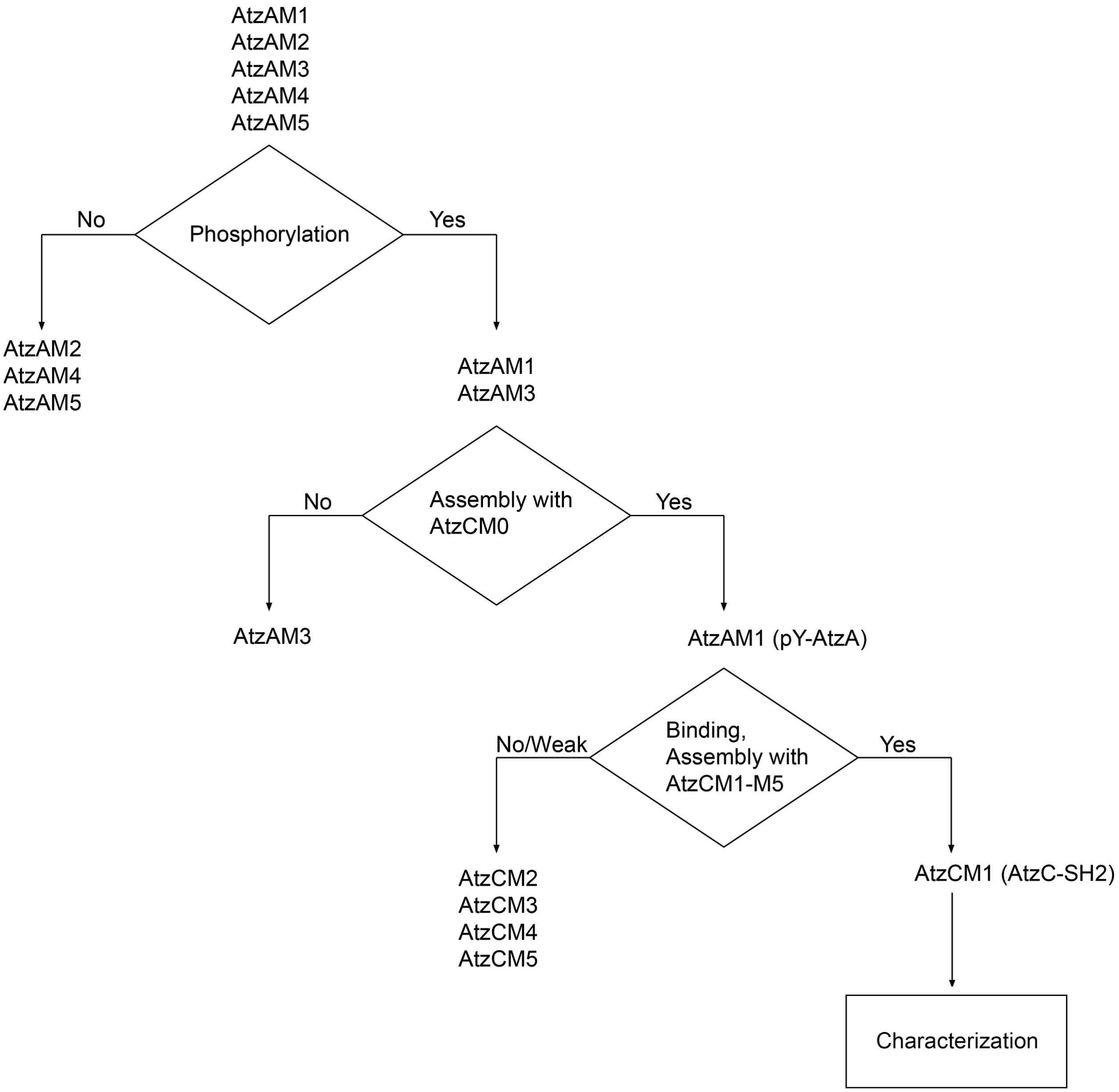


**Fig. S8. Experimental selection process for pY-AtzA and AtzC-SH2.** Five N-terminal SH2 binding peptide AtzA fusions (AtzAM1-AtzAM5) and five C-terminal SH2 binding domain AtzC fusions (AtzCM1-AtzCM5) were selected, cloned, expressed, and purified. AtzAM1-M5 were screened for having the ability to be phosphorylated via ELISA with anti-phosphotyrosine. Only two AtzA designs, AtzAM1 and AtzAM3, showed strong phosphorylation. The ability for assembly formation to occur with a direct C-terminal SH2 binding domain AtzC fusion (no mutations; AtzCM0) was used to select the best AtzA design. AtzAM1 was chosen for superior assembly formation ability, becoming pY-AtzA. The five AtzC designs AtzCM1-AtzCM5 were screened for the ability to effectively bind and assemble with pY-AtzA. The combination of pY-

AtzA and AtzCM1 (which we call AtzC-SH2) showed the strongest binding and the most robust assembly formation. This pair was then chosen for further characterization.

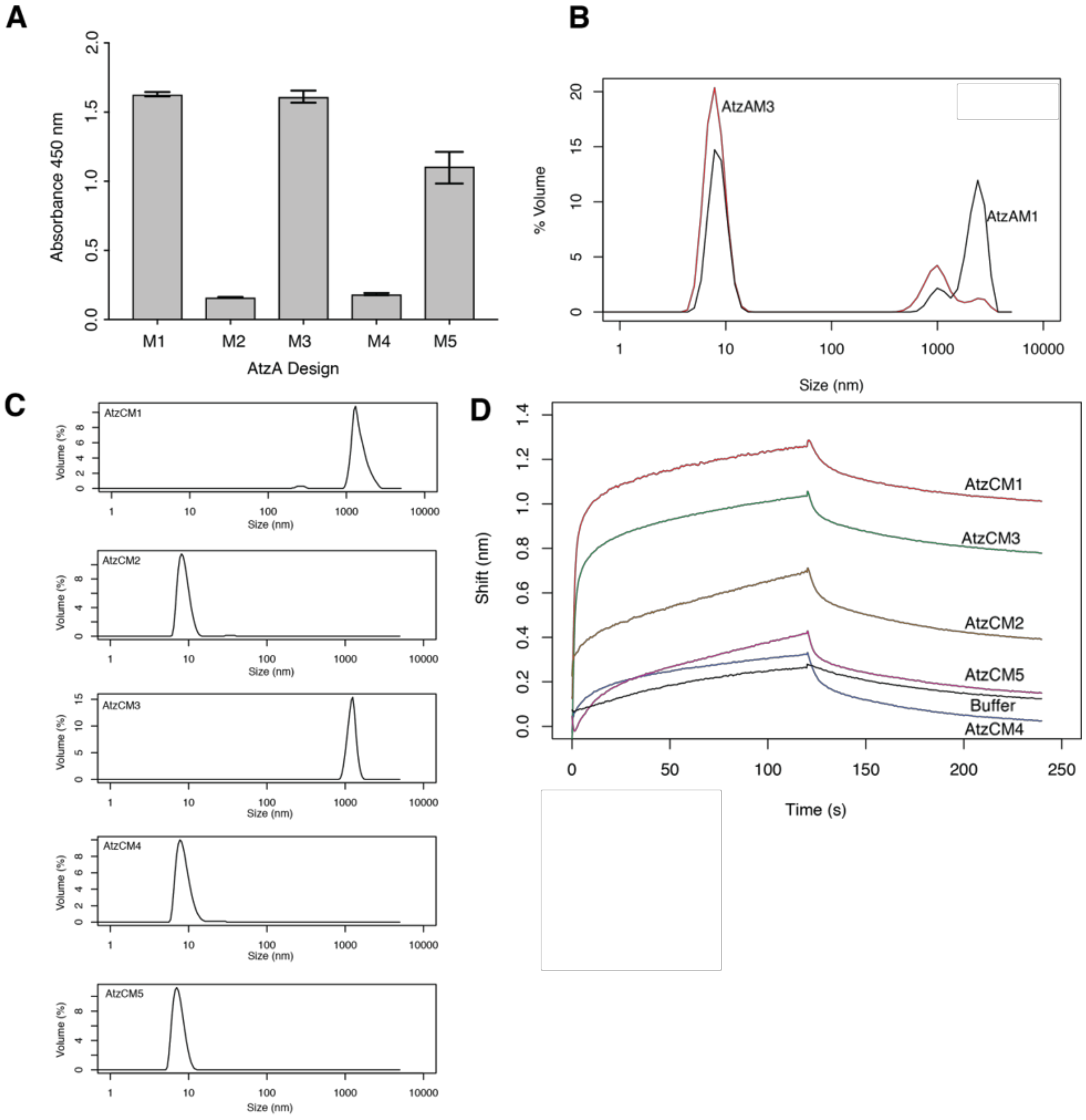


**Fig. S9. Experimental selection of AtzA, AtzC subunits for characterization.** (**A**) ELISA screening of AtzA designs to determine phosphorylation levels. (**B**) DLS size distribution of AtzA designs with AtzCM0. (**C**) DLS size distribution of AtzC-SH2 designs with pY-AtzA. Samples prepared at 3 µM pY-AtzA, 2 µM AtzC-SH2 design. Only AtzCM1 and AtzCM3 showed assembly formation with pY-AtzA. Volume distribution reported. (**D**) BLI binding traces of AtzC-SH2 designs with pY-AtzA. AtzC-SH2 designs were screened for binding with BLI, using pY-AtzA as the load. Out of all AtzC-SH2 designs prepared, AtzCM1 had the highest binding affinity to pY-AtzA. Based on the assembly formation and binding data, AtzCM1 was chosen for further investigation.

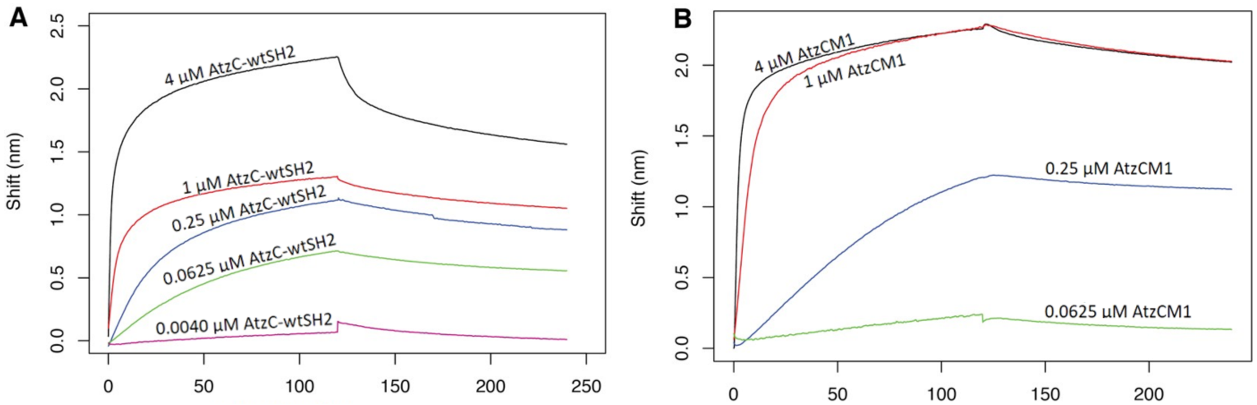


**Fig. S10. Biolayer interferometry (BLI) binding profiles of AtzC wildtype SH2 fusion (AtzC-wtSH2) and AtzC superbinder SH2 fusion (AtzC-SH2) to phosphorylated SH2 binding peptide AtzA fusion (pY-AtzA).** (**A**) Binding profile of AtzC-wtSH2 to pY-AtzA. PY-AtzA was loaded onto the biosensor via a streptavidin-biotin interaction. AtzC-wtSH2 was flowed into the sample. KD = 41.79 ± 0.32 nM. (**B**) Binding profile of AtzCM1 (superbinder) to pY-AtzA. PY-AtzA was loaded onto the biosensor via a streptavidin-biotin interaction. AtzC-SH2 was flowed into the sample. KD = 7.67 ± 0.52 nM.

```
AtzCM0  MSKDFDLIIRNAYLSEKDSVYDIGIVGDRIIKIEAKIEGTVKDEIDAKGNLVSPGFVDAH  60
AtzCM1  MSKDFDLIIRNAYLSEKDSVYDIGIVGDRIIKIEAKIEGTVKDEIDAKGNLVSPGFVDAH  60
AtzCM2  MSKDFDLIIRNAYLSEKDSVYDIGIVGDRIIKIEAKIEGTVKDEIDAKGNLVSPGFVDAH  60
AtzCM3  MSKDFDLIIRNAYLSEKDSVYDIGIVGDRIIKIEAKIEGTVGDEIDAKGNLVSPGFVDAH  60
AtzCM4  MSKDFDLIIRNAYLSEKDSVYDIGIVGDRIIKIEAKIEGTVKDEIDAKGNLVSPGFVDAH  60
AtzCM5  MSKDFDLIIRNAYLSEKDSVYDIGIVGDRIIKIEAKIEGTVKDEIDAKGNLVSPGFVDAH  60

AtzCM0  THMDKSFTSTGERLPKFWSRPYTRDAAIEDGLKYYKNATHEEIKRHVIEHAHMQVLHGTL  120
AtzCM1  THMDKSFTSTGERLPKFWSRPYTRDAAIEDGLKYYKNATHEEIKRHVIEHAHMQVLHGTL  120
AtzCM2  THMDKSFTSTGERLPKFWSRPYTRDAAIEDGLKYYKNATHEEIKRHVIEHAHMQVLHGTL  120
AtzCM3  THMDKSFTSTGERLPKFWSRPYTRDAAIEDGLKYYKNATHEEIKRHVIEHAHMQVLHGTL  120
AtzCM4  THMDKSFTSTGERLPKFWSRPYTRDAAIEDGLKYYKNATHEEIKRHVIEHAHMQVLHGTL  120
AtzCM5  THMDKSFTSTGERLPKFWSRPYTRDAAIEDGLKYYKNATHEEIKRHVIEHAHMQVLHGTL  120

AtzCM0  YTRTHVDVDSVAKTKAVEAVLEAKEELKDLIDIQVVAFAQSGFFVDLESESLIRKSLDMG  180
AtzCM1  YTRTHVDVDSVAKTKAVEAVLEAKEELKDLIDIQVVAFAQSGFFVDLESESLIRKSLDMG  180
AtzCM2  YTRTHVDVDSVAKTKAVEAVLEAKEELKDQIDIQVVAFAQSGFFVDLESESLIRKSLDMG  180
AtzCM3  YTRTHVDVDSVAKTKAVEAVLEAKEELKDSIDIQVVAFAQSGFFVDLESESLIRKSLDMG  180
AtzCM4  YTRTHVDVDSVAKTKAVEAVLEAKEELKDLIDIQVVAFAQSGFFVDLESESLIRKSLDMG  180
AtzCM5  YTRTHVDVDSVAKTKAVEAVLEAKEELKDLIDIQVVAFAQSGFFVDLESESLIRKSLDMG  180

AtzCM0  CDLVGGVDPATRENNVEGSLDLCFKLAKEYDVDIDYHIHDIGTVGVYSINRLAQKTIENG  240
AtzCM1  CDLVGGVDPATRENNVEGSLDLCFKLAKEYDVDIDYHIHDIGTVGVYSINRLAQKTIENG  240
AtzCM2  CDLVGGVDPATRENNVEGSLDLCFKLAKEYDVDIDYHIHDIGTVGVYSINRLAQKTIENG  240
AtzCM3  CDLVGGVDPATRENNVEGSLDLCFKLAKEYDVDIDYHIHDIGTVGVYSINRLAQKTIENG  240
AtzCM4  CDLVGGVDPATRENNVEGSLDLCFKLAKEYDVDIDYHIHDIGTVGVYSINRLAQKTIENG  240
AtzCM5  CDLVGGVDPATRENNVEGSLDLCFKLAKEYDVDIDYHIHDIGTVGVYSINRLAQKTIENG  240

AtzCM0  YKGRVTTSHAWCFADAPSEWLDEAIPLYKDSGMKFVTCFSSTPPTMPVIKLLEAGINLGC  300
AtzCM1  YKGRVTTSHAWCFADAPSEWLDEAIPLYKDSGMKFVTCFSSTPPTMPVIKLLEAGINLGC  300
AtzCM2  YKGRVTTSHAWCFADAPSEWLDEAIPLYKDSGMKFVTCFSSTPPTMPVIKLLEAGINLGC  300
AtzCM3  YKGRVTTSHAWCFADAPSEWLDEAIPLYKDSGMKFVTCFSSTPPTMPVIKLLEAGINLGC  300
AtzCM4  YKGRVTTSHAWCFADAPSEWLDEAIPLYKDSGMKFVTCFSSTPPTMPVIKLLEAGINLGC  300
AtzCM5  YKGRVTTSHAWCFADAPSEWLDEAIPLYKDSGMKFVTCFSSTPPTMPVIKLLEAGINLGC  300

AtzCM0  ASDNIRDFWVPFGNGDMVQGALIETQRLELKTNRDLGLIWKMITSEGARVLGIEKNYGIE  360
AtzCM1  ASDNIRDFWVPFGNGDMVQGALIETQRLELKTNRDLGLIWKMITSEGARVLGIEKNYGIE  360
AtzCM2  ASDNIRDFWVPFGNGDMVQGALIETQRLELKTNRDLGLIWKMITSEGARVLGIEKNYGIE  360
AtzCM3  ASDNIRDFWVPFGNGDMVQGALIETQRLELKTNRDLGLIWKMITSEGARVLGIEKNYGIE  360
AtzCM4  ASDNIRDFWVPFGNGDMVQGALIETQRLELKTNRDLGLIWKMITSEGARVLGIEKNYGIE  360
AtzCM5  ASDNIRDFWVPFGNGDMVQGALIETQRLELKTNRDLGLIWKMITSEGARVLGIEKNYGIE  360

AtzCM0  VGKKADLVVLNSLSPQWAIIDQAKRLCVIKNGRIIVKDEVIVASIQAEEWYFGKLGRKDA  420
AtzCM1  VGKKADLVVLNSLSPQWAIIDQAKRLCVIKNGRIIVKDEVIVAGGSAEEWYFGKLGRKDA  420
AtzCM2  VGKKADLVVLNSLSPQWAIIDQAKRLCVIKNGRIIVKDEVIGAGVAAEEWYFGKLGRKDA  420
AtzCM3  VGKKADLVVLNSLSPQWAIIDQAKRLCVIKNGRIIVKDEVIIASGAAEEWYFGKLGRKDA  420
AtzCM4  VGKKADLVVLNSLSPQWAIIDQAKRLCVIKNGAIIVKDEYILAGGSAEEWYFGKLGRKDA  420
AtzCM5  VGKKADLVVLNSLSPQWAIIDQAKRLCVIKNGSICVKDEAIMASGSAEEWYFGKLGRKDA  420
```

**Fig. S11a.** Sequence alignment of AtzC-SH2 designs AtzCM0-AtzCM5.

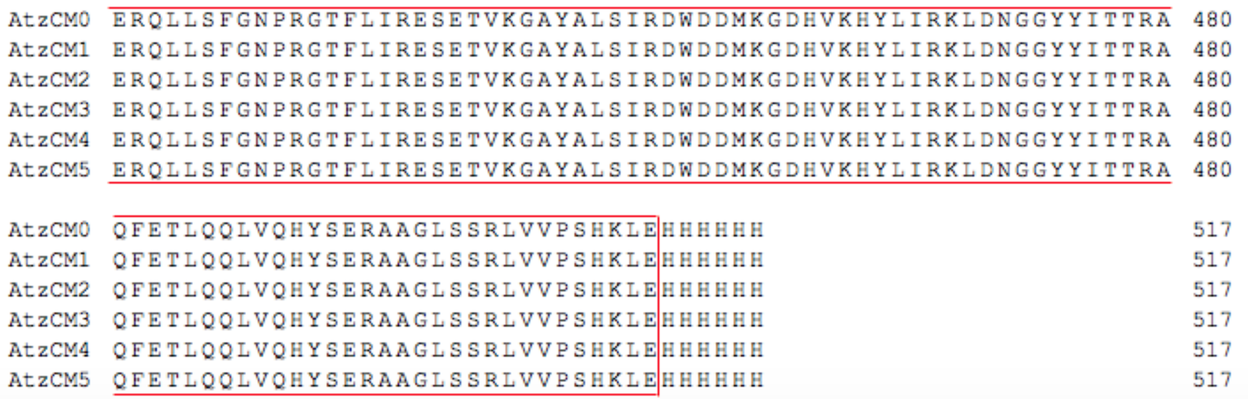


**Fig. S11b. Sequence alignment of AtzC-SH2 designs AtzCM0-AtzCM5 (con’t).** Sequence alignment of AtzC-SH2 designs prepared. AtzCM0 is a direct fusion of AtzC and superbinder SH2 domain without mutations. Mutations made are highlighted in black or grey (similar residues). The red box highlights the region where the superbinder SH2 domain is located.

```
AtzAM0      MGSSHHHHHHSSGLVPRGSHMEPQYEEIPIYQTLSIQHGTLVTMDQYRRVLGDSWVHVQD    60
AtzAM1      MGSSHHHHHHSSGLVPRGSHMEPQYEEIPYYQTLSIQHGTLVTMDQYRRVLGDSWVHVQD    60
AtzAM2      MGSSHHHHHHSSGLVPRGSHMEPQYEEIPNYGPLSIQHGTLVTMDQYRRVLGDSWVHVQD    60
AtzAM3      MGSSHHHHHHSSGLVPRGSHMEPQYEEIPNYGGLSIQHGTLVTMDQYRRVLGDSWVHVQD    60
AtzAM4      MGSSHHHHHHSSGLVPRGSHMEPQYEEIPDYGGLSIQHGTLVTMDQYRRVLGDSWVHVQD    60
AtzAM5      MGSSHHHHHHSSGLVPRGSHMEPQYEEIPDYGTLSIQHGTLVTMDQYRRVLGDSWVHVQD    60

AtzAM0      GRIVALGVHAESVPPPADRVIDARGKVVLPGFINAHTHVNQILLRGGPSHGRQFYDWLFN    120
AtzAM1      GRIVALGVHASSVPPPADRVIDARGKVVLPGFINAHTHVNQILLRGGPSHGRQFYDWLFN    120
AtzAM2      GRIVALGVHAESVPPPADRVIDARGKVVLPGFINAHTHVNQILLRGGPSHGRQFYDWLFN    120
AtzAM3      GRIVALGVHAESVPPPADRVIDARGKVVLPGFINAHTHVNQILLRGGPSHGRQFYDWLFN    120
AtzAM4      GRIVALGVHAESVPPPADRVIDARGKVVLPGFINAHTHVNQILLRGGPSHGRQFYDWLFN    120
AtzAM5      GRIVALGVHAESVPPPADRVIDARGKVVLPGFINAHTHVNQILLRGGPSHGRQFYDWLFN    120

AtzAM0      VVYPGQKAMRPEDVAVAVRLYCAEAVRSGITTINENADSAIYPGNIEAAMAVYGEVGVRV    180
AtzAM1      VVYPGQKAMRPEDVAVAVRLYCAEAVRSGITTINENADSAIYPGNIEAAMAVYGEVGVRV    180
AtzAM2      VVYPGQKAMRPEDVAVAVRLYCAEAVRSGITTINENADSAIYPGNIEAAMAVYGEVGVRV    180
AtzAM3      VVYPGQKAMRPEDVAVAVRLYCAEAVRSGITTINENADSAIYPGNIEAAMAVYGEVGVRV    180
AtzAM4      VVYPGQKAMRPEDVAVAVRLYCAEAVRSGITTINENADSAIYPGNIEAAMAVYGEVGVRV    180
AtzAM5      VVYPGQKAMRPEDVAVAVRLYCAEAVRSGITTINENADSAIYPGNIEAAMAVYGEVGVRV    180

AtzAM0      VYARMFFDRMDGRIQGYVDALKARSPQVELCSIMEETAVAKDRITALSDQYHGTAGGRIS    240
AtzAM1      VYARMFFDRMDGRIQGYVDALKARSPQVELCSIMEETAVAKDRITALSDQYHGTAGGRIS    240
AtzAM2      VYARMFFDRMDGRIQGYVDALKARSPQVELCSIMEETAVAKDRITALSDQYHGTAGGRIS    240
AtzAM3      VYARMFFDRMDGRIQGYVDALKARSPQVELCSIMEETAVAKDRITALSDQYHGTAGGRIS    240
AtzAM4      VYARMFFDRMDGRIQGYVDALKARSPQVELCSIMEETAVAKDRITALSDQYHGTAGGRIS    240
AtzAM5      VYARMFFDRMDGRIQGYVDALKARSPQVELCSIMEETAVAKDRITALSDQYHGTAGGRIS    240

AtzAM0      VWPAPATTTAVTVEGMRWAQAFARDRAVMWTLHMAESDHDERIHGMSPAEYMECYGLLDE    300
AtzAM1      VWPAPATTTAVTVEGMRWAQAFARDRAVMWTLHMAESDHDERIHGMSPAEYMECYGLLDE    300
AtzAM2      VWPAPATTTAVTVEGMRWAQAFARDRAVMWTLHMAESDHDERIHGMSPAEYMECYGLLDE    300
AtzAM3      VWPAPATTTAVTVEGMRWAQAFARDRAVMWTLHMAESDHDERIHGMSPAEYMECYGLLDE    300
AtzAM4      VWPAPATTTAVTVEGMRWAQAFARDRAVMWTLHMAESDHDERIHGMSPAEYMECYGLLDE    300
AtzAM5      VWPAPATTTAVTVEGMRWAQAFARDRAVMWTLHMAESDHDERIHGMSPAEYMECYGLLDE    300

AtzAM0      RLQVAHCVYFDRKDVRLLHRHNVKVASQVVSNAYLGSGVAPVPEMVERGMAVGIGTDNGN    360
AtzAM1      RLQVAHCVYFDRKDVRLLHRHNVKVASQVVSNAYLGSGVAPVPEMVERGMAVGIGTDNGN    360
AtzAM2      RLQVAHCVYFDRKDVRLLHRHNVKVASQVVSNAYLGSGVAPVPEMVERGMAVGIGTDNGN    360
AtzAM3      RLQVAHCVYFDRKDVRLLHRHNVKVASQVVSNAYLGSGVAPVPEMVERGMAVGIGTDNGN    360
AtzAM4      RLQVAHCVYFDRKDVRLLHRHNVKVASQVVSNAYLGSGVAPVPEMVERGMAVGIGTDNGN    360
AtzAM5      RLQVAHCVYFDRKDVRLLHRHNVKVASQVVSNAYLGSGVAPVPEMVERGMAVGIGTDNGN    360

AtzAM0      SNDSVNMIGDMKFMAHIHRAVHRDADVLTPEKILEMATIDGARSLGMDHEIGSIETGKRA    420
AtzAM1      SNDSVNMIGDMKFMAHIHRAVHRDADVLTPEKILEMATIDGARSLGMDHEIGSIETGKRA    420
AtzAM2      SNDSVNMIGDMKFMAHIHRAVHRDADVLTPEKILEMATIDGARSLGMDHEIGSIETGKRA    420
AtzAM3      SNDSVNMIGDMKFMAHIHRAVHRDADVLTPEKILEMATIDGARSLGMDHEIGSIETGKRA    420
AtzAM4      SNDSVNMIGDMKFMAHIHRAVHRDADVLTPEKILEMATIDGARSLGMDHEIGSIETGKRA    420
AtzAM5      SNDSVNMIGDMKFMAHIHRAVHRDADVLTPEKILEMATIDGARSLGMDHEIGSIETGKRA    420

AtzAM0      DLILLDLRHPQTTPHHHLAATIVFQAYGNEVDTVLIDGNVVMENRRLSFLPPERELAFLE    480
AtzAM1      DLILLDLRHPQTTPHHHLAATIVFQAYGNEVDTVLIDGNVVMENRRLSFLPPERELAFLE    480
AtzAM2      DLILLDLRHPQTTPHHHLAATIVFQAYGNEVDTVLIDGNVVMENRRLSFLPPERELAFLE    480
AtzAM3      DLILLDLRHPQTTPHHHLAATIVFQAYGNEVDTVLIDGNVVMENRRLSFLPPERELAFLE    480
AtzAM4      DLILLDLRHPQTTPHHHLAATIVFQAYGNEVDTVLIDGNVVMENRRLSFLPPERELAFLE    480
AtzAM5      DLILLDLRHPQTTPHHHLAATIVFQAYGNEVDTVLIDGNVVMENRRLSFLPPEHELAFLE    480

AtzAM0      EAQSRATAILQRANMVANPAWRSL        504
AtzAM1      EAQSRATAILQRANMVANPAWRSL        504
AtzAM2      EAQSRATAILQRANMVANPAWRSL        504
AtzAM3      EAQSRATAILQRANMVANPAWRSL        504
AtzAM4      EAQSRATAILQRANMVANPAWRSL        504
AtzAM5      EAQSRATAILQRANMVANPAWRSL        504
```

**Fig. S12. Sequence alignment of pY-AtzA designs.**
Sequence alignment of pY-AtzA designs prepared. AtzAM0 is a direct fusion of AtzA and SH2 binding peptide without mutations. Mutations made are shown in black. The red box indicates the SH2 recognition peptide sequence.

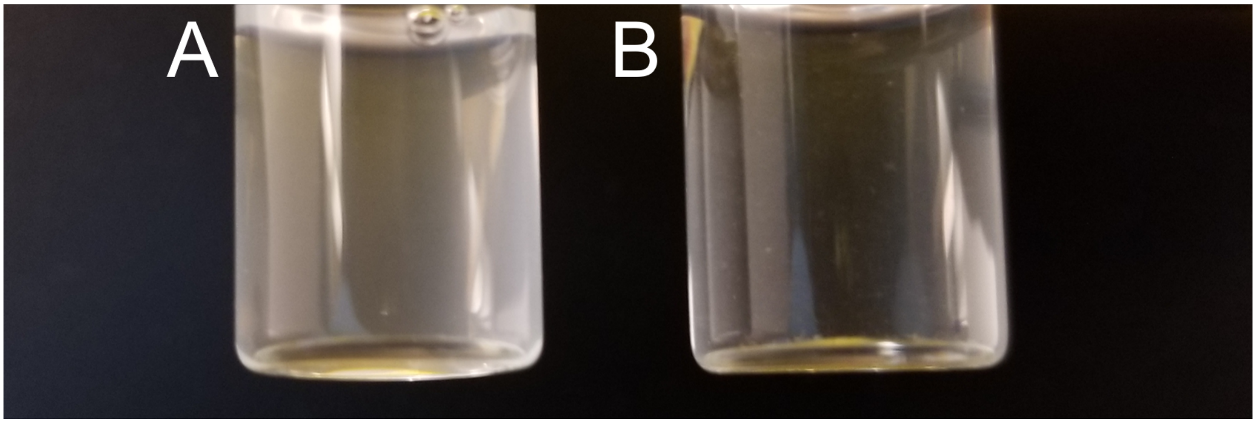


**Fig. S13.** (**A)** 3 μM pY-AtzAM1 and 2 μM AtzCM1, shows a turbid solution that represents the assembly formed. (**B**) 3 μM non-pY-AtzAM1 and 2 μM AtzCM1, shows a clear solution with no assembly formation.

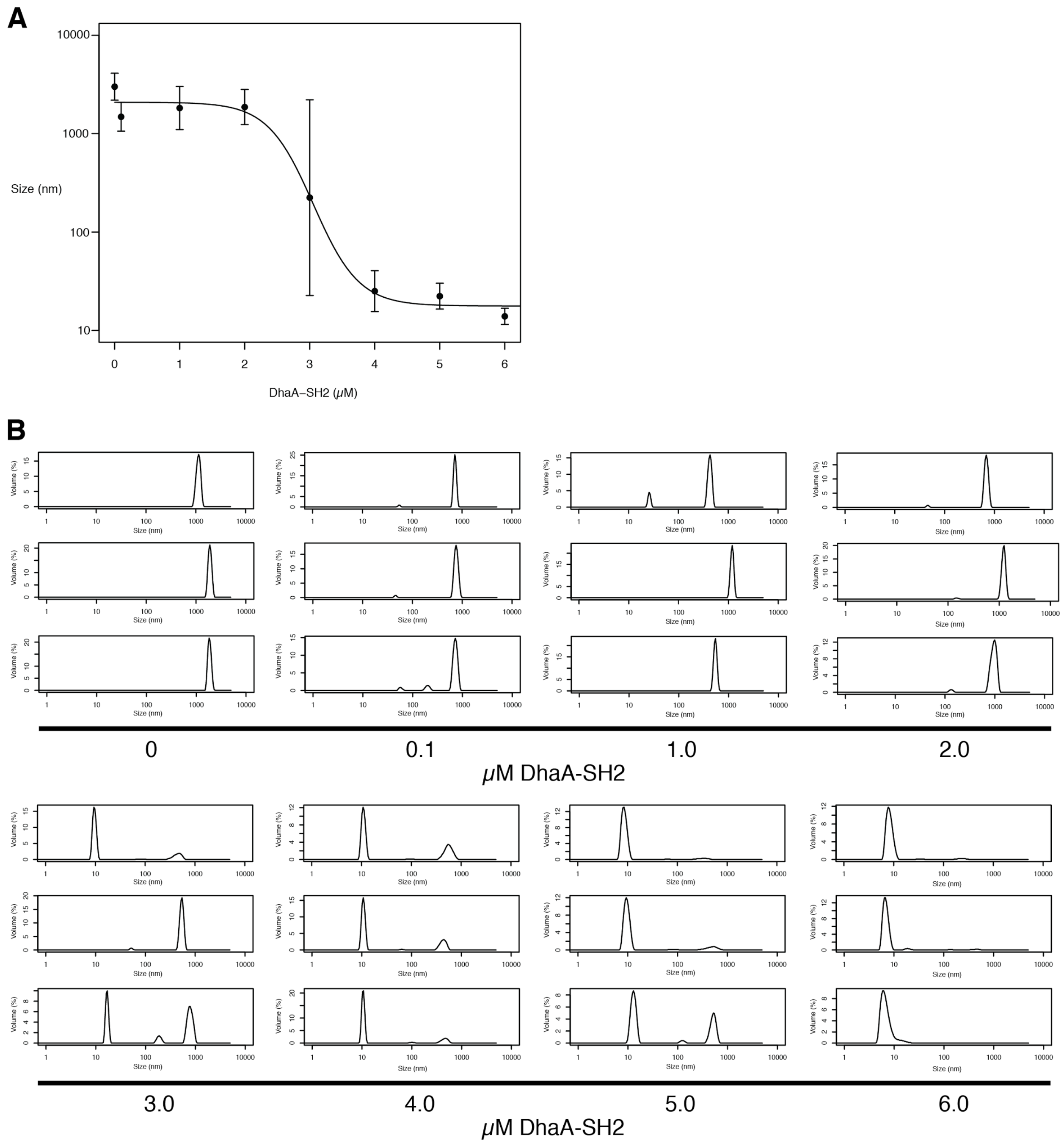


**Fig. S14. Inhibition of assembly at 0.66 μM AtzC-SH2, 1 μM pY-AtzA, 0-6 μM SH2-DhaA.** (**A**) **Inhibition graph of SH2-DhaA on 0.66 μM AtzC-SH2, 1 μM pY-AtzA assembly.** Size recorded represents most predominant DLS sizing peak. Data are presented as mean ± 1 standard deviation. IC50 = 3.05 μM. Adjusted $R^2$ = 0.98. (**B**) DLS traces of assembly from 0 - 6 μM SH2-DhaA. DLS traces are of triplicates.

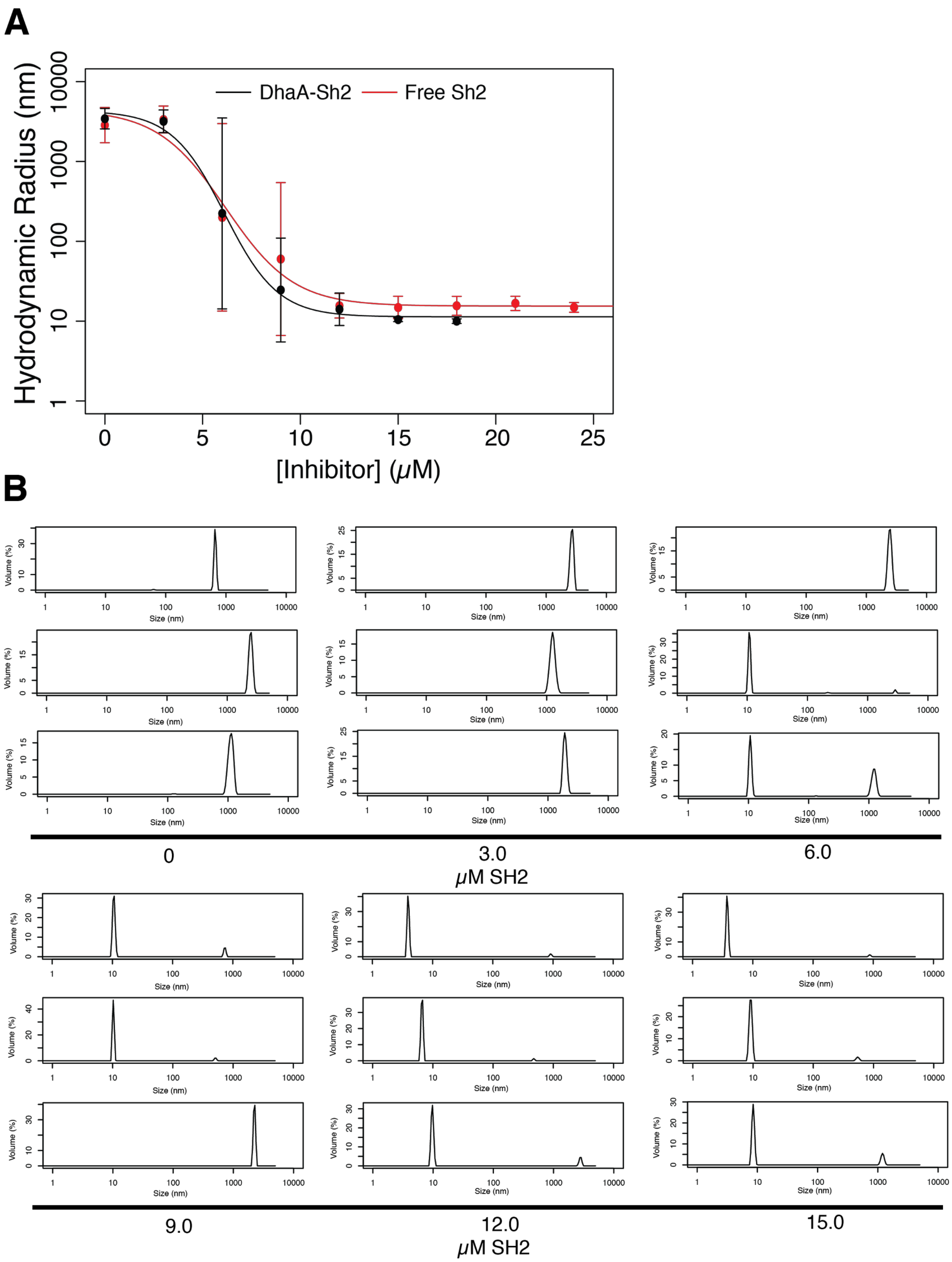


**Fig. S15. Inhibition of assembly at 2 μM AtzC-SH2, 3 μM pY-AtzA with 0-15 μM inhibitor.**

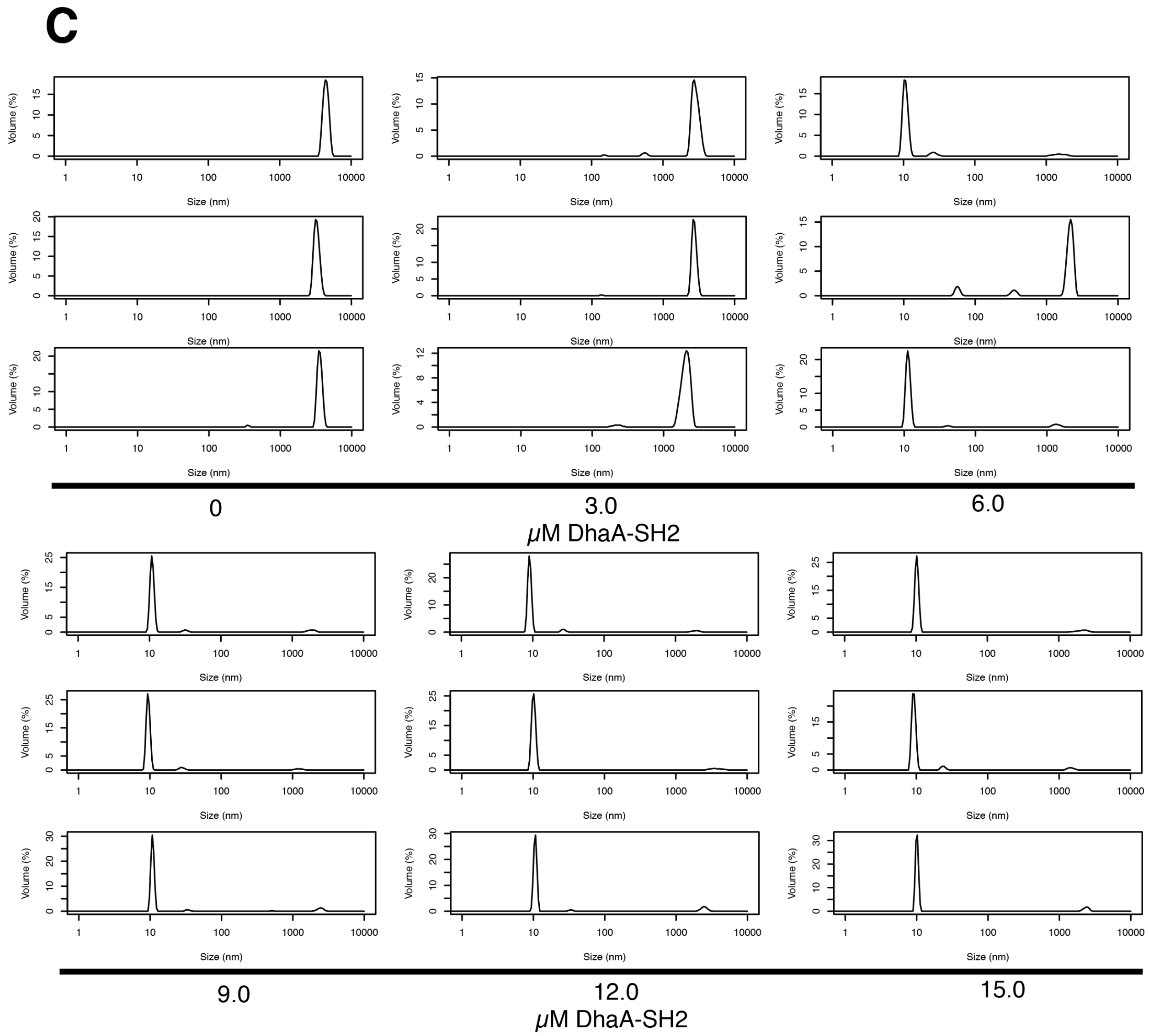


**Fig. S16. Inhibition of assembly at 2 µM AtzC-SH2, 3 µM pY-AtzA with 0-15 µM inhibitor (con't).**

All DLS traces were performed in triplicate (**A**) Inhibition graph of SH2-DhaA of 2 µM AtzC-SH2, 3 µM pY-AtzA assembly. Size recorded represents most predominant DLS sizing peak. Data are presented as mean ± 1 standard deviation. IC50 (SH2) = 6.18 µM, IC50 (SH2-DhaA) = 6.13 µM. Adjusted $R^2$ (SH2) = 0.97. Adjusted $R^2$ (SH2-DhaA) = 0.99. (**B**) DLS traces of assembly from 0-15 µM SH2. (**C**) DLS traces of assembly from 0-15 µM SH2-DhaA.

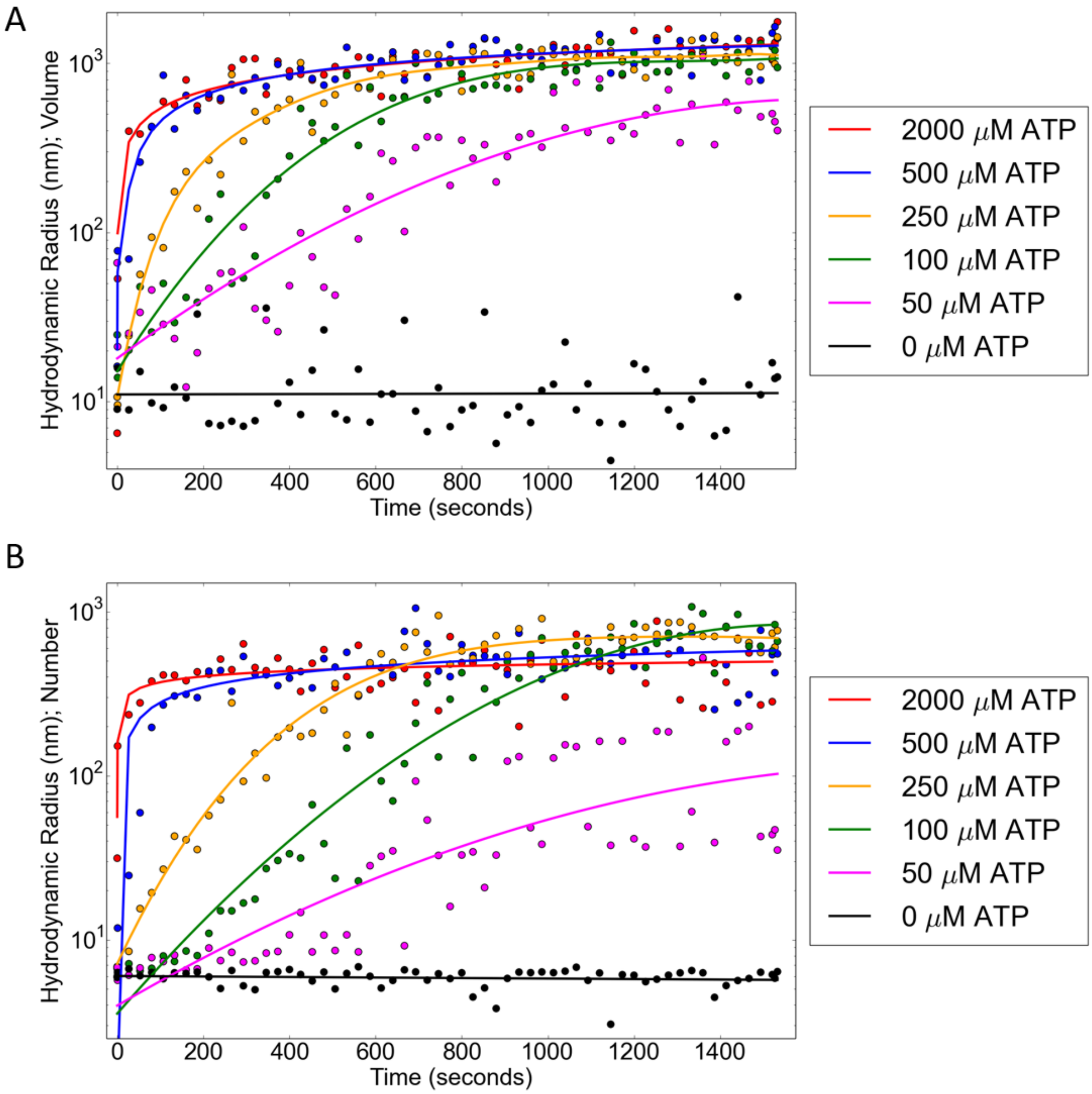


**Fig. S17. Rate of assembly formation is dependent on ATP concentration.** (**A**) Volume mean of sample from 0 – 1500 sec. Each point represents average of triplicates. (**B**) Number mean of sample from 0 – 1500 sec. Each point represents average of triplicates. Curve fitting performed using sloping spline with smoothness parameter (p) and adjusted $R^2$ value given in Table S1 (highest concentration of ATP to lowest, starting from top to bottom at time 0 for both graphs).

**Table S2. Curve fitting data for Figure S16.** Adjusted $R^2$ and smoothing parameter (p) value given for curve fitting done on assembly kinetics data.

| Distribution | ATP µM | Adjusted R-square | *p* |
|---|---|---|---|
| Vol | 2000 | 0.7889 | 1.31E-05 |
| Vol | 500 | 0.888 | 1.31E-05 |
| Vol | 250 | 0.898 | 3.25E-08 |
| Vol | 100 | 0.9374 | 3.25E-08 |
| Vol | 50 | 0.867 | 3.25E-08 |
| Vol | 0 | 0.2638 | 0.000182922 |
| Num | 2000 | 0.5044 | 2.16E-05 |
| Num | 500 | 0.9303 | 2.16E-05 |
| Num | 250 | 0.9678 | 2.16E-05 |
| Num | 100 | 0.9543 | 2.16E-05 |
| Num | 50 | 0.7189 | 7.25E-09 |
| Num | 0 | 0.3338 | 0.000110956 |

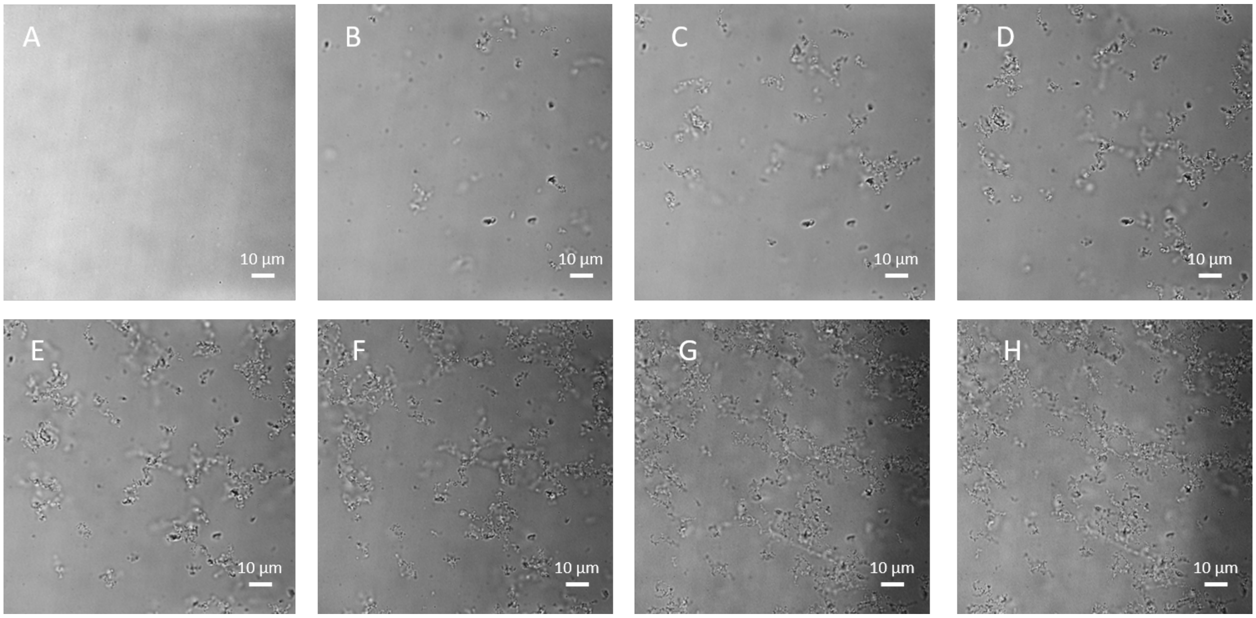


**Fig. S18. Bright-field view of the assembly growing after the addition of Src kinase.** (**A**) 3 minutes after addition of Src kinase, no assemblies shown. (**B**) 14 minutes after addition of Src kinase, small assemblies shown. (**C**) 18 minutes after addition of Src kinase, small 10 µm assemblies start to grow (**D**) 24 minutes after the addition of Src kinase, growth continues. (**E**) 30 minutes after addition of Src kinase, over 50 µm size assemblies form. (**F**) 35 minutes after addition of Src kinase, 100 µm size assemblies appear. (**G**) 40 minutes after addition of Src kinase, assemblies continue to grow. (**H**) 50 minutes after addition of Src kinase, assemblies have fully matured into fractal-like structures.

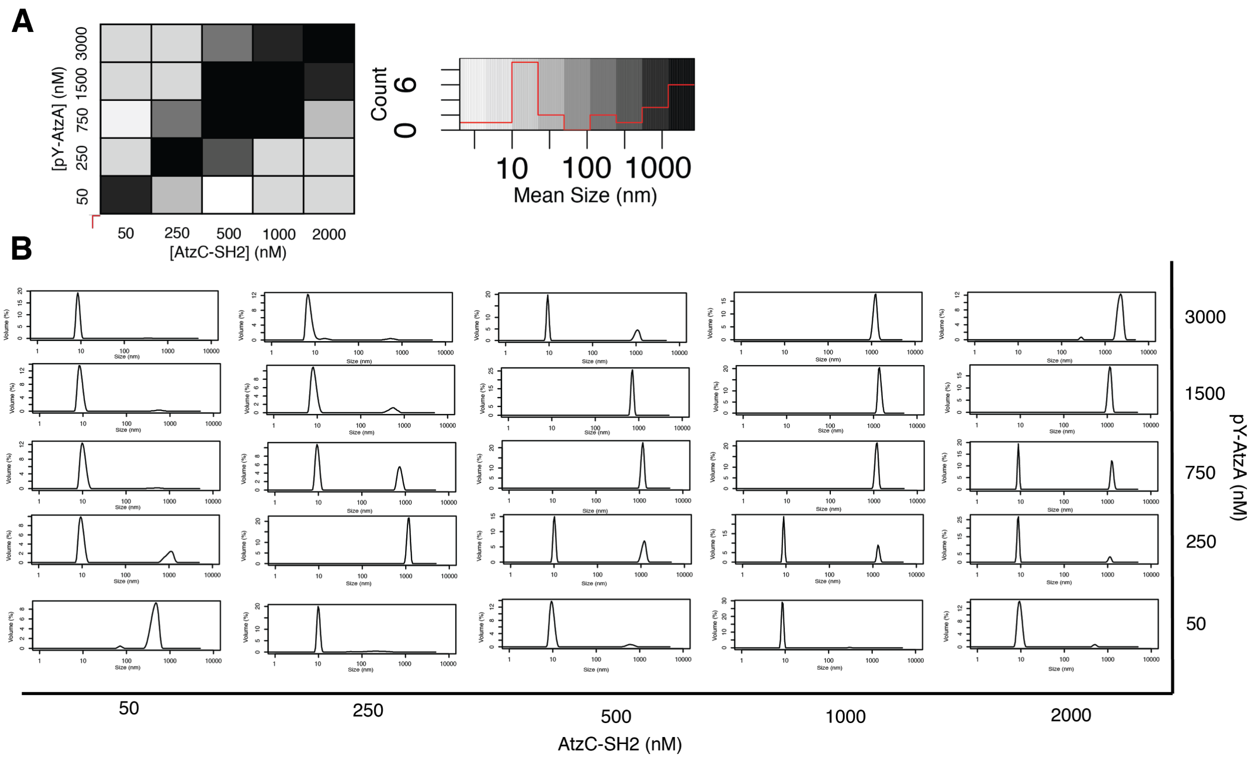


**Fig. S19. Average size of particle formed by pY-AtzA and wild type AtzC-SH2.** (**A**) Heat map showing volume-weighted mean size of particles found from 50-3000 nM pY-AtzA and 50-2000 nM AtzC-SH2. Value shown is average of two physical samples. Histogram illustrates distribution of sizes found on heatmap. (**B**) Volume distributions of heat map. Distributions shown are representative of other traces in the sample.

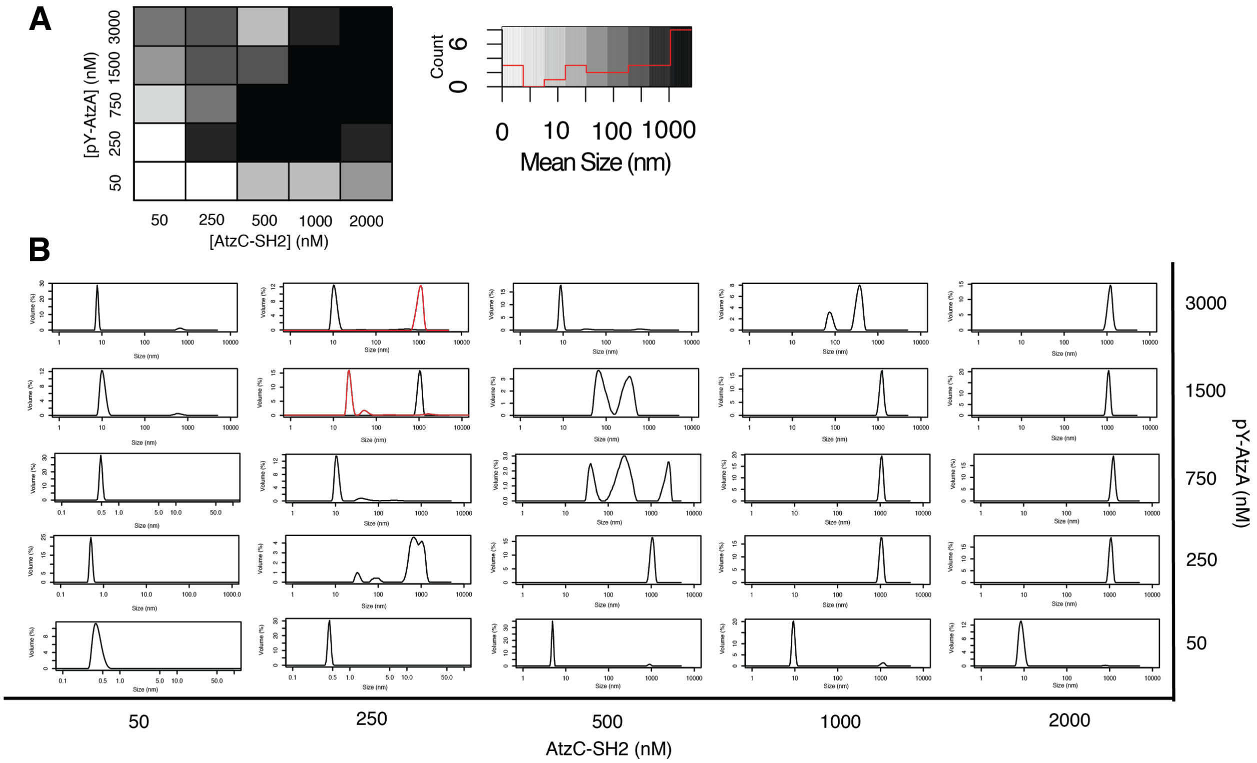


**Fig. S20. Average size of particle formed by pY-AtzA and super-binder AtzC-SH2.** (**A**) Heat map showing volume-weighted mean size of particles found from 50-3000 nM pY-AtzA and 50-2000 nM AtzC-SH2. Value shown is average of two physical samples. Histogram illustrates distribution of sizes found on heatmap. (**B**) Volume distributions of heat map. Distributions shown are representative of other traces in the sample.

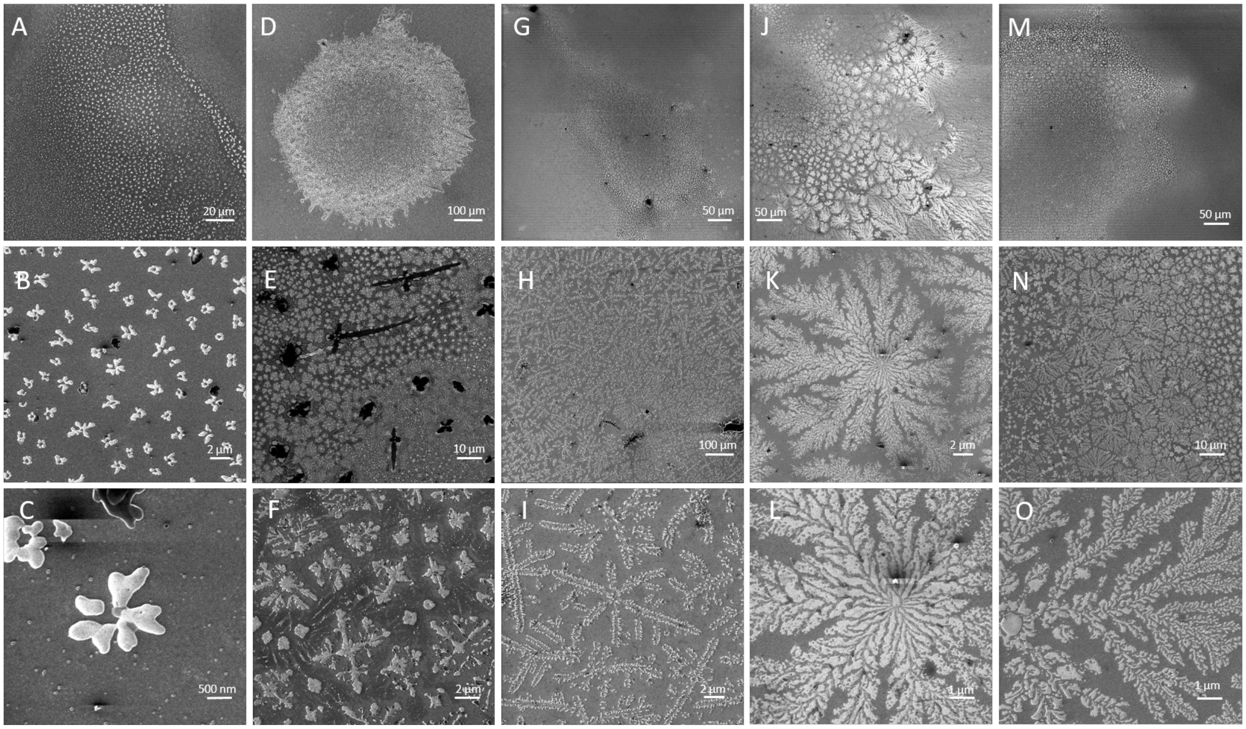


**Fig. S21. Helium ion microscopy (HIM) depict fractal-like assembly with increasing AtzA concentrations.** (**A** to **C**) 0.250 µM AtzAM1 and 2 µM AtzCM1. (**D** to **F**) 0.950 µM AtzAM1 and 2 µM AtzCM1 (**G**-**I**) 1.5 µM AtzAM1 and 2 µM AtzCM1. (**J** to **L**) 3 µM AtzAM1 and 2 µM AtzCM1. (**M** to **O**) 3 µM AtzAM1 and 1 µM AtzCM1.

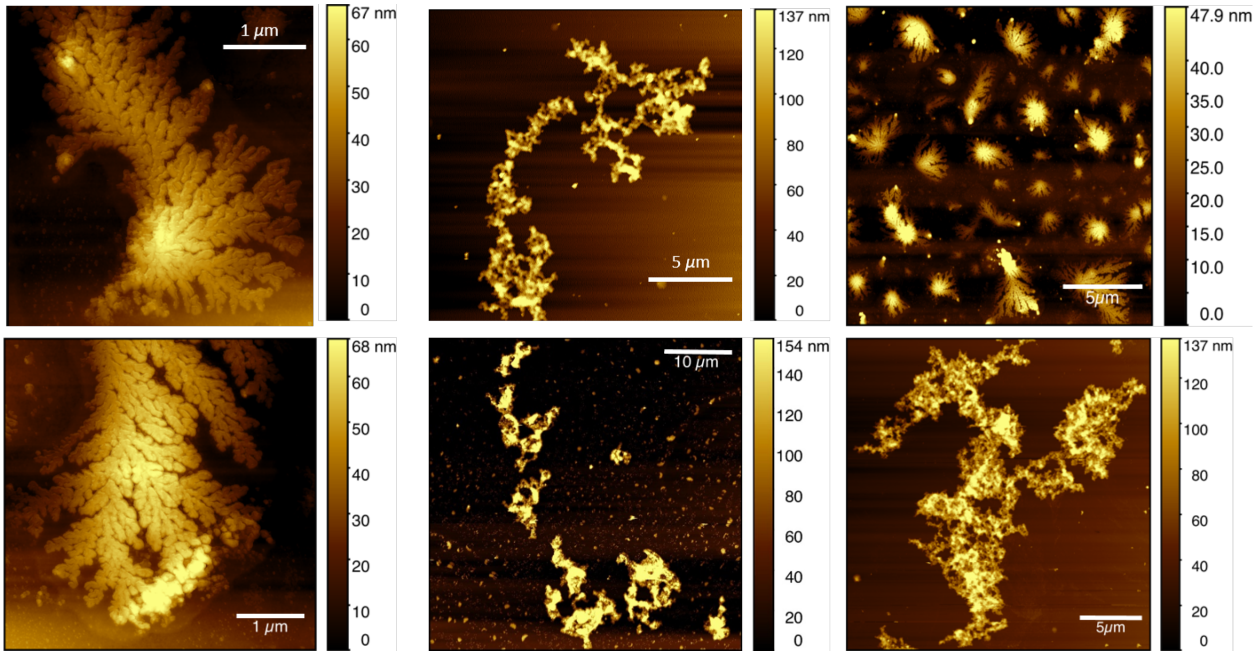


**Fig. S22. Atomic Force Microscopy (AFM) images show fractal-like structures, fern-like, and petal-like structures, similar to HIM.**

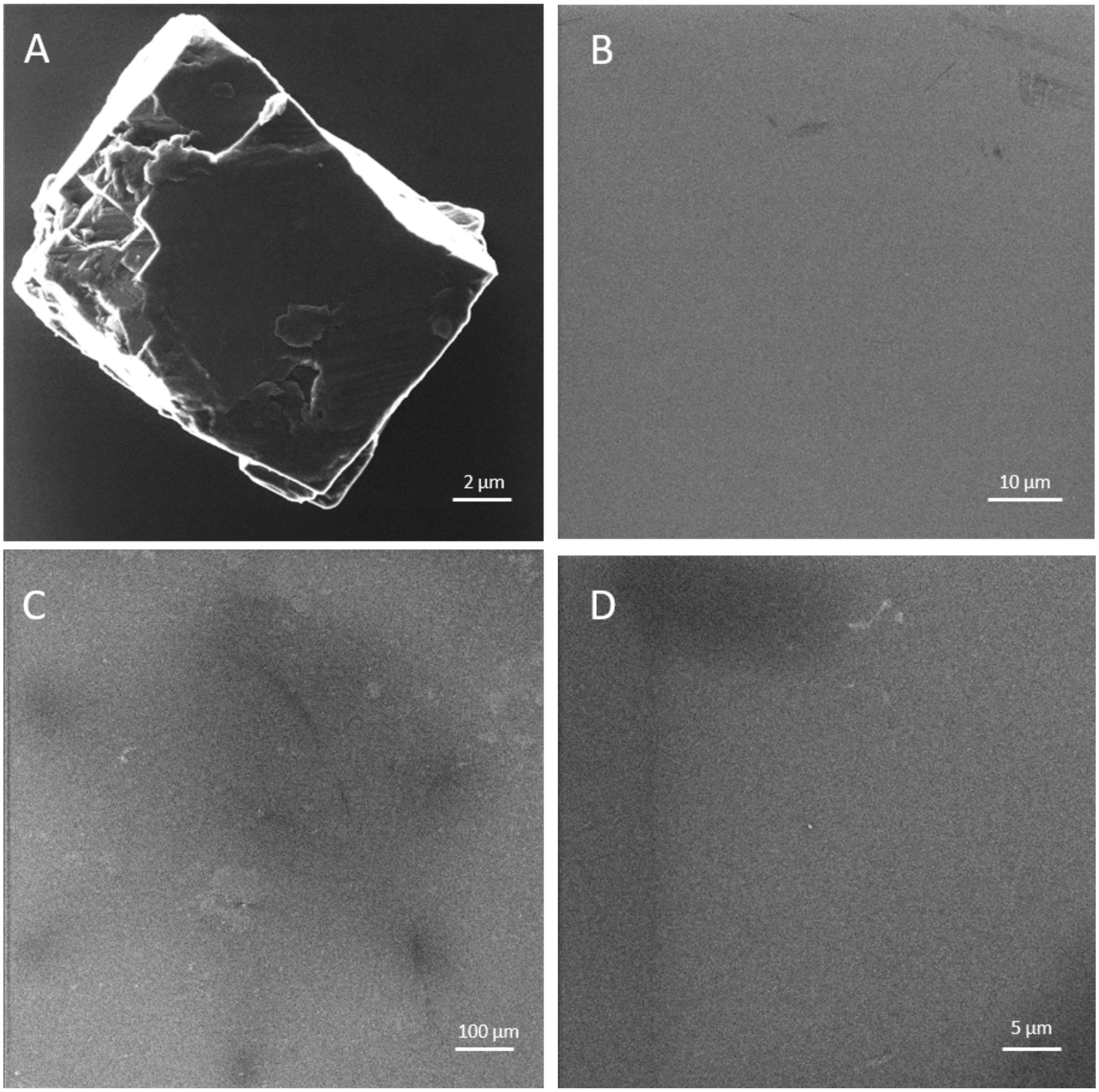


**Fig. S23. Helium ion microscopy (HIM) buffer and non-phosphorylated controls preclude satlt precipitation.** In order to determine that our proteins were forming fractal-like patterns and it was not salt inducing the patterns, a buffer and non-phosphorylated proteins sample controls were used to preclude salt precipitation. (**A**) Usual HIM square salt crystals on a glass surface. (**B**) Deposited HNG buffer (50 mM Hepes, 100 mM NaCl, 5% glycerol, pH.7.4, buffer proteins are stored in) on silicon wafer shows no structures on the surface. (**C**) 3 µM non-pY-AtzAM1 and 2 µM AtzCM1 control shows no fractal-like structures. (**D**) 3 µM non-pY-AtzAM1 and 1 µM AtzCM1 show no fractal-like structures. All controls demonstrate that fractal structures are formed by phosphorylated protein components.

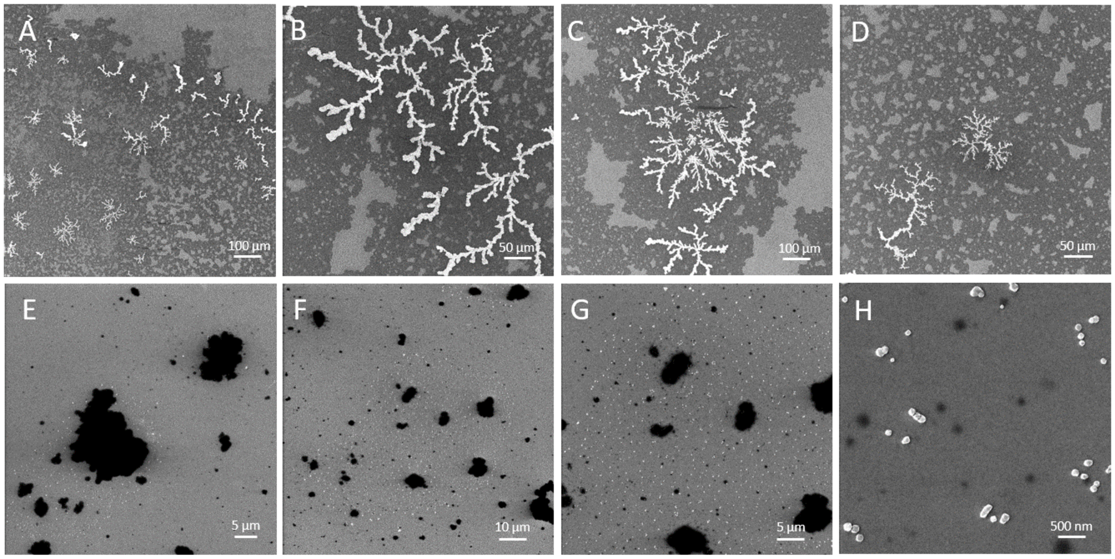


**Fig S24. Helium ion microscopy comparison of fractal assembly and globular assembly.** HIM Images depict fractal-like assembly with 3 uM AtzAM1 and 2 uM AtzCM1 final concentrations (**A** to **D**), while the 3 uM AtzAM1-ExtendedLinker and 2 uM AtzCM1-ExtendedLinker final concentrations show both large and small globular shape proteins on the silicon surface (**E** to **H**).

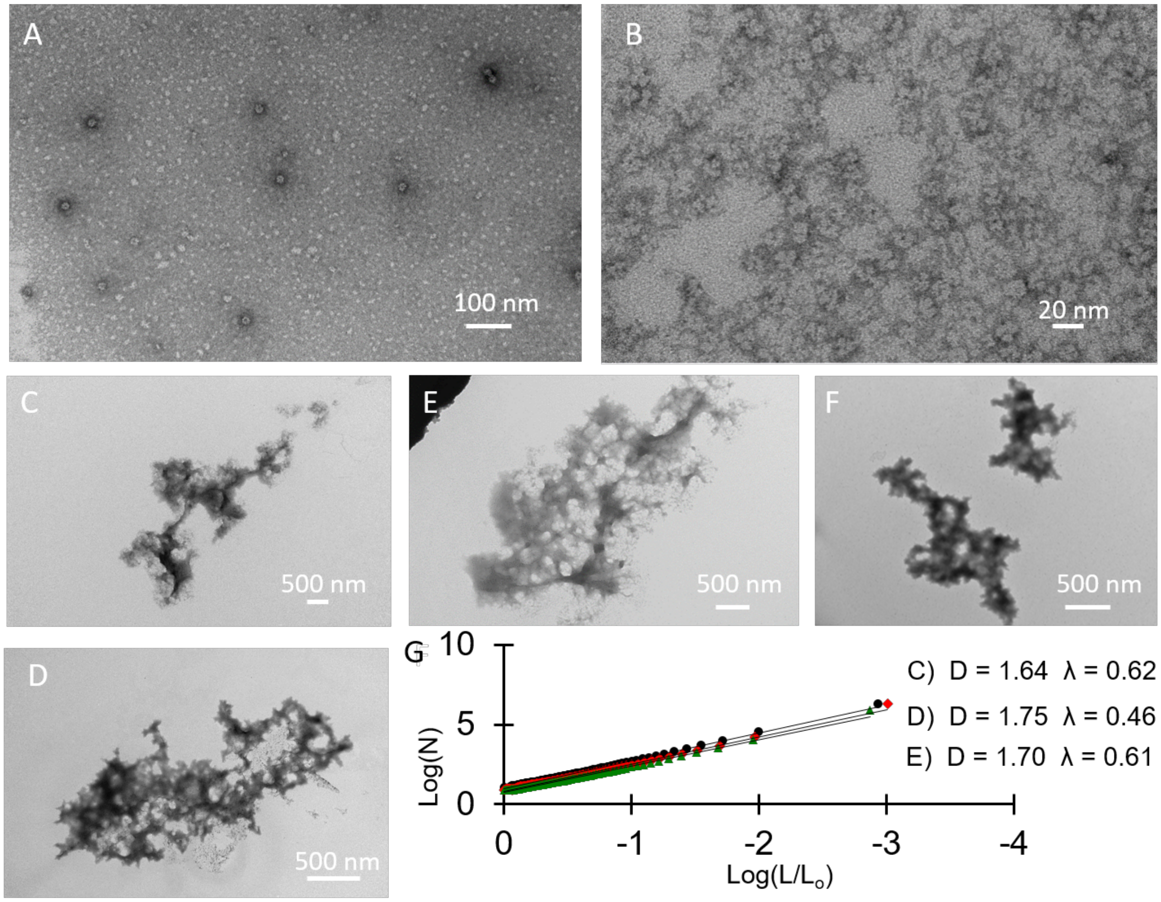


**Fig. S25. Transmission Electron Microscopy (TEM) depicts fractal-like assemblies in the phosphorylated samples while the non-phosphorylated samples depict individual proteins.** (**A** and **B**) ten-fold dilution of 3 μM non-pY-AtzAM1 and 2 μM AtzCM1, which shows the individual proteins. (**C** to **F**) Various assembly images of the ten-fold dilution of 3 μM pY-AtzAM1 and 2 μM AtzCM1 sample which form the fractal-like assembly consistently. (**G**) Image analysis (2D) using box counting yields the expected fractal dimension of ~1.7 for the C, D, and E, TEM images.

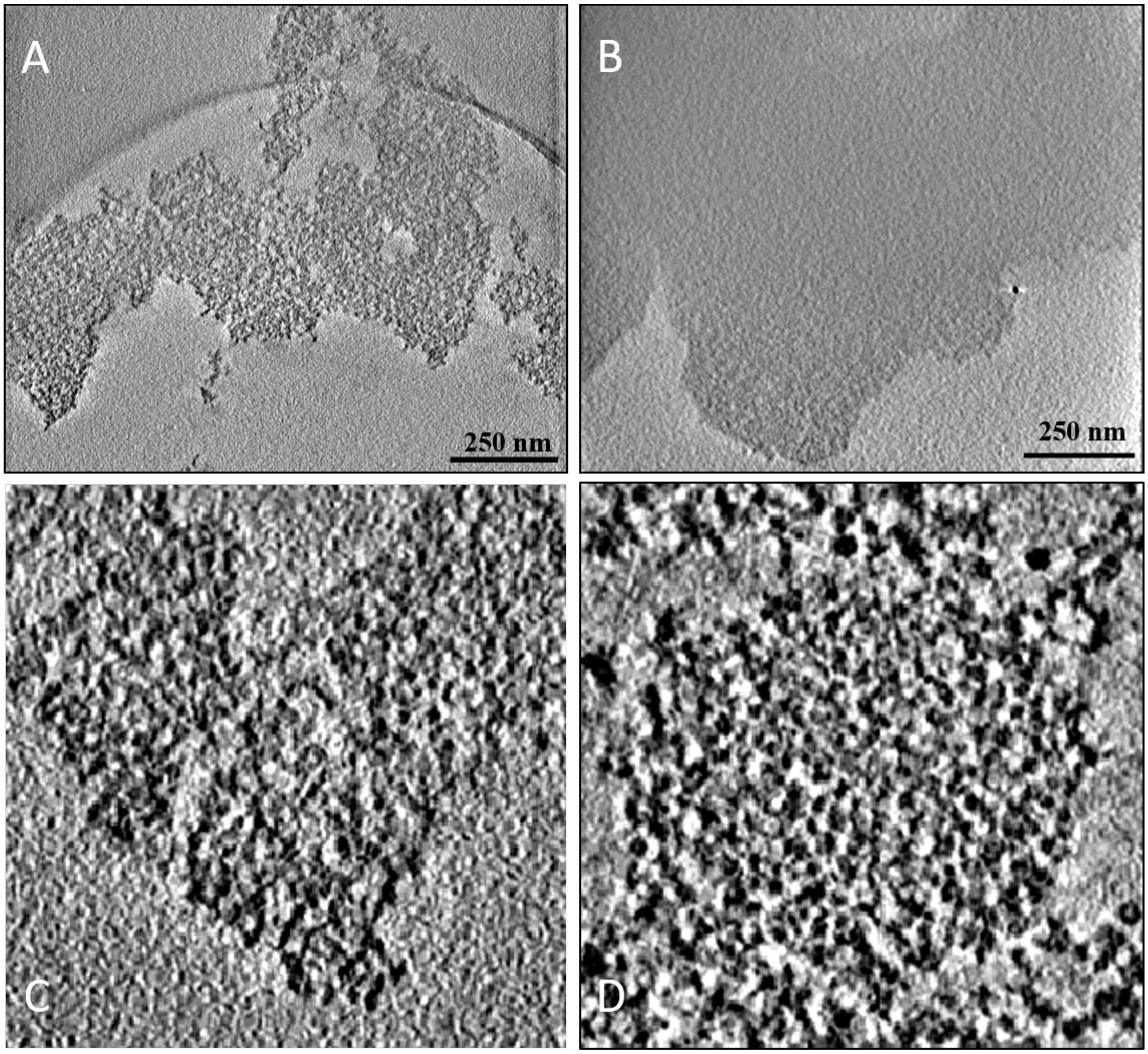


**Fig S26. Comparison of the fractal assembly CryoEM tomograms and the extended linker globular assemblies.** CryoEM tomograms of the fractal-like assemblies (**A**) and the extended linker assemblies (**B**) show a difference in the overall topology of the two different assemblies. Zoomed in versions of the images show representatives of a fractal assembly (**C**) and of a very dense and globular structure (**D**).

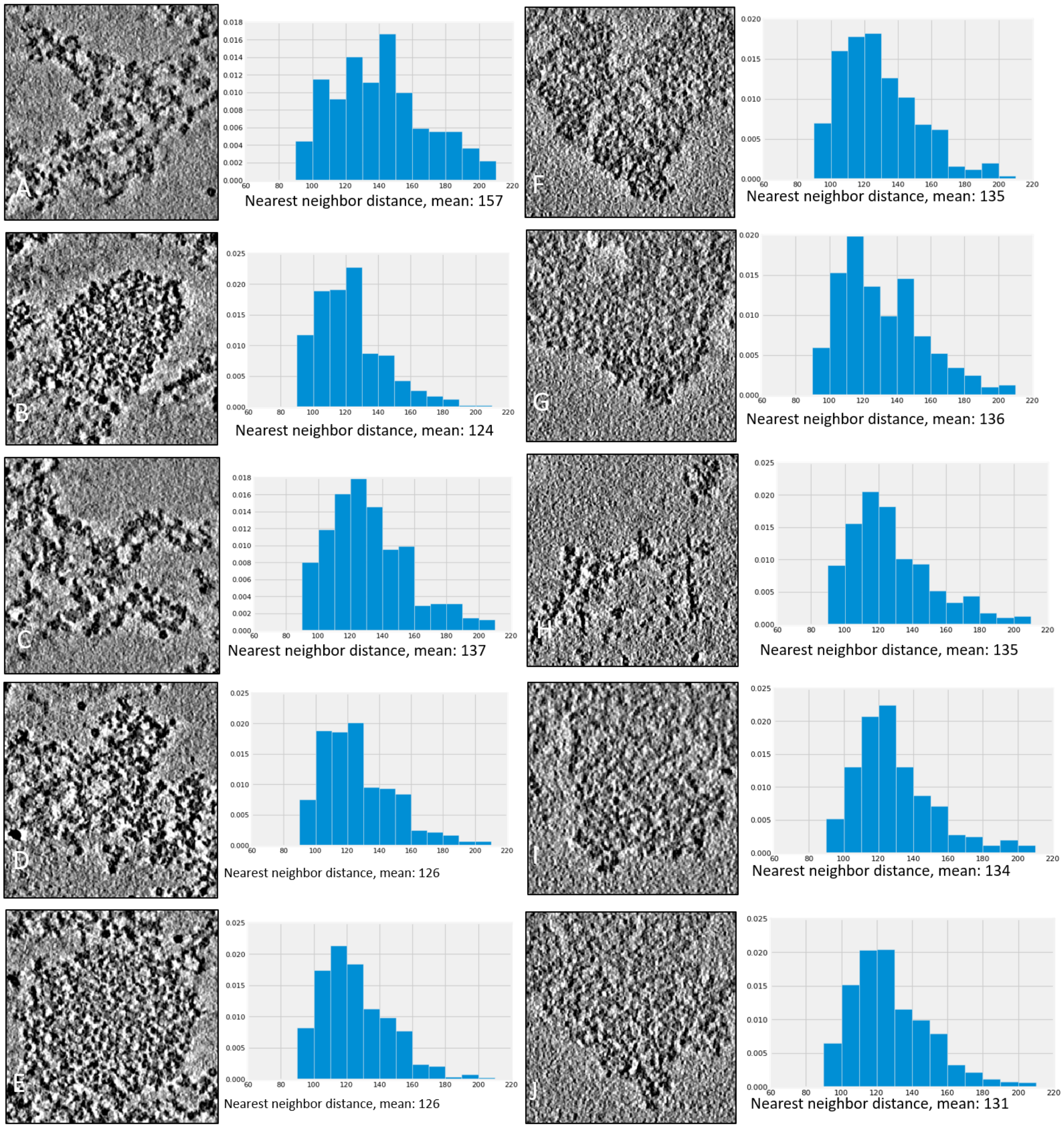


**Fig. S27. Analysis of the fractal assembly CryoEM tomograms and the extended linker globular assemblies.** CryoEM tomograms of the GS-linker assemblies (A-E) and fractal assemblies (F-J) next to the calculated nearest neighbor distance and mean average distance are shown.

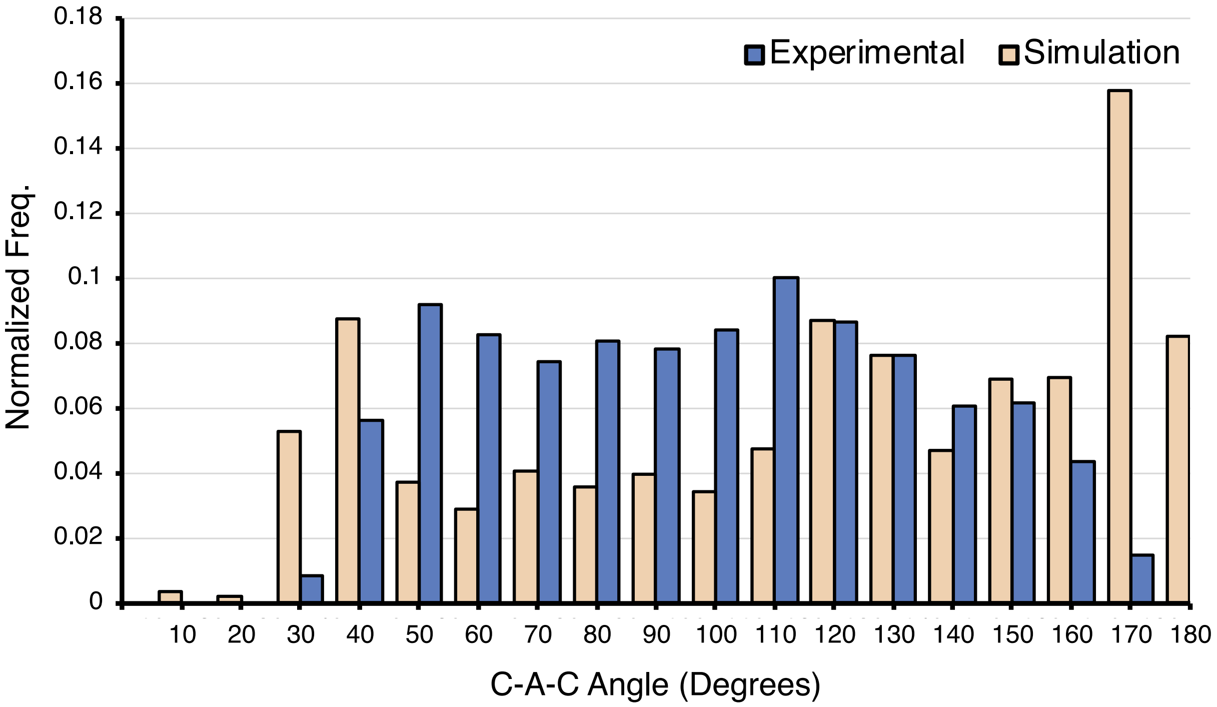


**Fig. S28. Spatial angle comparison of Cryo-EM structure to simulation.** For the large assembly discussed in Figure 4, the angle formed from three proximal components (C-A-C) was calculated using experimentally derived Cryo-EM density fits and the geometric centers of the simulated assembly structures. While both the experimental and simulated assemblies distributions are relatively flat, simulated assemblies show a marked preference for values 30, 170, and 180 compared to the experimental assemblies. The disagreement between the distributions at 170 and 180 may arise due to lack of flexibility in the simulated assemblies which effectively decreases sampling of non-linear connections due to detected steric clashes in a growing assembly. The experimentally derived angular distribution is significantly more sensitive to the assignment of density to individual components compared to inter-component distances. The uncertainty in placement at this resolution (~40Å) may also contribute to the observed differences between experimentally derived and calculated angular distributions more than the errors in distance distributions in Fig.4G)

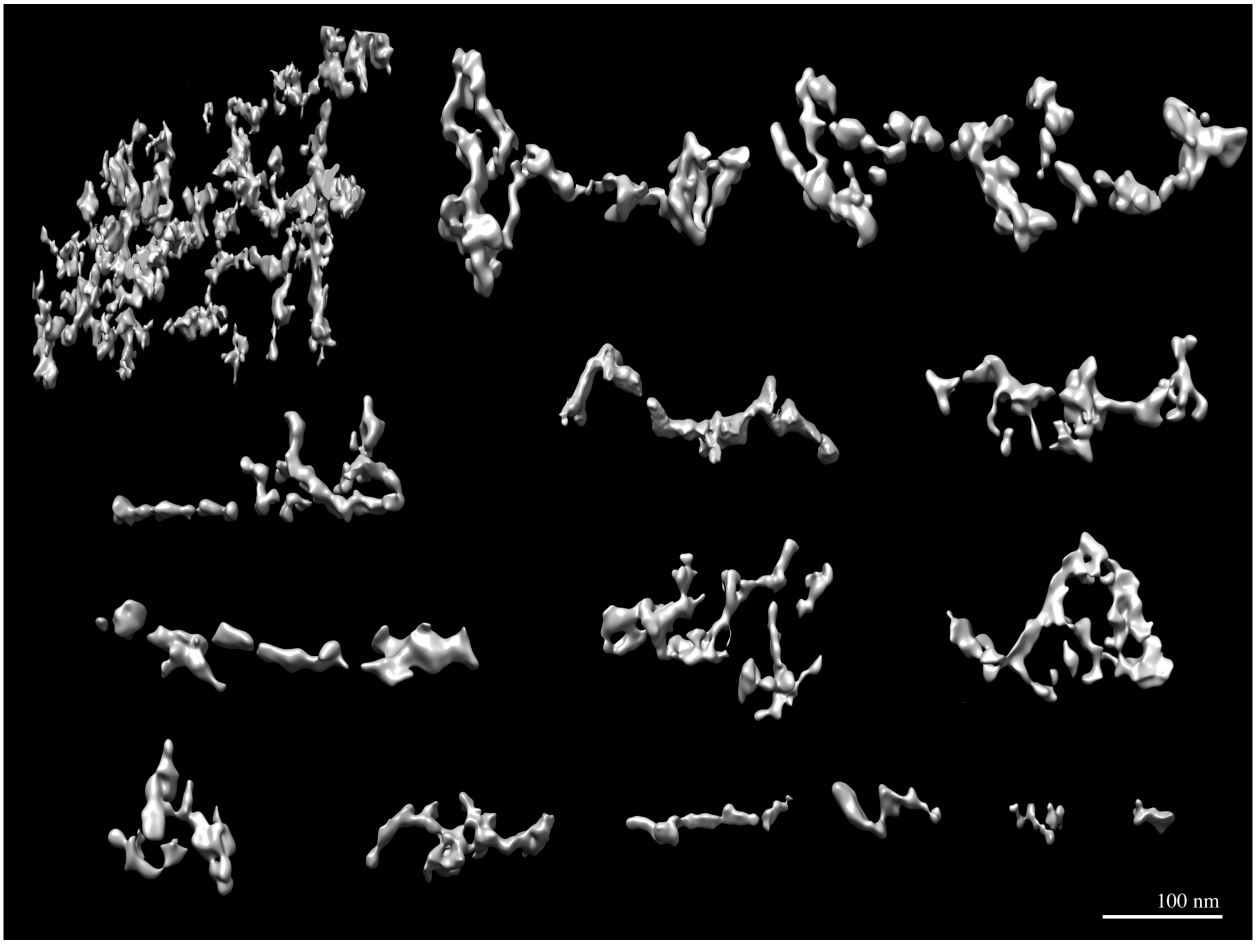


**Fig S29.** Isosurface views of the assembly tomograms, from large to small.

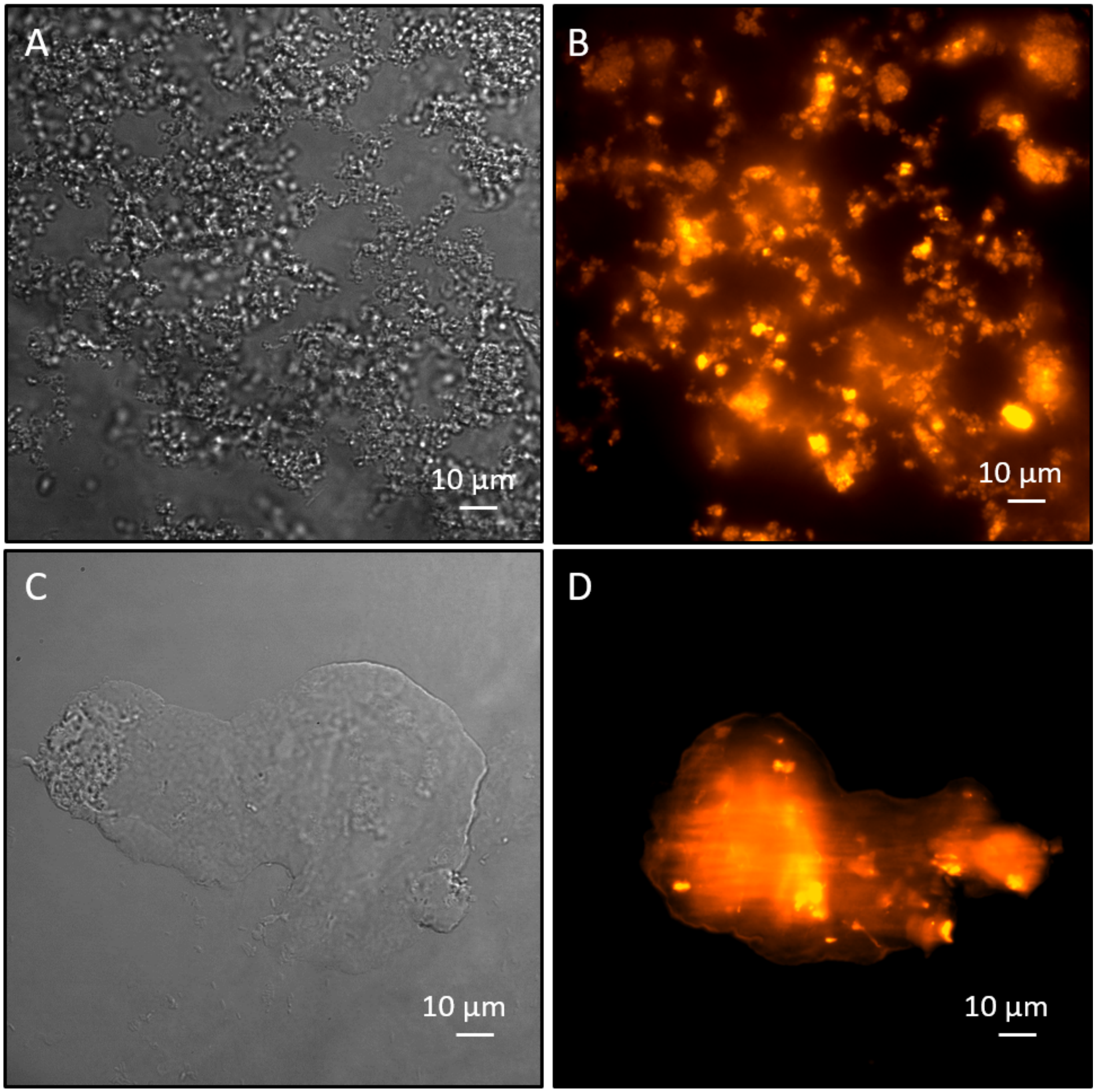


**Fig. S30. Fluorescence microscopy and bright-field images of the 4-component assembly (AtzAM1, AtzCM1, ProteinA-SH2, and antibody, along with extended linker versions of AtzA and AtzC) confirm incorporation of IgG-Antibody-Alexa Fluor 568 into assemblies.** (**A**) Fractal assembly in DIC and (**B**) fluorescent image of fractal indicating incorporation of antibody into assembly. (**C**) Globular assembly in DIC and (**D**) fluorescent images of globular assembly indicating incorporation of antibody into assembly. The depiction of a fractal and globular topology is easily distinguishable in these images.

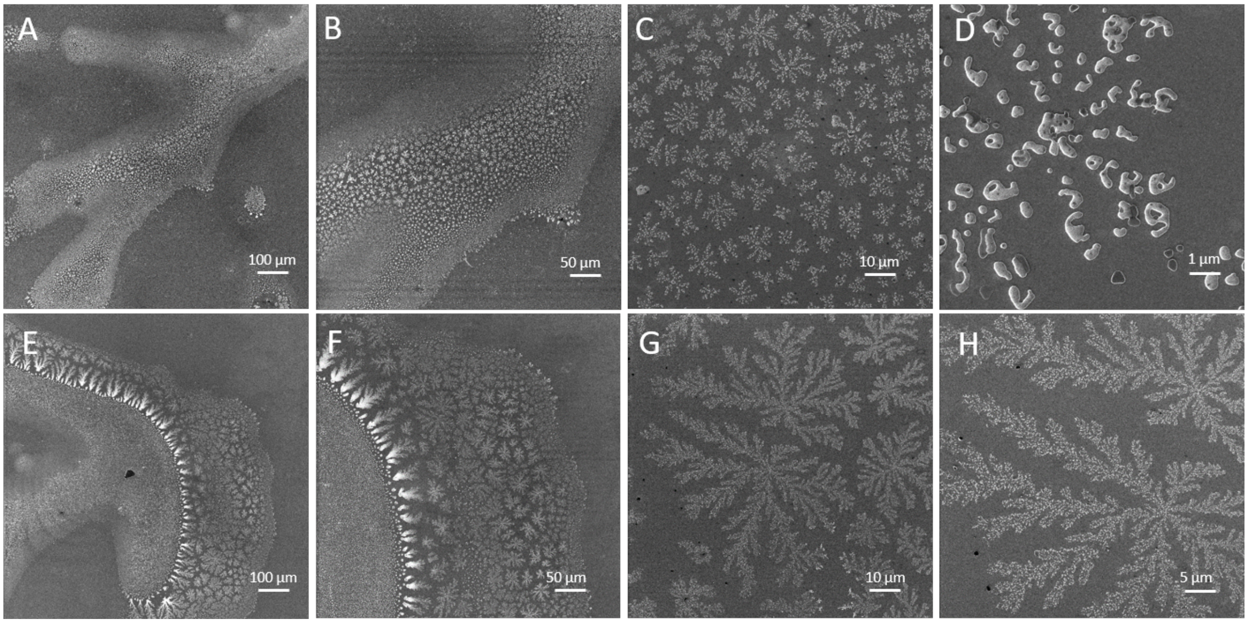


**Fig. S31. Helium ion microscopy (HIM) images depict fractal-like assembly with 3 µM AtzAM1, 1 µMAtzBSH2, 1 µM AtzCM1 final protein concentrations.** (**A** to **D**) Various views of the fractal-like 3-component assembly are shown.

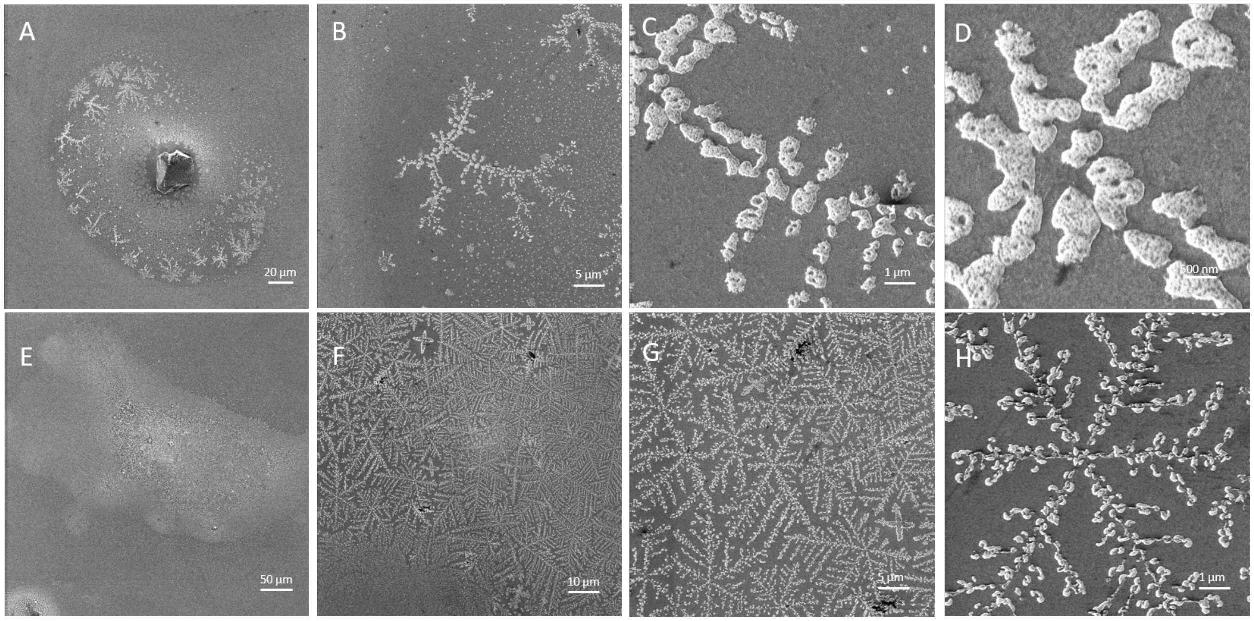


**Fig. S32. Helium ion microscopy (HIM) images depict fractal-like assembly with 3 µM AtzAM1, 1 µMAtzBSH2, 2 µM AtzCM1 final concentrations.** (**A** to **H**) Various views of the 3-component assembly with fractal-like structures are shown.

**Table S3. Comparison of the different AtzA and AtzC ratio components with their fractal dimensions ($D_f$) and λ.**

| Protein Components | $D_f$ ($\mu \pm \sigma$) | λ ($\mu \pm \sigma$) |
|---|---|---|
| A:C (1:8) | 1.60 ± 0.05 | 0.39 ± 0.07 |
| A:C (1:2) | 1.54 ± 0.01 | 0.48 ± 0.03 |
| A:C (3:4) | 1.52 ± 0.08 | 0.51 ± 0.15 |
| A:C (3:2) | 1.66 ± 0.05 | 0.59 ± 0.13 |
| A:C (3:1) | 1.56 ± 0.09 | 0.61 ± 0.21 |
| A:B:C (3:1:2) | 1.49 ± 0.06 | 0.45 ± 0.10 |
| A:B:C (3:1:1) | 1.46 ± 0.10 | 0.39 ± 0.04 |

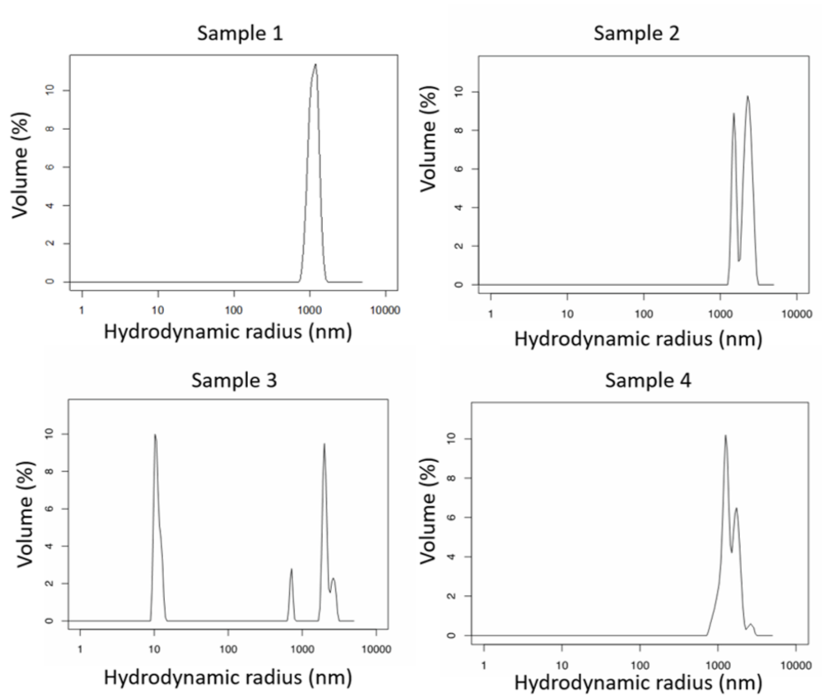


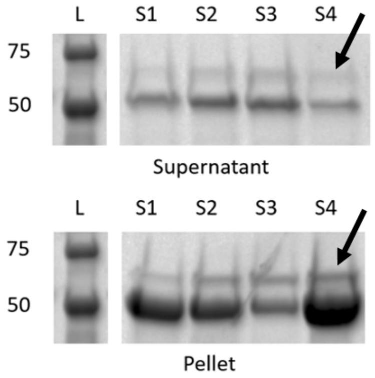


| Sample | AtzAM1 μM | AtzBSH 2μM | AtzCM 1 μM |
|---|---|---|---|
| 1 | 3.0 | 0.5 | 1.5 |
| 2 | 3.0 | 1.0 | 1.0 |
| 3 | 3.0 | 1.5 | 0.5 |
| 4 | 3.0 | 1.0 | 2.0 |

**Fig. S33. DLS and SDS PAGE confirm AtzBSH2 incorporation into the 3-component assembly.** AtzAM1, AtzBSH2, and AtzCM1 were added and allowed to incubate at various concentrations, then analyzed with DLS which showed that the addition of AtzBSH2 continues to have an assembly at ~1 μm. The SDS Page gel samples were pelleted and samples of the three component assembly supernatant and pellet were analyzed. If AtzBSH2 is incorporated into the assembly, it should partition preferentially into the pellet. The gels show that a band at the expected MW weight of AtzBSH2 ~69kda is seen predominantly in the pellet with increasing AtzBSH2 concentrations (Also see Fig S34).

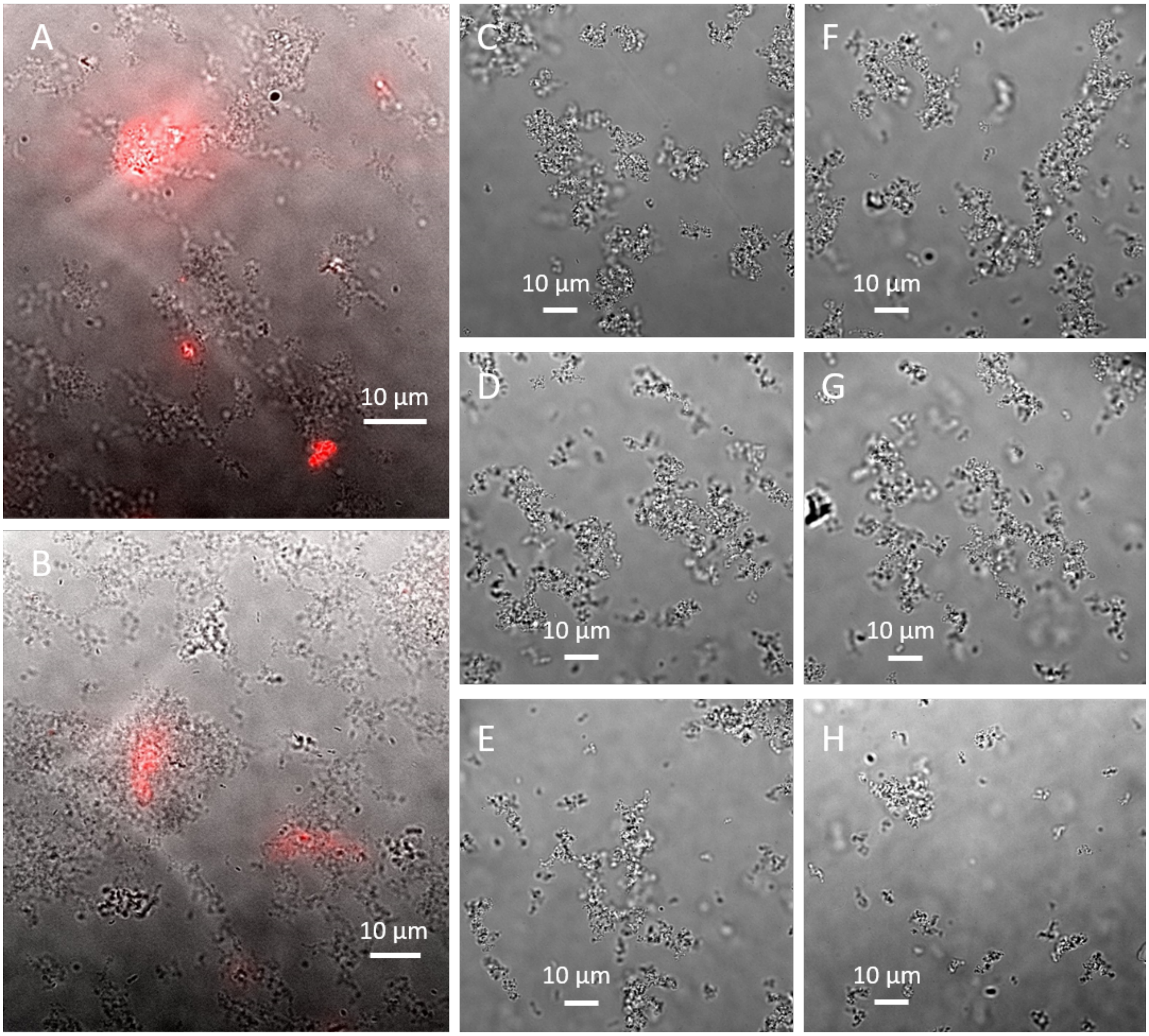


**Fig. S34. Fluorescence microscopy and bright-field images of the 3-component assembly confirm incorporation of AtzBSH2 into assembly while bright-field images confirm the fractal-like nature of the 2-component assembly.** (**A** and **B**) 3 µM AtzAM1, 1 µMAtzBSH2 dye labeled with Alexa Fluor™ 647, 2 µM AtzCM1 image shows AtzBSH2 incorporation into 3-component assembly at various locations (**C** to **H**) 3 µM AtzAM1 and 2 µM AtzCM1 assembly images depict fractal-like assembly structure.

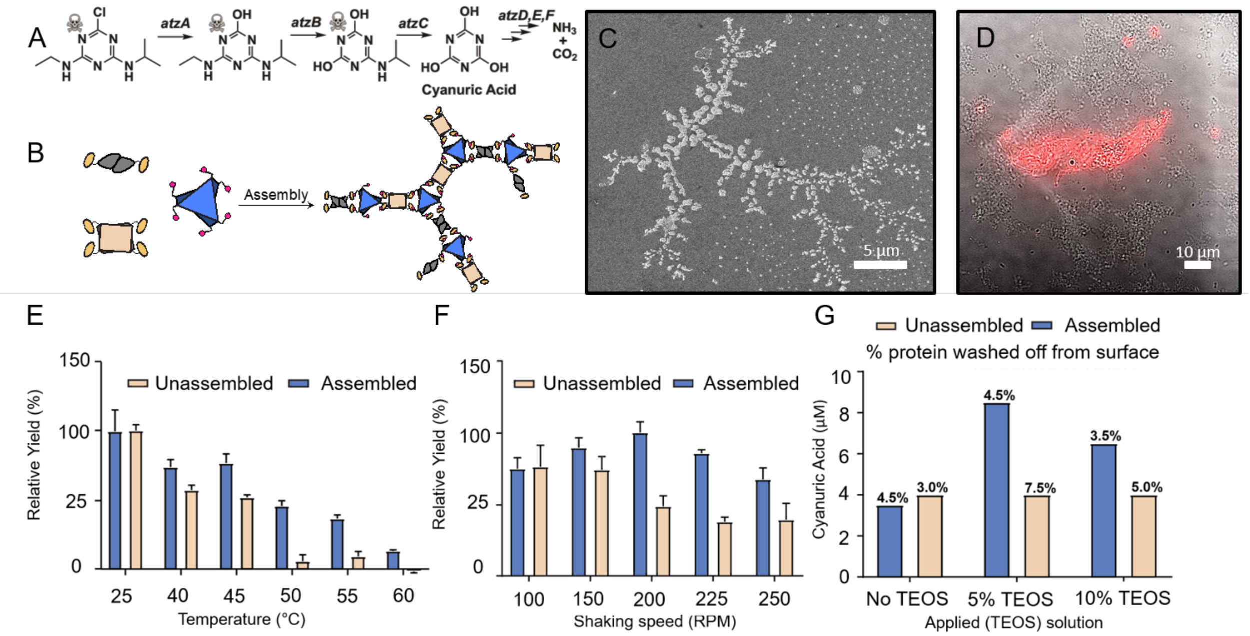


**Fig. S35. AtzBSH2 incorporation to construct a three-enzyme assembly**. (A) Atrazine degradation pathway, enzymatic conversion of atrazine to cyanuric acid, and further enzymatic conversion to NH3 and CO2. (B) AtzB was added as an SH2-domain fusion to the two-component (AtzA-AtzC) assembly. c and d, Three-component assembly formation was validated using HIM (C), and the incorporation of AtzB was confirmed with fluorescence microscopy (using an Alexa-658-labeled AtzB) (D). (E) and (F), Assemblies were found to be more thermotolerant, as detected by incubation at a given temperature for 30 min followed by activity assays, and more robust to mechanical shearing forces, as detected by ability to withstand shaking. (G), Assemblies and free enzymes were incorporated into a Basotect® polymer foam with different TEOS % layers, to trap proteins, and assayed for cyanuric acid production. Proteins can be lost during the wash step after crosslinking and the % of protein lost under each condition is indicated on top of the bars.

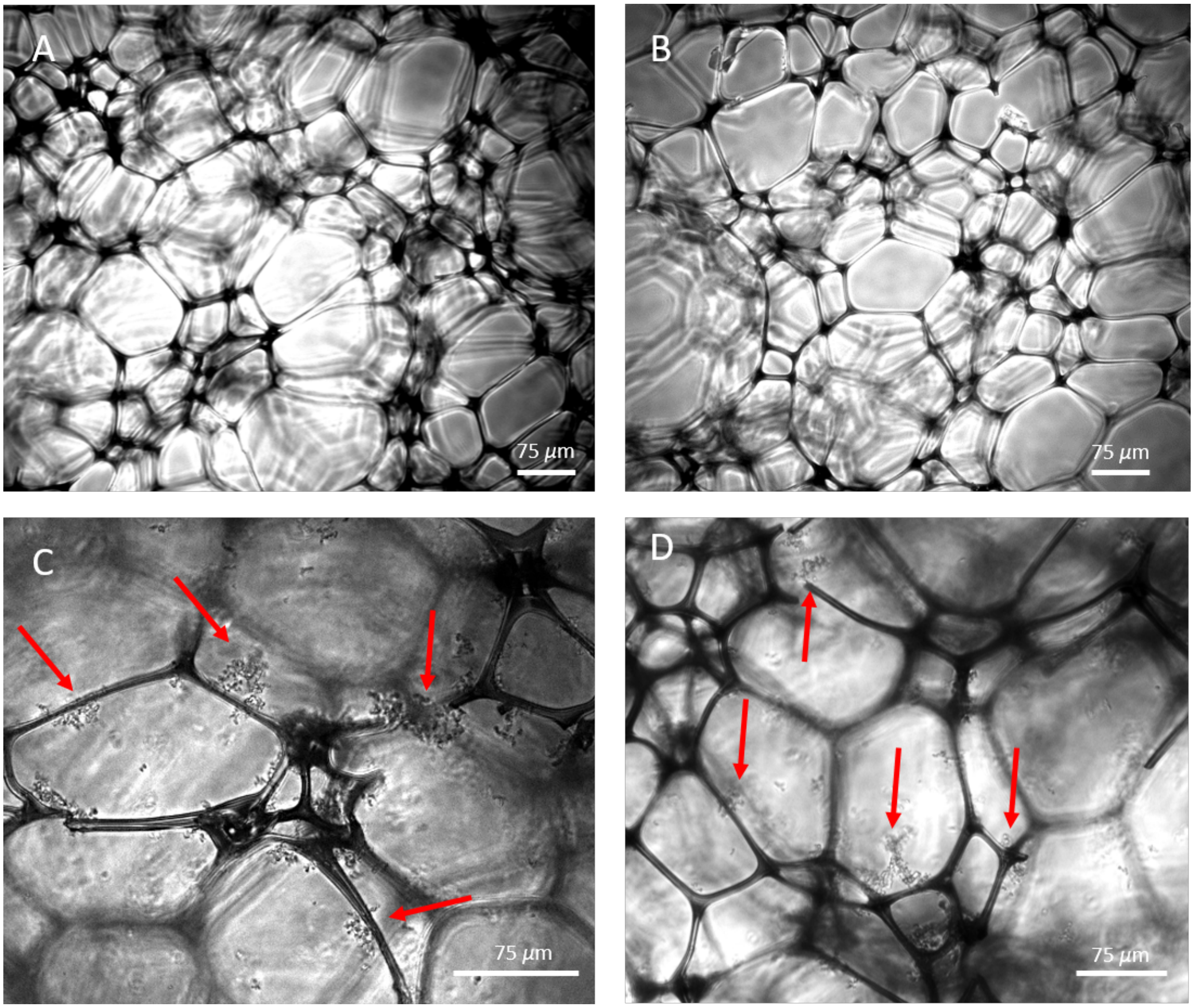


**Fig. S36. Phase contrast micrographs of the Basotect® polymer foam with and without assemblies for the AtzAM1, AtzBSH2, and AtzCM1 components.** (**A** and **B**) The microporous polymer foam with no assemblies. (**C** and **D**) The assemblies have been immobilized into the polymer foam, red arrows depict locations with assemblies. Images were taken with a Leica DM4000 B LED microscope, 10X objective (100X total magnification).

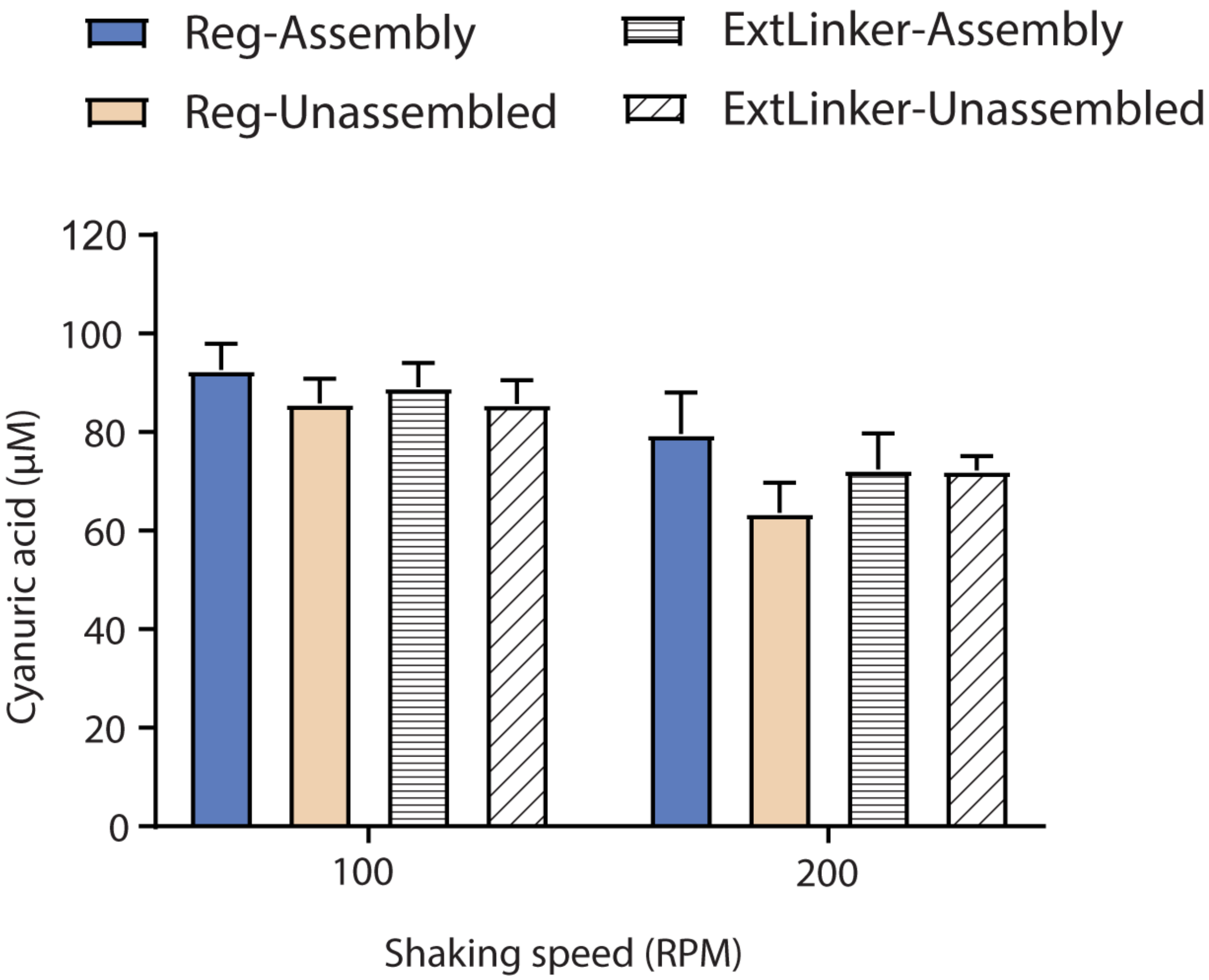


**Fig. S37. The fractal-like assemblies (Reg-Assembly) and the extended linker globular assemblies (ExtLinker-Assembly) enzymatic conversion of atrazine to cyanuric acid demonstrates no enzymatic benefit of a globular assembly.** (**A**) AtzB was incorporated into the two-component assembly as an SH2-domain fusion as previously described to create the three-component assembly for both the fractal and globular assemblies. The activity of the fractal assembly was similar than the extended linker assemblies (GS linker) under different shaking speeds.
